# Genome-Wide Epigenomic Profiling of Primary Non-Small Cell Lung Cancer Reveals Specific and Recurrent DNA Methylation Alterations in Smoker Versus Never-Smoker Patients

**DOI:** 10.1101/2022.10.06.511208

**Authors:** Jennifer A. Karlow, Erica C. Pehrsson, Xiaoyun Xing, Mark Watson, Siddhartha Devarakonda, Ramaswamy Govindan, Ting Wang

## Abstract

Epigenetic alterations are widespread in cancer and can complement genetic alterations to influence cancer progression and treatment. To better understand the potential contribution of DNA methylation alterations to tumor phenotype in non-small cell lung cancer (NSCLC) in both smoker and never-smoker patients, we performed a comprehensive, genome-wide profiling of DNA methylation in 17 primary non-small cell lung cancer and 10 matched normal lung samples using the complementary methylation assays MeDIP-seq and MRE-seq. Compared to patient-matched non-malignant lung tissue, we report recurrent methylation changes of several gene promoters, many previously implicated in cancer, including *FAM83A* and *SEPT9* (hypomethylation), and *PCDH7*, *NKX2-1*, and *SOX17* (hypermethylation). Although smoker and never-smoker patients shared many methylation changes, several were specific and recurrent within a particular smoking status. In particular, never-smokers displayed a greater proportion of hypoDMRs and exhibited a greater number of recurrently hypomethylated promoters, including the promoter of the oncogene *ASPSCR1*, and others previously linked to cancer, including *TOP2A, DPP9,* and *USP39*. Methylation changes outside of promoters were also widespread and often recurrent, particularly the loss of methylation over repetitive elements, highly enriched for ERV1 subfamilies. Recurrent hypoDMRs were also enriched for several transcription factor (TF) binding motifs, often for genes involved in signaling and cell proliferation, including 71% encoding a binding site of *NKX2-1*, which was found to be significantly upregulated in TCGA LUAD samples. Furthermore, the overwhelming majority of DMRs identified in this study were found to reside in an active chromatin state in at least one tissue profiled using the Roadmap Epigenome data, suggesting that methylation changes may contribute to altered regulatory programs through the adaptation of cell type-specific expression programs.

## Introduction

Lung cancer is the leading cause of cancer deaths in the US, with a 5-year relative survival rate of 20.5% [1]. Non-small cell lung cancer (NSCLC) represents 85% of lung cancer cases and is highly heterogeneous, comprising lung adenocarcinoma, squamous cell carcinoma, and large cell carcinoma. In addition to cytotoxic chemotherapies, currently approved therapeutics for NSCLC include inhibitors of EGFR, ALK, ROS1, VEGF, and other proteins, as well as checkpoint blockade therapies targeting PD-1 and PD-L1 [2]. Lung adenocarcinoma and squamous cell carcinoma both exhibit high somatic mutation rates relative to other cancers [3].

Although most NSCLC cases are associated with tobacco smoking [4], the proportion in never-smokers is rising and now represents 10-40% of cases worldwide [5]. Lung cancer in never-smokers is highest among women and East Asians, is enriched for the lung adenocarcinoma subtype, and is linked to genetic susceptibilities and certain environmental exposures [4]. Never-smoker tumors exhibit distinct molecular characteristics, including a higher frequency of *EGFR* and *HER2* mutations and *ALK*/*RET*/*ROS* fusions that improve their response to targeted therapy. In contrast, smokers exhibit a higher rate of mutations in *KRAS*, *TP53*, *STK11*, *BRAF*, *JAK2*, *JAK3*, and mismatch repair genes [5, 6]. The mutation burden in smokers is >10X higher [5] and is characterized by C>A nucleotide transversions caused by direct benzo[*a*]pyrene exposure. This signature is clonal, suggesting that mutations occur before transformation, and correlates with pack years smoked [7]. Smokers also exhibit a higher rate of copy-number alterations, non-synonymous mutations, and neoepitopes [7, 8], which may explain their greater response to immunotherapy.

In the past, cancer analysis has focused primarily on genomic mutations. However, cancer is also characterized by massive epigenetic dysregulation. Epigenetic modifiers are frequently mutated in cancer, suggesting a role in tumorigenesis. Cancer exhibits global DNA hypomethylation coupled with focal promoter hypermethylation [9], which can silence tumor suppressor genes in lung cancer [6]. In addition, epigenetic alterations can lead to genomic instability, the dysregulation of genomic architecture [10], histone modification spreading, and widespread DNA hypomethylation resulting in aberrantly activated transposable elements. Cryptic promoters within de-repressed transposable elements can drive oncogene expression [11–13] and create chimeric transcripts that may produce neoepitopes [14, 15]. Additionally, demethylated endogenous retroviruses produce double-stranded RNA and trigger an anti-viral immune response [16, 17], potentiating treatment with checkpoint blockade therapy [18]. In sum, epigenetic alterations can complement genomic alterations and have profound effects on cancer progression and treatment.

The Cancer Genome Atlas (TCGA) has profiled thousands of NSCLC samples and matched normal lung [19, 20]. However, DNA methylation data was generated using the Human Methylation 450K array, which covers only a fraction of the CpGs in the genome. To supplement this analysis, we profiled genome-wide DNA methylation in 17 primary lung adenocarcinoma tumors, as well as matched normal lung from 10 of the patients (see Materials and methods). Clinicopathologic data is summarized in Supplementary Table 1. For each sample, we performed methylated DNA immunoprecipitation sequencing (MeDIP-seq) and methylation sensitive restriction enzyme sequencing (MRE-seq), which capture methylated and unmethylated CpGs, respectively (Figure S1). Data generated by these complementary assays was then integrated using methylCRF [21], estimating the methylation level at over 28 million CpGs genome-wide, resulting in comprehensive, environmentally matched pairs of normal lung and primary tumor methylomes. We also used the M&M algorithm, which integrates MeDIP-seq and MRE-seq from two comparative samples [22] to identify differentially methylated regions, providing a complete profile of common DNA methylation alterations across a heterogeneous set of primary NSCLC tumors.

## Results

### Global methylation changes

We first profiled genome-wide DNA methylation changes between primary NSCLC and patient-matched, histologically non-malignant lung tissue. Although there was substantial variation across samples (Figure S2a), on average, primary tumor samples exhibited a shift in overall methylation density, with fewer highly- and lowly-methylated CpGs and a significant increase in intermediately methylated CpGs (**Figure 1**a, Wilcox *p*- value = 0.001423; Supplementary Text; Figure S2b). The genome-wide methylation changes observed in this study are in line with previously published results [23, 24], and analysis of MRE-seq data alone confirmed that tumors lose methylation over intergenic regions and repeats (Figure S3a-h; Supplementary text), consistent with previous studies [25].

**Figure 1.**
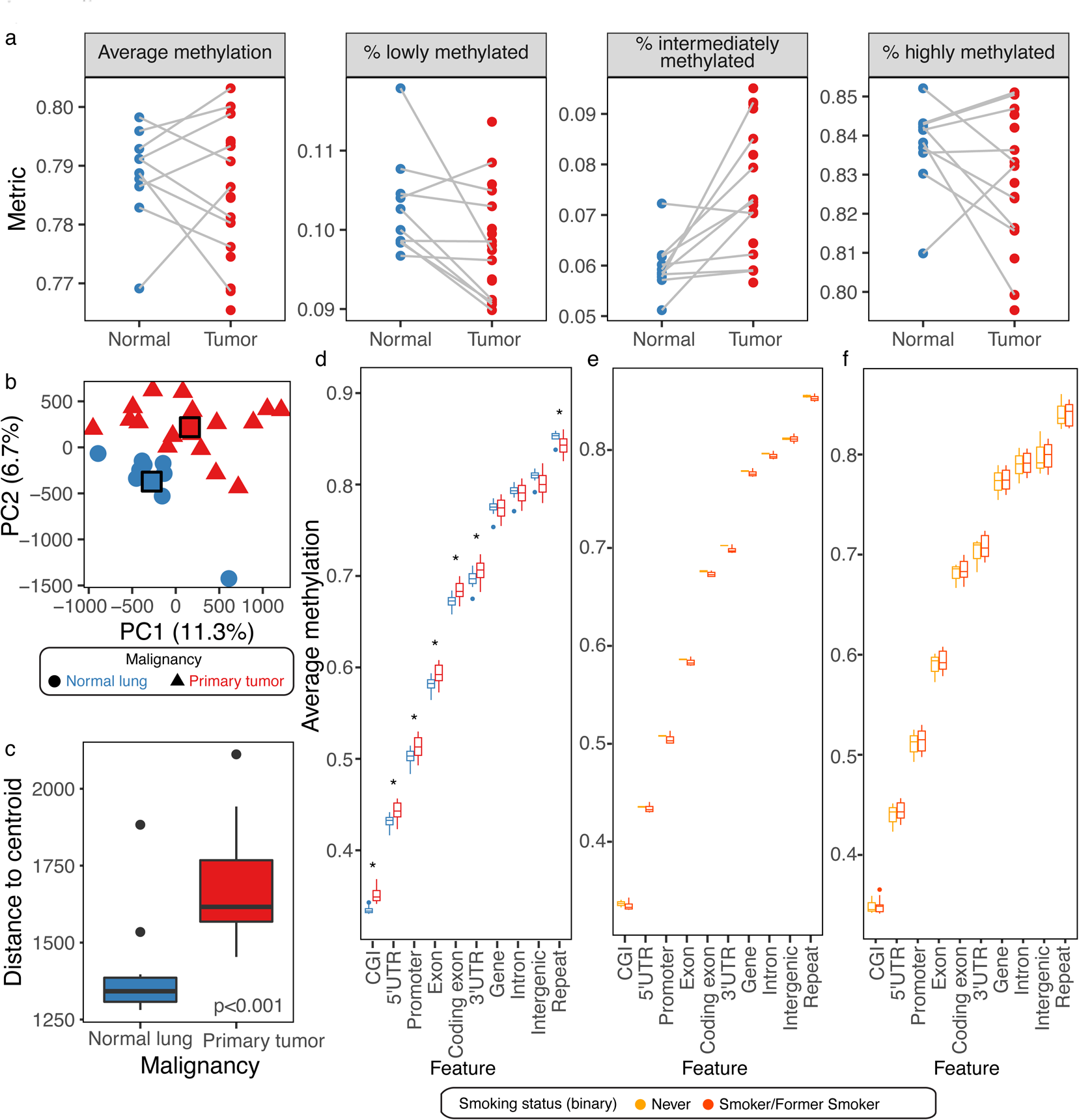
Genome-wide CpG methylation alterations in primary NSCLC. **A.** Average genome-wide methylation and proportion of CpGs at each methylation level per sample (lowly methylated, < 30% methylation; intermediately methylated, 30 – 70% methylation; highly methylated, > 70% methylation). **B.** PCA on normal lung and primary tumor samples, using the mean methylation over genome-wide 1 kb windows as features. Group centroids are indicated with squares (color legend to the right of Figure 1e). Axis titles display the amount of variance explained by each PC. P2385_N_UC is the outlier on PC2. **C.** Distance to centroid by sample malignancy (Wilcox test *p*-value = 0.0004497). Normal outliers are P4999_N_S and P2385_N_UC; tumor outlier is P14658_T_NS. **D.** Average methylation level over all CpGs overlapping each feature in each sample, colored by sample malignancy and ordered by median methylation level. Stars indicate Wilcox *p*-value < 0.05. **E.** Average methylation level over all CpGs overlapping each feature in normal samples, separated by smoking status. **F.** Average methylation level over all CpGs overlapping each feature in tumor samples, separated by smoking status.

At a finer resolution, local methylation changes separated normal lung from tumor samples. Principal components analysis on the average methylation level over 1 kb windows separated normal lung from primary NSCLC along PC1 and PC2 (Figure 1b). The degree of correlation between tumor and paired normal was not different when separating patients based on smoking status (Figure S4a-c, Wilcox *p*-value = 1) or tumor stage (Figure S4d-g, early vs. late Wilcox *p*-value = 1) and was not correlated with tumor purity (Figure S4h-j, Pearson’s correlation *p*-value = 0.6079). The two patients with adenosquamous carcinoma demonstrated a greater divergence from their paired normal samples than patients with adenocarcinomas, although this also did not meet statistical significance (Wilcox *p*-value = 0.09524) (Figure S4k-m). Interestingly, interpatient normal-to-primary tumor distances were comparable to intrapatient distances (Figure S4n-q, Wilcox *p*-value = 0.06136), suggesting that tumors undergo a great deal of epigenetic repatterning relative to non-malignant tissue.

Non-malignant lung tissue was equally homogenous among smokers and never-smokers (Wilcox *p*-value = 0.10052, distance to centroid; Figure S5a-c), and was significantly more homogenous than primary NSCLC (*p* < 0.001, Wilcox test, distance to centroid; Figure 1c), suggesting that epigenetic changes during tumor progression can take a number of paths. Primary tumors were also equally homogenous among smokers and never-smokers (Wilcox *p*-value = 0.8363, distance to centroid; Figure S5d-f), across tumor stages (early- vs. late-stage Wilcox *p*-value = 0.8981, distance to centroid; Figure S5g-i), and across subtypes (adenocarcinoma vs. adenosquamous carcinoma Wilcox *p*-value = 0.7676, distance to centroid; Figure S5j-l). No correlation was found between the degree of variation of tumor samples and their tumor purity (Pearson product moment correlation *p*-value = 0.7664; Figure S5m-o).

Finally, we looked at methylation alterations in primary NSCLC by genomic location (Figure 1d). Compared to normal lung, primary NSCLC gained CpG methylation over promoters, exons (both UTRs and coding exons), and CpG islands but lost methylation over repeats (Wilcox *p* < 0.05), in agreement with previous studies [25]. Interestingly, the two normal lung samples from never-smokers exhibited higher methylation across each profiled genomic feature compared to normal lung from smokers, although small sample sizes prohibit significance (Figure 1e). On the other hand, tumors separated by smoking status did not show any differences in methylation across genomic features (Figure 1f).

### Differentially methylated regions

Next, we identified differentially methylated regions (DMRs) across the genome between patient-matched normal lung and primary NSCLC, as well as between normal lung samples and between normal lung and tumor samples from different patients. A *q*-value threshold of 0.001 was selected (Figure S6a-g; Supplementary Text).

The number of DMRs per patient was highly variable, ranging from 239 to 16,676 (**Figure 2**a), independent of tumor purity (Figure S7a). However, all but one patient-matched comparison exhibited more DMRs than were found between pairs of normal lung samples (59 to 1,708 DMRs in normal vs. normal comparisons; Figure S7b). The tumor in question (Patient 4999) was a bronchioloalveolar carcinoma, a minimally invasive subtype of NSCLC that may have accumulated fewer methylation changes due to a slower rate of cell division [26]. Based on genome-wide methylation profiles, normal lung samples were more homogenous than primary NSCLC (Figure S7c-f; Supplementary Text). The proportion of DMRs that were hypomethylated in normal lung compared to tumor also varied by patient, from 2 to 73% (Figure 2a), and did not correlate with the total number of DMRs (Pearson correlation p > 0.5).

**Figure 2.**
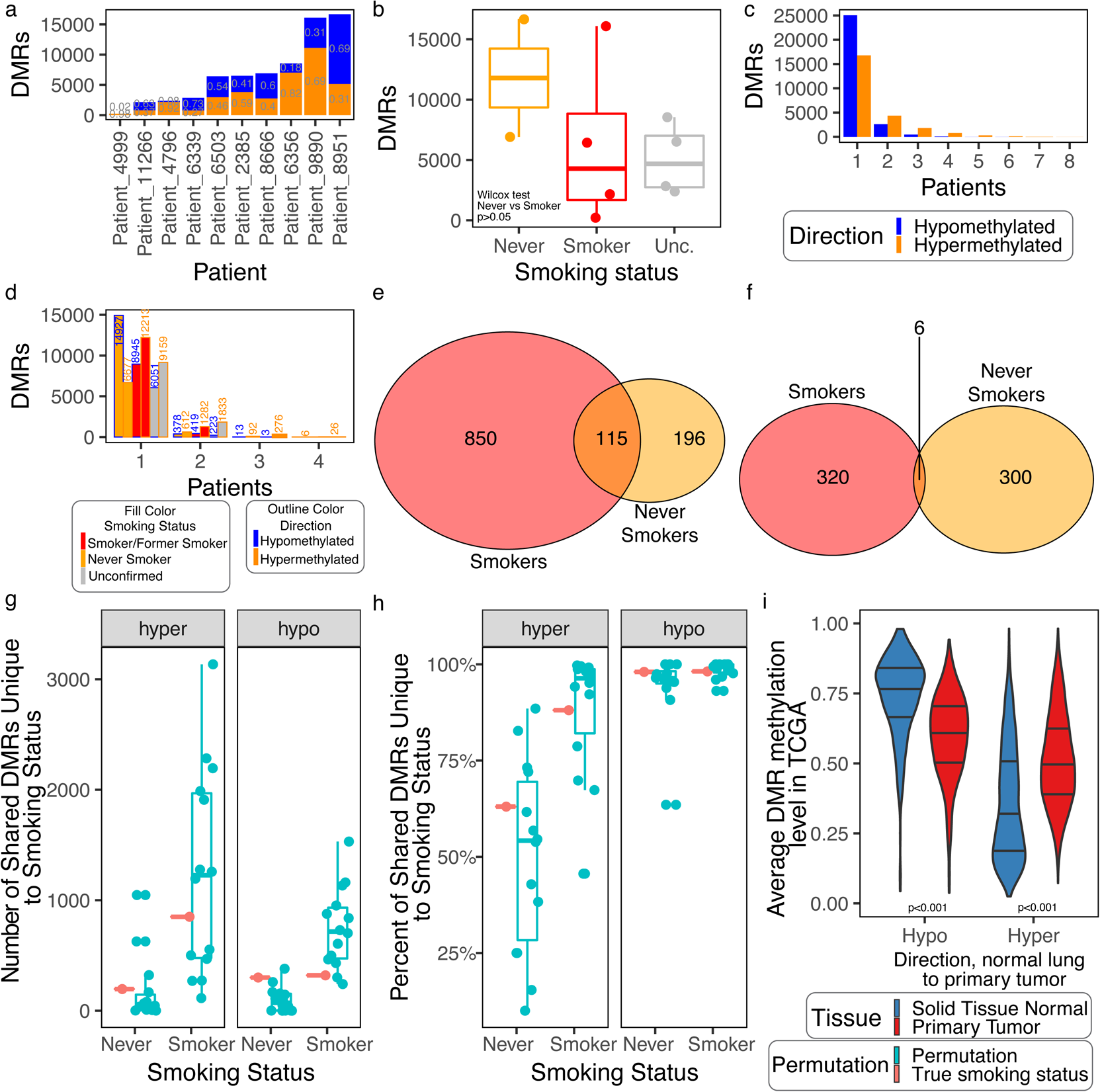
DMRs between normal lung and primary NSCLC. **A.** Number of DMRs per patient, including the proportion by DMR direction. Proportions < 0.1 are not shown. **B.** Number of DMRs per patient, according to smoking status. Wilcox test comparing never-smokers (*n* = 2) to smokers (*n* = 4) was not significant (*p* = 0.2667). **C.** Number of DMRs shared by each number of patients, by direction. **D.** Number of DMRs shared by each number of patients, by direction and smoking status. **E.** hyperDMRs recurrent in smokers and never-smokers or both. **F.** hypoDMRs recurrent in smokers and never-smokers or both. **G-H.** Number **(G)** and percentage **(H)** of recurrent hyper- (left) or hypoDMRs (right) unique to the specified smoking status. Red indicates the values using true patient smoking statuses; blue indicates values using permutations of patients (smoker *n* = 4, never-smoker *n* = 2). **I.** Mean methylation level of each hypo- or hypermethylated DMR shared by at least two patients across all TCGA LUAD samples, split by sample type. Violin plot lines indicate quartiles. Wilcox *p* < 0.001 between LUAD normal lung and primary tumor for both hypo- and hypermethylated DMRs. Unc. = unconfirmed.

To determine whether smoking created DNA methylation field effects prior to malignant transformation, we identified DMRs between normal lung samples based on smoking category (*n* = 1,214 unique DMRs). Normal samples of the same smoking status did not have fewer DMRs, as would be expected if there were characteristic changes in CpG methylation due to smoking (Wilcox *p* > 0.05; Figure S8). Furthermore, DMRs found between never-smoker and smoker normal samples, but not between the two never-smoker normal samples (*n* = 785), were not significantly enriched in any GO Biological Processes. However, there were several examples of methylation loss in only smoker samples, including DMRs in the promoters of *PM20D1* (chr1:205818500-205819000) and *ARRB2* (chr17:4612500-4613000). *ARRB2* variants have been previously linked to smoking status [27], and its depletion promotes lung cancer growth in a mouse model [28]. Additionally, there were DMRs overlapping the gene bodies of *METAP1D*, *RGPD8*, and *PKP3*, which have been shown to be upregulated in lung adenocarcinoma and promote cancer growth [29], and whose methylation status has been linked to *in utero* nicotine exposure [30]. The number of DMRs identified in non-malignant lung tissue specifically between smokers and never smokers was not significantly different from the number of DMRs identified among the entire cohort, suggesting that unlike somatic mutations, smoking history may not contribute to a systematic difference in methylation status.

While underpowered to reach significance (Wilcox test *p*-value = 0.2667), there were fewer DMRs between paired normal and tumor samples from smokers (*n* = 4 patients) than from never-smokers (*n* = 2 patients), with the exception of Patient 9890 (Figure 2b). While never-smoker patients tended to have higher numbers of both hyper- and hypo-DMRs, they tended to have a higher percentage of hypoDMRs relative to smokers (Figure S9a-c, Wilcox *p*-values > 0.05). The number of DMRs as well as percentage hypomethylated did not correlate with tumor stage (Figure S9d-e), subtype (Figure S9f-g), or total methylation change (Figure S9h-i).

The majority of patient-matched DMRs were exclusive to a single patient (Figure 2c), with 11% of hypomethylated DMRs (*n* = 3,158 of 28,204 DMRs) and 31% of hypermethylated DMRs (*n* = 7,457 of 24,242 DMRs) shared by multiple individuals. However, hypomethylated DMRs were shared by up to six patients, and hypermethylated DMRs were shared by up to eight. When considering patients according to smoking status, 8% (*n* = 612 of 7289) hyper- and 2% (*n* = 378 of 15,305) hypoDMRs were shared by both never-smokers, and 10% (1,380 of 13,593) hyper- and 5% (432 of 9,377) hypoDMRs were shared by at least two smokers (Figure 2d). Of the DMRs identified in more than one patient (recurrent), 850 and 320 hyper- and hypoDMRs, respectively, were shared by at least two smokers and absent in never-smokers, 196 and 300 shared by both never-smokers and absent in smokers, and 115 and 6 recurrent in both smokers and never-smokers (Figure 2e, Figure 2f). Interestingly, the number of recurrent never-smoker DMRs not recurrent in smokers was generally greater than when randomly subsetting patients, suggesting that these recurrent epigenetic changes may contribute to never-smoker cancer progression (Figure 2g, Figure 2h).

Finally, we confirmed the DMRs observed in this study using a much larger cohort from TCGA. Although only 14% of patient-matched hypoDMRs and 52% of hyperDMRs overlapped a CpG from the 450K array (*n* = 3,815 and 12,668, respectively), the average methylation level of the DMRs in TCGA lung adenocarcinoma (LUAD) and matched normal lung samples recapitulated the DMR directions in our study (Figure 2i, Figure S10a). Furthermore, hyperDMRs identified in smoker patients from our study (at all or recurrently) exhibited significantly higher methylation levels in TCGA patients with confirmed smoking history compared to never-smokers/patients without data (Figure S10b, Figure S10d). HypoDMRs found in smoker patients from our study exhibited lower methylation levels in TCGA LUAD samples with confirmed smoking history (Figure S10c) but failed to exhibit a significant difference compared to TCGA LUAD never-smokers/patients without data when restricted to recurrent smoker hypoDMRs (Figure S10e). Differences in average methylation between TCGA LUAD tumor samples from patients with confirmed smoking history and never-smokers/patients without data were not observed over hyper- or hypoDMRs identified in never smokers from our study (Figure S10f-S10i), with the exception of significantly lower methylation in TCGA patients with confirmed smoking status over hypoDMRs identified in non-smokers from our study (Figure S10g).

### Genomic location of DMRs

The locations of DMRs between patient-matched primary and non-tumor samples recapitulated recognized patterns. In general, hypermethylated loci in tumors (analyzing 500 bp bins containing CpGs) were enriched in promoters, exons, CpG islands, and partially methylated domains (PMDs), and were depleted over intergenic regions and repeats, whereas hypoDMRs were enriched to a lesser extent in promoters, exons, CpG islands, and PMDs (**Figure 3**a, Figure S11a). Stratifying DMRs based on identification in smokers, never-smokers, or both revealed similar trends. Enrichment was slightly higher over promoters, exons, and CpG islands for hyperDMRs identified in smokers than in never-smokers, whereas hypoDMRs identified in never-smokers demonstrated a slightly higher enrichment over these regions (Figure S11b, Figure S11c). Enrichment over genic features was stronger for DMRs shared between patients than for patient-exclusive DMRs, where the pattern was again more pronounced for hyperDMRs (Figure S11d).

**Figure 3.**
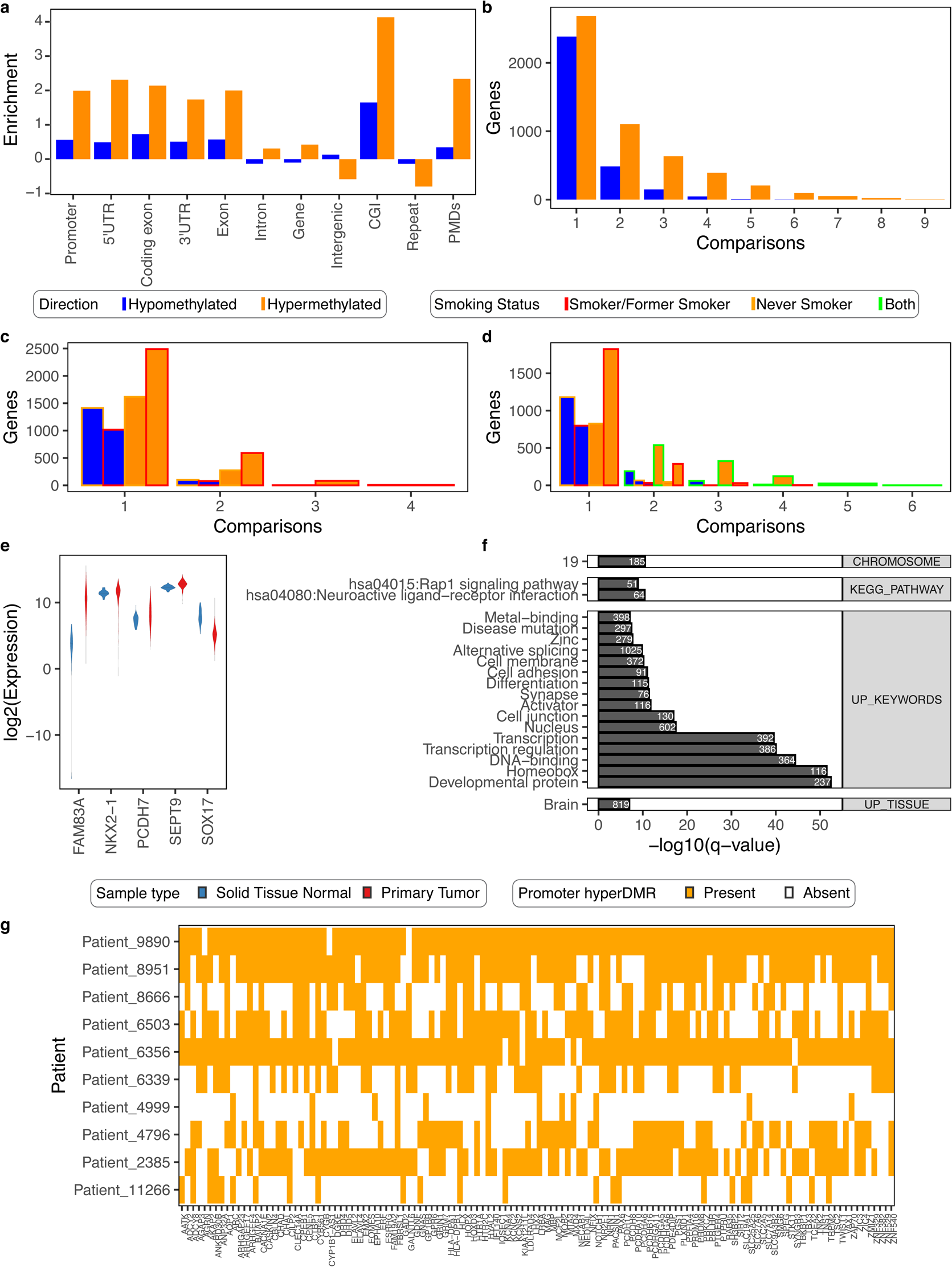
DMRs within gene promoters. **A.** Log odds ratio enrichment of DMRs over genic features, intergenic regions, CpG islands (CGI), and repeats compared to the background distribution of 500 bp bins containing CpGs, by DMR direction. **B-D.** Number of normal vs. patient-matched tumor comparisons in which each gene contains a DMR in its promoter(s), **(B)** by DMR direction, **(C)** by DMR direction and smoking status, **(D)** and by DMR direction and unique smoking status (classifying DMRs as identified in smokers, never smokers, or both), **E.** Expression of select genes in TCGA LUAD samples (red) and matched normal lung (blue). A pseudocount of 0.00001 was added to each value. Lines represent median values. **F.** Significantly enriched gene sets among genes with a hypermethylated DMR in the promoter in at least two patient comparisons, as determined by DAVID (top 20 terms by corrected *p*-value, out of 160 terms). Only terms with a Benjamini-corrected *p*-value < 0.05 are included. Terms are labelled with the number of selected genes and ordered by corrected *p*-value within each category. **G.** Indication of DAVID Brain UP_TISSUE Pathway genes with a hyperDMR in the promoter in at least 5 patients, stratified by patient.

The enrichment of DMRs over genic features also varied by transcript type. Figure S11e displays DMR enrichment over protein-coding, antisense, lincRNA, and processed transcripts, which together comprise 72% of all GENCODE-annotated transcripts. HyperDMRs were more strongly enriched over the promoters and exons of protein-coding and antisense transcripts, with a 5’ versus 3’ UTR bias evident only in protein-coding transcripts. In contrast, hypoDMRs were most strongly enriched over lincRNA, although still to a lesser degree than hyperDMRs. Stratifying DMRs based on identification in smokers, never-smokers, or both revealed similar enrichment trends (Figure S11f). Finally, hyperDMRs were more strongly enriched over protein-coding transcripts with a CpG island in the promoter, particularly over the promoter (Figure S11g), where smoker hyperDMRs showed a slightly higher enrichment in CpG island-associated promoter transcripts, and never-smoker hyperDMRs showed a slightly higher enrichment in non-CpG island-associated transcripts (Figure S11h).

Both recurrent never-smoker- (*n* = 2) and smoker- (*n* > 1) specific hyperDMRs were enriched in functions related to transcription and embryogenesis (Figure S12a, Figure S12b), but smoker-specific hyperDMRs were also enriched in terms related to cell signaling, epithelial tissue, and the respiratory system. Several genes, including *NOTCH1*, *STK11*, *AKT1*, *CDKN2B*, *DLX5*, *EYA1*, *GATA3*, and *SKI*, were represented by many of the terms. In contrast, recurrent never-smoker- and smoker-specific hypoDMRs did not display significant gene ontology enrichment.

89% of CpG islands overlapped a hyperDMR, and 39% overlapped a hypoDMR, which was reduced to 78% and 15%, respectively, when only DMRs shared between patients were considered. 12% of genes (*n* = 5,193) contained a hyperDMR in the promoter, and 7% (*n* = 3,075) contained a hypoDMR, although only 5% (*n* = 2,171) and 1% (*n* = 465), respectively, overlapped shared DMRs. The results by gene biotype are presented in Supplementary Table 2. Genes with a DMR in the promoter included 48 genes previously shown to be altered (amplified, mutated, methylated, fused, etc.) in NSCLC (27 of which contained a DMR in more than one patient (recurrent)), 188 genes from the COSMIC Cancer Gene Census database (including 39 oncogenes, 25 TSGs, and 15 genes annotated as both) (104 recurrent), 164 epigenetic regulators from the Epifactors database (78 recurrent), 43 CTAs (25 recurrent), and 29 of the top 100 most highly expressed genes in lung (16 recurrent).

Although most genes with a promoter methylation change were unique to an individual, genes were found to overlap a hypoDMR in up to 6 patients and a hyperDMR in up to 9 (Figure 3b). When considering smoking status, both smokers (*n* = 4) and never-smokers (*n* = 2) displayed recurrent hyperDMRs (679 and 273, respectively) and recurrent hypoDMRs (79 and 99, respectively) (Figure 3c). Interestingly, despite having only two never-smokers, the number of recurrent hypoDMRs was greater than the number recurrent in the four smokers, suggesting perhaps that the never-smoker hypoDMRs might be more relevant to tumor formation. Classifying genes with altered promoter methylation according to whether the alteration was seen in smokers, never-smokers, or both, revealed that the majority of recurrently altered genes appear in both smokers and never smokers (Figure 3d). However, 65/99 genes with recurrent hypoDMRs and 46/273 genes with recurrent hyperDMRs in never-smokers are not seen in smokers, and 34/79 genes with recurrent hypoDMRs and 318/679 genes with recurrent hyperDMRs in smokers are not in never-smokers. Genes with recurrent never-smoker-specific hypomethylated DMRs in the promoter include *ASPSCR1*, a known fusion oncogene whose methylation is linked to prenatal smoking and reduced lung function [31]; the DNA topoisomerase *TOP2A*, a pan-cancer upregulated gene [32, 33]; and *DPP9* and *USP39*, which have been implicated in lung cancer [34, 35]. Genes with recurrent never-smoker-specific hypermethylated DMRs in the promoter include *SMYD2*, a lysine methyltransferase that targets EML4-ALK fusion proteins [36], and *CYP1B1*. CYP1B1 converts benzo(a)pyrene from tobacco smoke into its more carcinogenic form and is upregulated in smokers compared to never-smokers [37]. In many cases, the DMR was one of many in the gene promoter, although no transcript contained both a never-smoker and a smoker-specific DMR.

Genes associated with recurrent promoter hypomethylation, regardless of smoking status, include *FAM83A*, a lung cancer biomarker and potential oncogene [38], whose expression in TCGA LUAD samples was almost 100-fold over that found in non-malignant lung samples (Wilcox *p* < 0.001; Figure 3e). The *FAM83A* promoter was hypomethylated in six patients, and several isoforms also lost methylation in TCGA (Wilcox *p* < 0.001). *SEPT9* exhibited promoter hypomethylation in 5 patients and hypermethylation in 3 patients. Although overall gene expression increased significantly in the TCGA LUAD samples (Figure 3e), individual transcripts could be up- or down-regulated (Figure S13). Other genes containing recurrently hypomethylated promoters included *TNFRSF10A* (5 patients), a TRAIL receptor that induces apoptosis [39], and *MET* (3 patients), a known oncogene implicated in NSCLC. In addition, *MUC5B*, which is involved in goblet cell mucus production, is downregulated in mucinous adenocarcinoma [40], and was previously shown to be associated with LUAD cancer-specific DNA methylation changes [41], displays promoter hypomethylation in 4 patients in our study.

Genes with promoter hypermethylation in multiple patients included those within the HOXA cluster, which is densely hypermethylated in NSCLC (see below) [42, 43]. Additionally, the promoter of *PCDH7*, a protocadherin gene with an oncogenic function in lung cancer [44], was hypermethylated in 9 patients. Its expression level significantly increased in LUAD samples (Wilcox *p* < 0.001; Figure 3e), although like many genes, its expression had a much larger range than in normal lung. The promoter of *NKX2-1*, a lung cancer biomarker involved in lung development [45], was hypermethylated in 5 patients (Figure S14), as was that of *SOX17*. Several genes with recurrent promoter hypermethylation and/or nearby intergenic hyperDMRs were also previously shown to be associated with LUAD or LUSC cancer-specific hypermethylation changes, including *HYAL2* (8 patients), *AQP1* (7 patients), *XRCC3* (5 patients), *RARA* (4 patients), *SPTBN1* (3 patients), *EPAS1* (3 patients), *CD34* (2 patients), and *CLU* (2 patients). Other notable genes included *SYT10* and *KCNC1* (8 patients each).

In many cases, the DMR was over the promoter of a shorter or non-coding isoform of the gene. However, we also identified potential instances of promoter switching, in which one transcript promoter of a gene became hypermethylated while another became hypomethylated (Figure S15a-f). For both *ADCY2* and *ASPG*, the promoter of the longest protein-coding transcript (*ADCY2*: ENST00000338316.4, *ASPG*: ENST00000455920.2, ENST00000551177.1, ENST00000551177.1) became hypermethylated in five patients, and the promoters of shorter isoforms became hypomethylated in a subset. The promoter methylation level of the longest transcripts also increased significantly in TCGA LUAD samples (Figure S15a/d), and the transcript expression levels dropped (Figure S15c/f; Wilcox *p*-value < 0.001). In both cases, the overall gene expression level dropped as well (Figure S15b/e; Wilcox *p*-value < 0.001); however, because the shorter isoforms were not captured by TCGA, we were unable to determine whether their expression increased.

In addition to individual genes with highly recurrent DMRs in the promoter, we looked at pathway-level effects using DAVID. Genes with hypoDMRs in their promoters in multiple patients were enriched in spleen-specific expression (*q*-value < 0.05; Figure S15g, Figure S15h), while genes overlapping hyperDMRs were enriched for brain-specific expression (Figure 3f, Figure 3g). Both sets were enriched on chromosome 19 and in alternative splicing functions. Those overlapping hyperDMRs were also enriched in several functions, including cell adhesion and terms related to homeobox genes and development, reflecting the abundance of hyperDMRs over HOX clusters.

We also determined whether DMRs overlapping gene promoters encoded binding motifs for transcription factors dysregulated in NSCLC, which could provide a coordinated method for regulating several genes at once. Based on HOMER known motif analysis, shared hypomethylated promoter DMRs were enriched in binding motifs for transcription factors involved in signaling and cell proliferation (Figure S15i). Additionally, 71% of the DMRs encoded a binding site for Nkx2-1 (up from the background observed value of 64%). In our dataset and in TCGA, promoters of the gene were recurrently hypermethylated, but its expression was not significantly different between normal lung and primary tumors. Hypermethylated promoter DMRs were enriched for transcription factors involved in embryonic development, as well as hypoxia and angiogenesis (Figure S15j).

DMRs in intronic or intergenic regions may also serve as alternative promoters or enhancers that become activated in NSCLC. In contrast to promoters, they are also less likely to have been captured by TCGA or other large studies that relied on probe-based methylation profiling methods. In this study, we identified several recurrent intergenic DMRs, including chr13:53775000-53775500, a bivalent enhancer that was hypermethylated in 8 of the 10 patients and is 49 kb from the nearest gene; and chr6:158182500-158183000, which was hypomethylated in 6 of the 10 patients and is 61 kb from the nearest gene. There were also recurrent intronic DMRs, including the hyperDMR chr14:37136000-37136500, a bivalent enhancer within a *PAX9* intron (8/10 patients), and the hypoDMR chr2:240169000-240169500 within an *HDAC4* intron (6/10 patients).

Like hypoDMRs overlapping promoters, shared intergenic DMRs were also enriched for AP-1 complex binding motifs, as well as other cell signaling and proliferation pathways (Figure S16a-b). Several of the transcription factors for which shared intergenic hypoDMRs have enriched binding motifs have altered expression levels in TCGA LUAD data, although half have lower expression in tumors, including *JUNB* and *ATF3* (Figure S16c).

Additionally, we determined whether intergenic-exclusive DMRs were enriched near genes with particular biological functions using GREAT. Shared hypoDMRs were significantly enriched near genes with the function “glucose import” (*q*-value < 0.05, 7 DMRs and 7 genes). In contrast, shared hypermethylated DMRs were enriched near functions involved in morphogenesis and transcription (Figure S16d).

Finally, we identified small RNA whose gene bodies overlapped patient-matched DMRs. 72 small RNA genes overlapped a hypoDMR and 82 overlapped a hyperDMR in any patient, although only 6 and 26 did so in multiple comparisons, respectively. *MIR663A*, which belongs to a processed transcript gene, was hypermethylated in six patients (chr20:26188821-26188914). The hypermethylated region overlapped the promoter of the other isoforms and was annotated as a bivalent promoter; no other genes were in the vicinity. MIR663A is a known NSCLC tumor suppressor that acts through a variety of downstream targets, including TGFβ, p53, p21, and JunD [46–48], and its predicted targets include ACSL3, TGFB1, and HOXC10. MIR487A (chr14:101518782-101518862), which promotes tumor growth and metastasis in other cancers [49], was hypomethylated in four patients. For both, the expression level in LUAD samples was higher than in normal lung (*p* < 0.001, Wilcox test; Figure S17). Additionally, the snRNA RNVU1-8 (chr1:146551294-146551419), an alternative snRNA involved in mRNA processing [50], was hypermethylated in five patients, while the vault RNA VTRNA1-2 (chr5:140098509-140098598), which may contribute to multidrug resistance in cancer cell lines [51], was hypermethylated in two.

### Chromatin state in other tissues

Next, to better understand which regions may be susceptible to changes in DNA methylation in NSCLC, we found the epigenetic state of each DMR in adult lung and other normal human tissues profiled by the Roadmap Epigenomics Project. For each of the 127 consolidated epigenomes, the Roadmap Project assigned a composite epigenetic state generated from five core histone modifications (15-state model) using chromHMM. 98 epigenomes were also annotated with an 18-state model that includes H3K27ac.

We looked first at the epigenetic state of the DMRs in adult lung (epigenome E096). Compared to all potential DMR regions, hypoDMRs were enriched in enhancer states, the ZNF/Rpts state, and the heterochromatin state (**Figure 4**a). Although hypoDMRs were also enriched in Polycomb-repressed states, hyperDMRs were enriched > 50-fold for the bivalent promoter and enhancer states (14_TssBiv and 15_EnhBiv) and > 25-fold for the Polycomb-repressed state (16_ReprPC), as well as active promoter and enhancer states. In contrast, both hypo- and hyperDMRs were strongly depleted in the 18_Quies quiescent state (29% and 11% of all DMR bases, respectively, versus 54% of potential DMR bases), which represents an absence of histone modification ChIP- seq signal. The enrichments held true even when DMRs were split by feature overlap and/or the number of patients in which they were found, although hyperDMRs were more likely to overlap Polycomb-repressed regions the more frequently they occurred (Figure S18a). Enrichment profiles were comparable for DMRs from smoker patients and those from never smokers (Figure S19a). However, hyperDMRs exclusive to smokers demonstrated increasing enrichment over active enhancers the more they were shared across patients, in contrast to hyperDMRs observed in both smokers and never smokers (Figure S19b).

**Figure 4.**
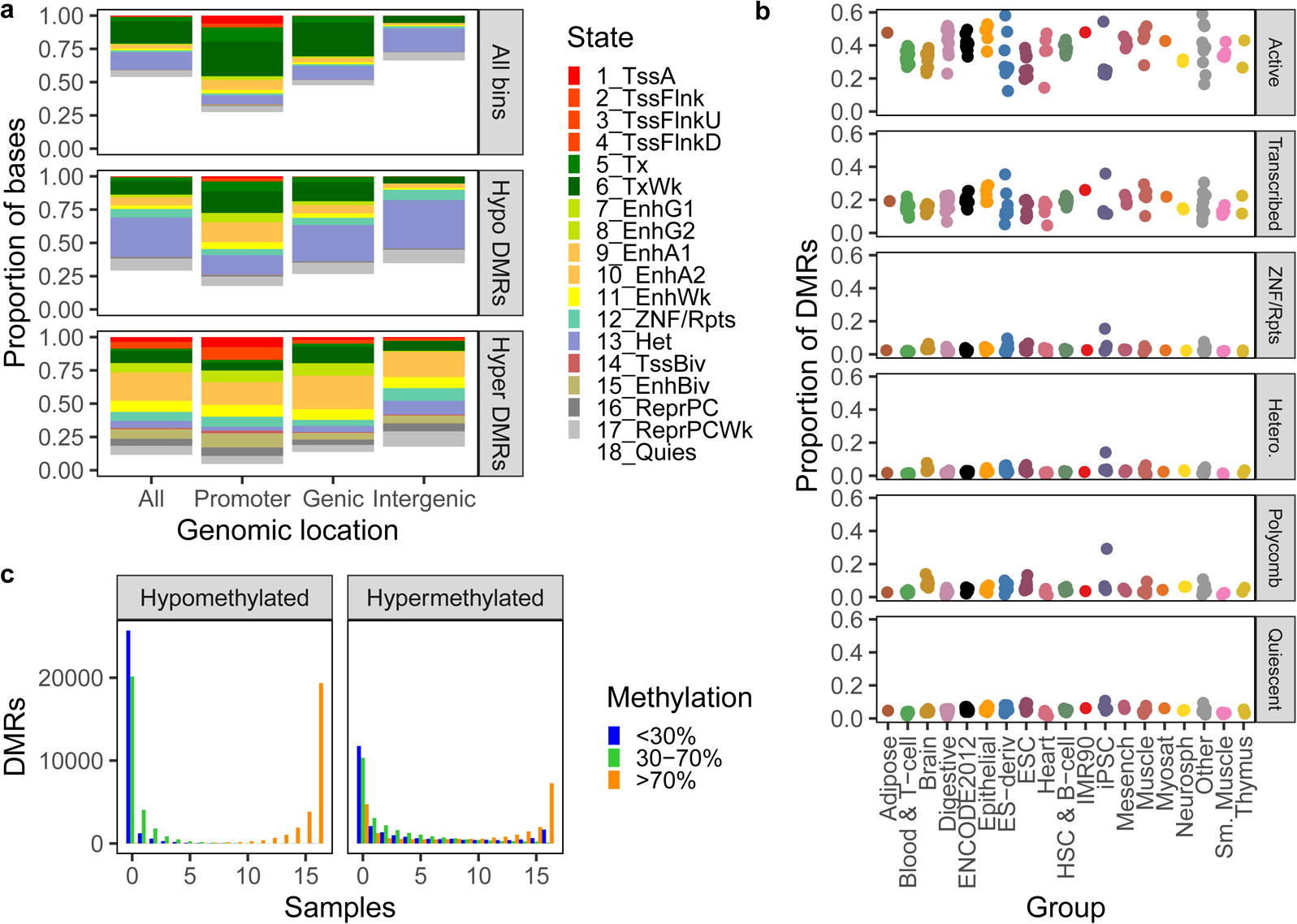
Epigenetic state of DMRs in Roadmap tissues. **A.** Proportion of Roadmap sample E096 (adult lung) bases in each 18-state chromHMM state, overall and overlapping hypo- or hyperDMRs, split by exclusive feature overlap. Genic DMRs overlap genes but not promoters, and intergenic DMRs do not overlap genes or promoters. The overall state in E096 was restricted to regions overlapping 500 bp bins that contained CpGs. 500 bp bins: promoter *n* = 453,670, genic *n* = 2,548,316, intergenic *n* = 2,267,290. HypoDMRs: promoter *n* = 4,663, genic *n* = 13,730, intergenic *n* = 14,472. HyperDMRs: promoter *n* = 12,628, genic *n* = 14,670, intergenic *n* = 7,128. **B.** The proportion of hypoDMRs in an active 15-state chromHMM state in each Roadmap sample, by sample group (columns) and its 18-state chromHMM state in E096 (rows) (see Materials and methods for composite state definitions). Hetero., heterochromatin. **C.** Distribution of the number of Roadmap samples in which each DMR had each average CpG methylation level, as assigned by methylCRF, split by DMR direction in the patient-matched NSCLC samples.

Overall, only 14% of hypoDMRs and 51% of hyperDMRs were in an active regulatory chromHMM state in normal lung, although shared hypoDMRs were more likely to be in those states. Repressed genomic regions that became activated in NSCLC may have been poised to do so because they were regulatory elements in another tissue. Indeed, 65% of hypo- and 94% of hyperDMRs were in an active regulatory state in a Roadmap sample besides adult lung (15-state model, excluding cancer cell lines), including > 50% of hypoDMRs in each repressed state in E096 (Figure S18b).

The proportion of DMRs in an active regulatory state in each sample varied by Roadmap sample group (Kruskal-Wallis *p* < 0.01, by DMR direction and composite chromHMM state in E096). Interestingly, hypoDMRs in the heterochromatin, Polycomb, or ZNF/Rpts states in adult lung were most likely to be in an active regulatory state in E022, an iPSC sample that did not have a higher overall proportion of bases in those states (Figure 4b). They were also more likely to be in active states in brain, ESCs, muscle, and other organs, and this was observed in hypoDMRs regardless of smoking status (Figure S19c). The three other normal lung samples profiled by the Roadmap project – fetal lung (E088), NHLF lung fibroblast primary cells (E128), and the IMR90 fetal lung fibroblast cell line (E017) – did not stand out in terms of the proportion of hypoDMRs in active regulatory states in those samples.

Additionally, we compared the epigenetic state of the DMRs in the A549 lung carcinoma cell line (E114) to the state in adult lung to determine whether changes in CpG methylation were matched by histone modification alterations. Both hypo- and hyperDMRs were enriched relative to genomic background in active regulatory and Polycomb-repressed states in A549 (Figure S18c). Regardless of epigenetic state in adult lung, hyperDMRs were far more likely to be in Polycomb-repressed states in A549.

Finally, we found the average CpG methylation level of each DMR in 16 Roadmap samples profiled with methylCRF, none of which was a lung sample. In general, DMRs that became hypomethylated in NSCLC were highly methylated in most or all other samples (see Materials and methods for complete listing of samples), suggesting that they lost regulation in NSCLC (Figure 4c). Interestingly, hypoDMRs were most likely to have lower methylation in epithelial samples (Figure S18d). In contrast, DMRs that become hypermethylated in NSCLC had a more variable profile and were hypo- or intermediately methylated in at least some other samples.

### Regions with high DMR density

As noted above, DMRs were not evenly distributed across the genome. Although the median DMR density per patient was 0.05% for both DMR directions (∼ 1 DMR per 2,000 bins; **Figure 5**a), hyperDMRs exclusive to never-smokers were less dense than hypoDMRs, whereas the ratio of hyperDMR to hypoDMR density was relatively comparable for smoker-exclusive DMRs (Figure S20a). DMR density also varied heavily by chromosome for some patients; for example, chromosome 19 had unusually high DMR density for many patients, reflecting its high gene density and unique epigenetic profile (Figure S20b). However, when DMRs were separated according to smoking status (DMRs unique to smokers, unique to never-smokers, or found in both), the hypoDMRs specific to never-smokers displayed high density across both never-smokers, particularly over chromosomes 16, 19, and 22 (Figure 5b). There were also instances of chromosomal outliers unique to a patient, such as chromosome 14 for Patient 9890 and chromosome 7 for Patient 8951. These may represent instances of genomic rearrangement or loss of a topologically associating domain (TAD) boundary during NSCLC transformation that permitted chromatin spreading, although in both cases, the MRE-seq and MeDIP-seq read density over the chromosome did not suggest a large-scale copy number change relative to other tumors (Figure S1).

**Figure 5.**
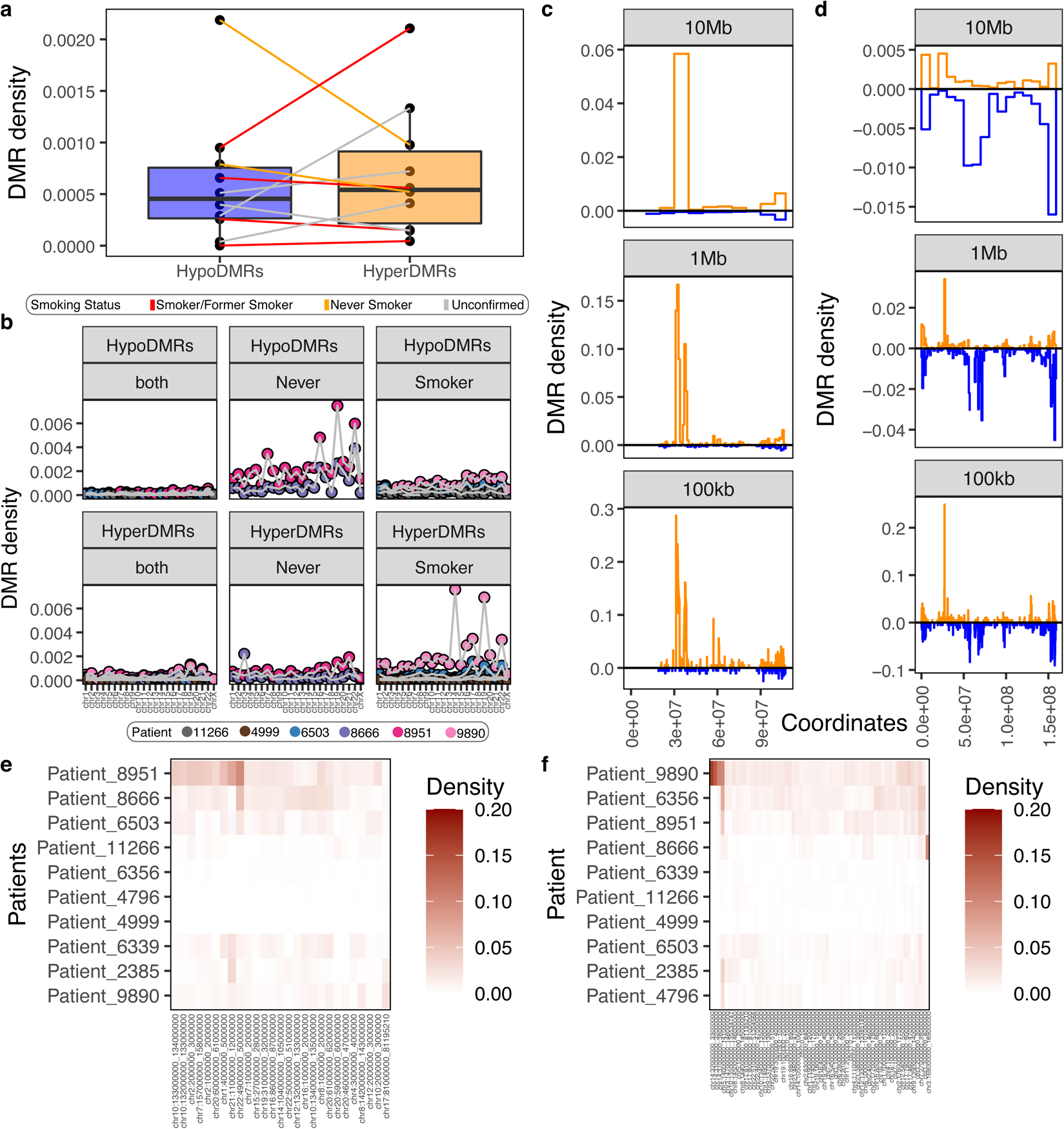
DMR density across patients, regions, and smoking status. **A.** Genome-wide DMR density per patient (number of DMRs versus number of 500 bp bins overlapping CpGs), by DMR direction, lines colored by smoking status of patient. **B.** DMR density per patient and chromosome, by DMR direction, for DMRs exclusive to smokers, exclusive to never-smokers, and DMRs identified in both. **C-D.** DMR density (number of DMRs versus 500 bp bins overlapping CpGs) at three levels of resolution for **(C)** Patient 9890, chromosome 14 and **(D)** Patient 8951, chromosome 7. **E-F.** Mean **(E)** hypo- and **(F)** hyperDMR density across 1 Mb windows in which the DMR density is > 1% in more than one patient or is one of the top 15 densest regions across samples, by patient.

To better understand the distribution of DMRs, we profiled their density at three resolutions ranging from 100 kb to 10 Mb, and identified windows with unusually high DMR densities (Figure 5c-d). For example, the 10 Mb window, chr14:30000000-40000000, had a hyperDMR density of 5.8% in Patient 9890, representing 81% (*n* = 1,075) of the hyperDMRs on that chromosome but only 11% of the potential DMRs. Most of the DMRs were within two ∼ 2 Mb-sized regions, with the density over the 1 Mb window, chr14:32000000-33000000, reaching 16.7%. Because many patient-specific DMRs were within high-density regions – for instance, 13% of hyperDMRs specific to Patient 9890 were within 1 Mb windows with a DMR density of > 10% (Figure S21) – most analyses focus on shared DMRs.

While many of the high-density windows were specific to a single patient, others were recurrent. At a resolution of 1 Mb (approximately the length of a TAD), 82 windows had a DMR density of > 1% in multiple patients (Figure 5e, 5f, Figure S20c). This included chr5:140000000-141000000, which was densely hypermethylated in 7 patients and is centered over a protocadherin gene cluster; chr7:27000000-28000000 over the HOXA cluster (> 1% hyperDMR density, 6 patients); and chr17:46000000-47000000 over the HOXB cluster (> 1% hyperDMR density, 4 patients). In contrast, the only protein-coding gene within chr1:4000000-5000000 (> 1% hypoDMR density, 4 patients) was *AJAP1*.

To further explore which regions of the genome may be predisposed to large-scale methylation alterations in NSCLC, we compared DMR density to features such as gene, CpG, and repeat density and the epigenetic state in normal lung. Many of these feature were highly correlated (Figure S22a), with the first several principal components separating 1) the proportion of the window in the quiescent state (18_Quies) in normal lung (E096) from gene density and active states, 2) active states from repressed states, and 3) Polycomb-repressed states from the heterochromatin and ZNF/Rpts states and repeat density (Figure S22b-e; Supplementary Text).

For both hypo- and hyperDMRs, DMR density in individual windows, the mean density across patients, and the number of patients in which the DMR density was > 1% were negatively correlated with the proportion of the window in the quiescent state (Figure S22f-g). HypoDMR density was most highly correlated with the heterochromatin, ZNF/Rpts, and Polycomb-repressed states as well as CpG density. HyperDMR density was most highly correlated with gene and CpG density and the Polycomb-repressed states, as well as active genic and enhancer states.

Interestingly, DMR density in both directions was also anticorrelated with the starting methylation level in the normal lung sample from that patient, as estimated by methylCRF. If changes in CpG methylation were stochastic, the starting methylation level should dictate the direction of methylation change; however, a higher normal lung methylation level lead to higher DMR density in both directions, likely because it is linked to gene and CpG density. Together, these results suggest that changes in methylation level during primary NSCLC transformation were enriched in regions with higher gene and CpG density, but that hypoDMRs were skewed towards heterochromatin-repressed regions while hyperDMRs were enriched over active regions, and that both were enriched over Polycomb-repressed regions.

### Methylation changes associated with tumor stage and type

Next, we identified DMRs associated with specific patient and/or tumor characteristics.

Although *TP53* mutations were previously shown to be associated with lung cancer grade [4], few patient-matched DMRs were exclusive to low-stage (1/1A) or high-stage (3A/3B) tumors (10 and 9 DMRs, respectively; see Materials and methods). However, 99 hypo- and 18 hyperDMRs were exclusive to acinar cell adenocarcinoma (*n* = 2 patients), and 37 hypo- and 389 hyperDMRs were exclusive to adenosquamous carcinoma (*n* = 2). For categories including Patient 4999, DMRs were counted as exclusive if they were present in all other members of that category. While many of the exclusive DMRs may be passenger events, the number of exclusive DMRs for adenosquamous carcinoma tumors was on the high end of a distribution generated with shuffled patient clinicopathologic data, suggesting that many are truly biology-specific (**Figure 6**a).

**Figure 6.**
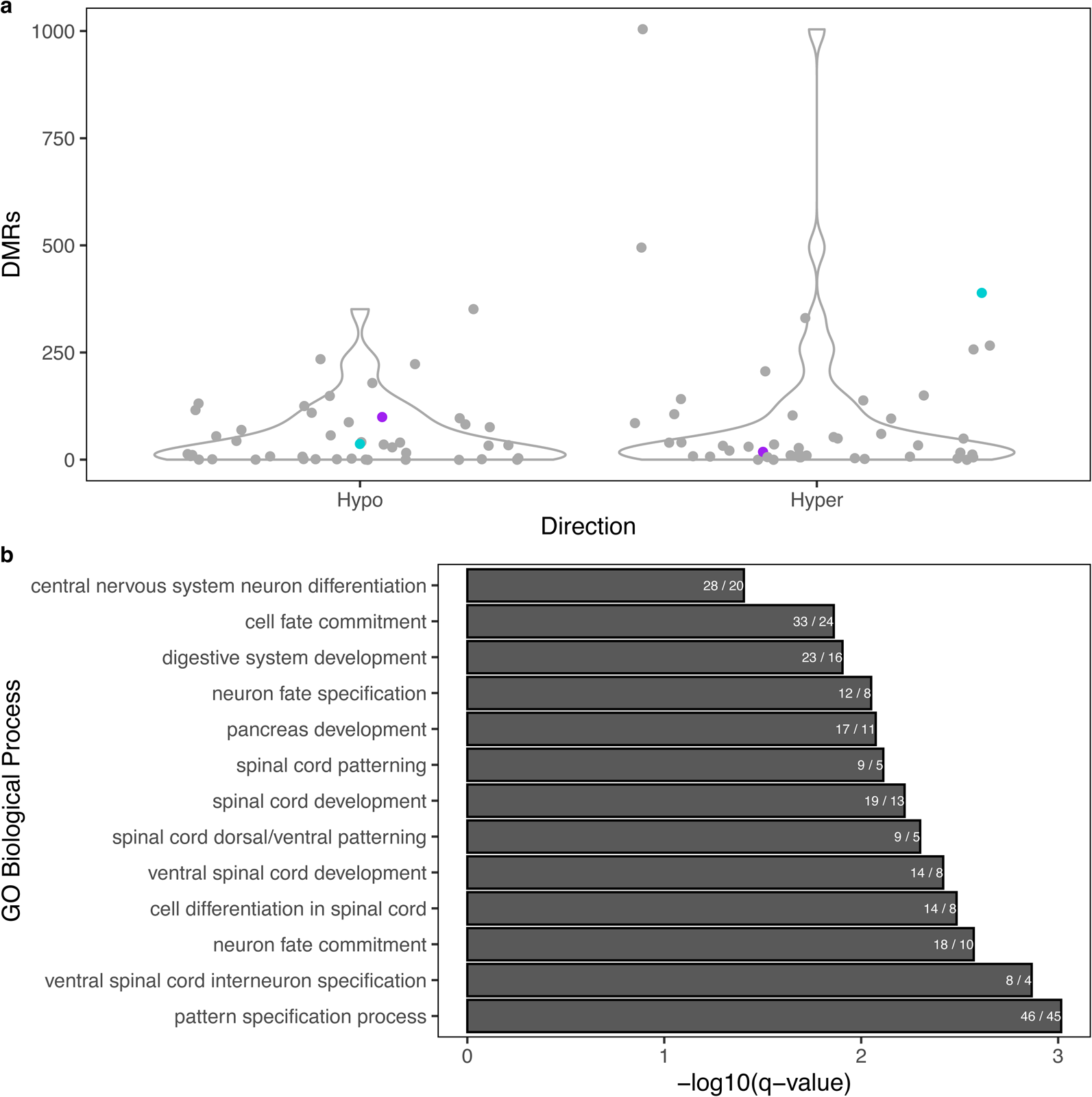
Tumor histological subtype-exclusive DMRs. **A.** Number of DMRs exclusive to acinar cell carcinoma (purple) or adenosquamous carcinoma (turquoise) using true tumor subtype or all possible pairs of patients (grey), by DMR direction. **B.** Top 20 significant GO Biological Processes for hyperDMRs specific to adenosquamous carcinoma, as identified by GREAT (see Materials and methods). Terms are ordered by FDR-corrected binomial *q*-value and are labelled by the number of DMRs over the number of genes involved.

The adenosquamous carcinoma-specific hyperDMRs were significantly enriched for several GO Biological Processes related to nervous system development (GREAT; Figure 6b). A small set of genes is included in many of the terms, including *NKX6-2*, *PAX6*, *LHX3*, *SOX1*, *ISL1*, and *HOXD10*. 44% (*n* = 186) of the adenosquamous-specific DMRs overlapped promoters. These included *ZSCAN30* and *ZIK1*, whose promoters became hypermethylated only in adenosquamous carcinoma.

### Repeat activation in NSCLC

Finally, we looked at methylation changes over repeats during NSCLC transformation. This is a relatively understudied aspect of cancer biology, especially considering that repeats comprise half of the human genome and can serve as alternative promoters and enhancers. Utilizing MeDIP-seq and MRE-seq data allowed us to better explore this component of the genome, as 53% of the CpGs profiled by methylCRF overlap a repeat, versus 16% of those profiled by TCGA.

As noted above (Figure S11a), 46% of hyperDMRs (*n* = 11,239) and 73% of hypoDMRs (*n* = 20,685) identified in this study overlapped a repeat. Across the 10 patients, this corresponded to 50,699 repeats that overlapped a patient-matched DMR, including 35,414 overlapping hypoDMRs and 15,652 overlapping hyperDMRs. 12% and 22% of the repeats overlapped a DMR in multiple patients, respectively (maximum 6 and 8 patients; **Figure 7**a). This included a DMR far from any gene that overlapped adjacent L1M5, LTR24C, and AluY elements and is hypomethylated in five patients (chr10:67554500-67555000).

**Figure 7.**
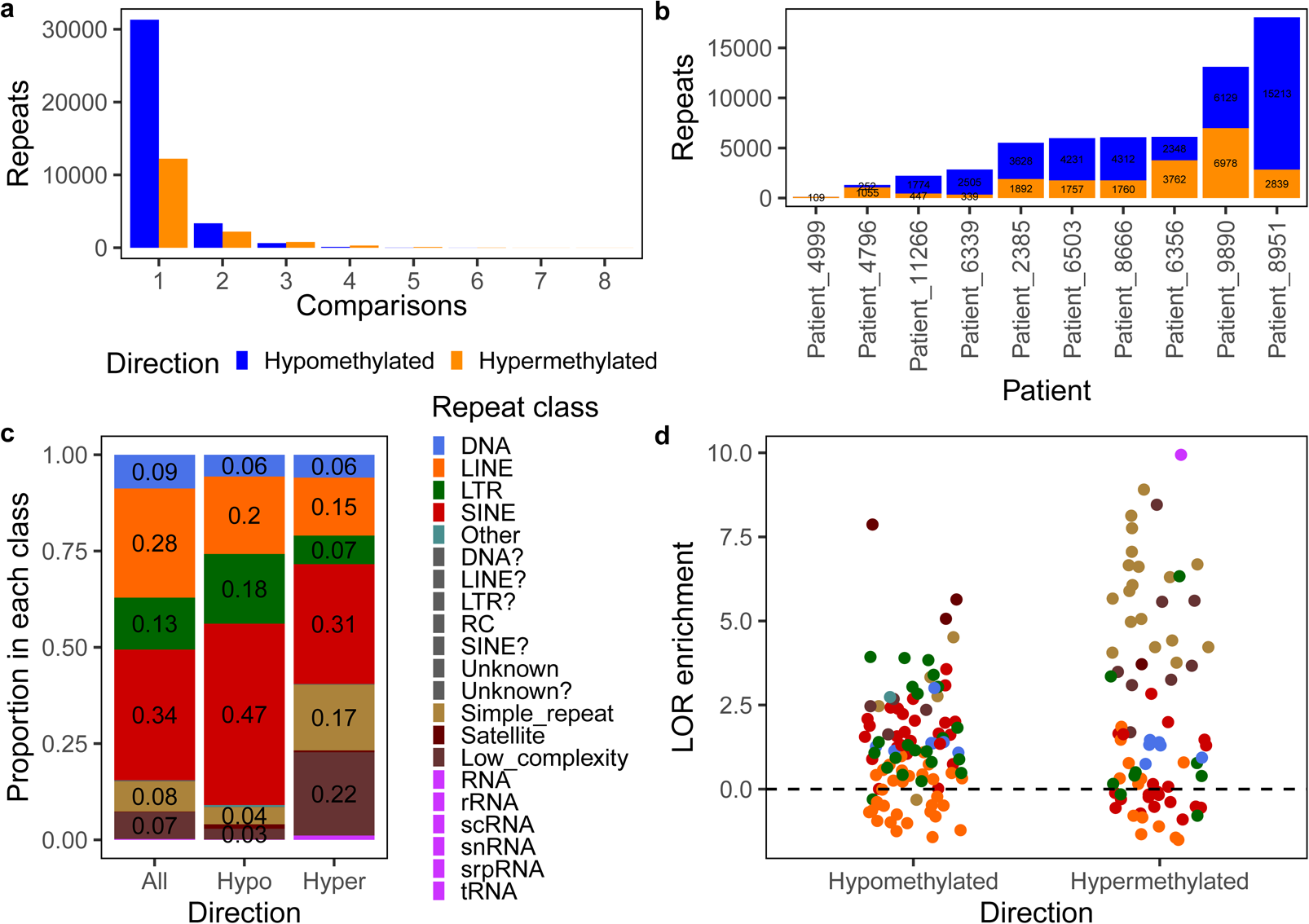
Repeat overlap with patient-matched DMRs. **A.** Number of comparisons in which each repeat overlapped a DMR, by DMR direction. **B.** Number of repeats overlapping DMRs per patient. Bars are labelled with the number of repeats. Counts < 100 are not shown. **C.** Proportion of repeats in each class, overall and overlapping hypo- or hyperDMRs shared between patients. Proportions < 0.02 are not shown. **D.** Log odds ratio enrichment of shared DMRs over repeat subfamilies, colored by repeat class, by DMR direction. Only subfamilies overlapping > 5 DMRs across all patients are shown. Dashed line represents no enrichment or depletion.

In a single patient, the number of repeats overlapping DMRs ranged from 8 to 15,213 for hypoDMRs (mean 4,040) and 109 to 6,978 for hyperDMRs (mean 2,093.8; Figure 7b), reflecting the landscape of CpG methylation changes in those patients.

Repeats were divided into several classes of retrotransposons, DNA transposons, and other repetitive elements. When compared to the number of repeats per class, SINE and LTR class elements were enriched among those overlapping shared hypoDMRs (Figure 7c). In contrast, low complexity regions and simple repeats were enriched among those overlapping hyperDMRs, and DNA and LINE class elements were depleted from both sets. These patterns held true when all patient-matched DMRs were considered (Figure S23a).

Next, we looked at CpG methylation changes over repeat subfamilies during NSCLC transformation. Several of the youngest TE subfamilies had the highest overall methylation in normal lung samples, including SVA and AluY subfamilies and LTR12C (Pearson correlation *p*-value > 0.05, median Jukes-Cantor distance vs. median methylation level; Figure S23b). Additionally, TE subfamilies with higher methylation in normal lung exhibited less variation in methylation level between normal samples (Pearson correlation −0.53, *p*-value < 0.001; Figure S23c), reflecting the importance of DNA methylation as a repressive tool. Younger TE subfamilies lost more CpG methylation on average in NSCLC (Pearson correlation −0.30, *p*-value < 0.001; Figure S23d), although interestingly, this was not primarily due to their higher starting methylation level, as there was little correlation between the median methylation level in normal samples and the change between normal lung and tumors (Pearson correlation 0.12, *p*-value < 0.001).

As expected, the variation in repeat subfamily methylation levels across samples increased in tumors (Figure S23e), although it was higher for those with greater variation in normal lung (Pearson correlation 0.75, *p*-value < 0.001; Figure S23f). Interestingly, many of the subfamilies with the largest median change in methylation between normal lung and tumor samples exhibited a bimodal distribution in tumors, with the same subfamily exhibiting increased methylation in some tumors and decreased methylation in others (Figure S23g).

In addition, we calculated the enrichment of DMRs over repeat subfamilies compared to their length to determine which were most prone to concentrated DNA methylation alterations. The majority of repeat subfamilies overlapped DMRs: 74% of all subfamilies and 85% of TE subfamilies overlapped hypoDMRs (*n* = 1,032 of 1,397 and 824 of 968, respectively), and 63% and 69% overlapped hyperDMRs (*n* = 877 and 665, respectively).

When considering all DMRs, the most highly enriched subfamilies in both directions were tRNA, low complexity, satellite, and simple repeats (Figure S24a). However, when restricted to DMRs shared between patients, TE subfamilies were better represented among those overlapping hypoDMRs, particularly ERV1 subfamilies (Figure 7d). Highly enriched subfamilies included LTR1, LTR1D, LTR12C, LTR17, MER45A, MER50, MER52A, MER52D, AluY, AluYc, AluYk4, and SVA_F. 11% of LTR12C elements (*n* = 278) overlapped a hypoDMR in at least one patient, and 53 were hypomethylated in multiple (LOR 3.04, *n* = 45 DMRs; maximum 3 patients, *n* = 8). This included the smoker-specific DMR in the *ARRB2* promoter mentioned above. Similarly, 10% of MER52D elements (*n* = 57) and 9% of MER52A elements (*n* = 149) overlapped a hypoDMR in at least one patient.

In contrast, the LTR subfamily MER57E3 was enriched in shared hyperDMRs (LOR 6.33, *n* = 7 DMRs). Of the eight MER57E3 elements that overlapped DMRs in multiple comparisons, most were within the first intron of a ZNF, pseudo-, or non-coding gene; they were also often adjacent to a MER21C element. MER57E3 had the lowest methylation level of any TE subfamily in normal lung samples (median 49.8% methylation), likely due to its overlap with promoter elements [52].

There was a moderate correlation between the CpG density of TE subfamilies and their enrichment for hypoDMRs (Spearman’s rho 0.47, *p* < 0.001), which was less true for hyperDMRs (rho = 0.17, *p* < 0.001; Figure S24b). However, although several of the most enriched subfamilies were relatively young, there was little correlation between TE subfamily age and DMR enrichment (Spearman’s rho < 0.2 in both directions). There was also little correlation between DMR enrichment and overall changes in CpG methylation level across the subfamily (Pearson correlation < 0.25 for hyperDMRs, *p*-value > 0.1 for hypoDMRs).

A recent paper used TCGA data to identify TEs that spliced into oncogenes to create alternative isoforms in cancer [15]. Four of the TEs identified in that study overlapped patient-matched hypoDMRs here, although only in one patient each. They included an intergenic AluSp element (chr19:15439585-15439882) that spliced into *BRD4*, as well as an intronic AluSp element (chr20:490814-491118) that formed an alternative *CSNK2A1* transcript in hundreds of TCGA LUAD and LUSC samples.

## Discussion

In this study, we use two complementary, genome-wide DNA methylation profiling techniques to interrogate DNA methylation changes in primary NSCLC and to detect local differentially methylated regions (DMRs) in comparison to patient-matched non-malignant lung tissue. Methylation changes in primary NSCLC were highly heterogeneous. As expected, tumors lost methylation over intergenic regions and repeats while gaining methylation over promoters. HOX and protocadherin clusters, as well as several lung biomarkers and oncogenes, were recurrently targeted by promoter methylation changes, as were small RNA. However, we also identified recurrent, intergenic DMRs, which may represent distal regulatory elements affected in NSCLC. These regions could potentially serve as biomarkers in cell-free DNA assays, for example, for early lung cancer detection.

Many of the genes identified in [20] as frequently methylated in the lung adenocarcinoma CIMP-H cluster, which were enriched for WNT pathway genes, were also hypermethylated in our study. As noted, *HOXA9* promoters overlapped hyperDMRs in 7 patients in our study, *SOX17* promoters in 5, and *GATA2* promoters in 4. *CDKN2A*, which is frequently methylated in lung adenocarcinoma and is inactivated by epigenetic silencing in 21% of squamous cell carcinomas [19], overlapped hyperDMRs in 3 patients. *STK11*, a frequently mutated tumor suppressor gene, overlapped hyperDMRs in 5 comparisons, but only over the promoter of a shorter isoform. In contrast, the most common driver genes in NSCLC, *EGFR*, *KRAS*, and *TP53*, rarely or never overlapped DMRs. Interestingly, one of the most commonly methylated genes in this study, *SEPT9*, is the basis for an FDA-approved, methylation-based colorectal cancer detection test (Epi proColon) and is being explored as a biomarker for lung cancer [53, 54]. Promoter methylation of *SOX17* also shows promise as a lung cancer biomarker [55].

Pathways related to cell signaling, cell cycle and proliferation, adhesion, chromatin and RNA splicing factors, and the oxidative stress response are commonly altered in lung adenocarcinoma, with 76% of tumors exhibiting RTK/RAS/RAF pathway activation [20][5][4]. In our study, similar pathways were enriched among genes with DMRs in their promoters. In line with previous studies, we also demonstrated recurrent gain of methylation over regions marked with Polycomb repressive marks, which were hypomethylated in most tissues [9]. Previous research has also indicated that enhancers lost in cancer target cell fate-specifying genes and are lineage-specific, while those activated in cancer are more universal and target growth-related genes [25, 56]. Here, recurrent hypoDMRs, including those in intergenic regions, were enriched for binding motifs for the AP-1 complex, as well as the lung development transcription factor NKX2-1, which is both lineage-specific and related to development.

Additionally, we identified DMRs exclusive to both tumor subtype and smoking status. While underpowered to reach significance due to small sample sizes, we did observe that smoker patients generally exhibited fewer DMRs than those seen in never-smokers, with the exception of Patient 9890. In addition, never-smokers tended to have a higher percentage of hypoDMRs than smokers, and more frequently displayed recurrent promoter hypomethylation despite fewer samples (*n* = 2 never-smokers, *n* = 4 smokers). These observations are consistent with the hypothesis that smoking-induced DNA damage recruits DNMT1 and leads to local increased methylation at repaired sites [57] and that loss of methylation may promote cancer development in never-smokers.

Alexandrov et al [7] identified only 434 differently methylated CpGs based on smoking status, most of which were in genes with no know cancer function. Similarly, although Freeman et al [58] identified 263 CpGs significantly associated with smoking in the TCGA LUAD and LUSC methylation data, only 5 replicated in an independent dataset. Few genes identified in earlier studies as preferentially mutated or silenced via promoter methylation based on smoking status overlapped DMRs in our study, including *p16*, *APC*, and *MLH1/2* [6]; *MICAL3* [5]; *CST6*, *EMILIN2*, *LAYN*, and *MARVELD3* [59]; and *RASSF1*, *MGMT*, *RARB*, *DAPK*, *WIF1*, and *FHIT* [60]. *CPEB1*, which was identified as specific to smokers in [59], and *CDKN2A*, identified in a meta-analysis of 97 studies by [60], overlap hyperDMRs here, although none are smoking status-specific. However, we identify DMRs overlapping additional genes known to be involved in NSCLC or smoking, such as *CYP1B1*.

Finally, we identified repeat subfamilies and individual repeats that were frequently affected by DNA methylation changes in NSCLC. Promoters overlapping repeats are more likely to be upregulated in cancer [33]. In our study, we observed a strong enrichment of hypoDMRs over ERV1 subfamilies. Although all major TE classes contain a sense-directional promoter [61], L1 elements are frequently 5’ truncated, while LTR repeats retain their regulatory elements after recombination removes the intervening viral genome [13], potentially explaining this phenomenon. We also observed hypomethylation over several young subfamilies, including SVA_F and several AluY subfamilies. Finally, we identified examples of repeats that have been shown to splice into known oncogenes in TCGA cancer samples to form alternative transcripts [15].

Here, we expand our knowledge of the epigenetic changes that occur during primary NSCLC transformation across lung adenocarcinoma subtypes and smoking status, potentially informing treatment strategies such as epigenetic therapy and immunotherapy. Future studies aimed at whole methylome profiling of patient matched normal and lung tumor samples of smokers and never-smokers should be conducted to further assess the frequency of DMR recurrence and specificity of methylation alterations for specific cohorts of patients.

## Materials and methods

### Samples and Clinical Data

Snap-frozen primary NSCLC tumor specimens (*n* = 17) and patient-matched non-malignant tissue (*n* = 10) along with de-identified pathology and demographic data were obtained from the Siteman Cancer Center Tissue Procurement Core Facility. All specimens were previously collected from surgical remnants with patient consent under an IRB-approved protocol. All frozen tissue specimens were sectioned and stained with hematoxylin and eosin to confirm histopathology and estimate tumor cell purity in each malignant tissue specimen. Serial frozen tissue sections were used for genomic DNA isolation using spin-column based purification (56304, Qiagen, Germantown, MD).Quantification of all DNA was performed by nanodrop spectrophotometry prior to methylation analyses.

Patient demographics and pathology data, including smoking status, pathologic diagnosis, stage, and tumor purity, were provided by the Tissue Procurement Core. Smoking status and patient sex were confirmed via chart review by an “honest data broker”. For analyses between never-smokers and smokers, former smokers were included in the latter category. Samples listed as “Acinar adenocarcinoma”, “Acinar cell carcinoma”, and “Acinic cell adenocarcinoma” were combined into a single category, “Acinar cell carcinoma”, for all analyses. Specimens for which demographic or pathology data could not be obtained were excluded from the relevant statistical tests. (see Supplementary Table 1)

### Features

GENCODE v19 comprehensive gene features and CpG islands (cpgIslandExtUnmasked) were downloaded from the UCSC Table Browser. Promoters were defined as 2000 bp upstream and 500 bp downstream of GENCODE transcription start sites.

A lookup table associating each Ensembl transcript ID with gene ID, name, and biotype was downloaded from the UCSC Table Browser. Chromosome and promoter overlap with CpG islands were added to each transcript.

The NCBI gene alias table (Homo_sapiens.gene_info.gz) was downloaded from NCBI.

The hg19 RepeatMasker rmsk file was used to identify repeats.

### MRE-seq and MeDIP-seq

MRE-seq and MeDIP-seq were performed on all samples using the procedures described in [62]. MRE-seq was performed with four enzymes: HpaII, SsiI, Hin6I, and HpyCH4IV (R0171S, R0551S, R0124S, R0619S, NEB, Ipswich, MA). A base offset of 3 was used during sequencing.

Adapter trimming, read alignment to the hg19 genome with BWA, and methylQA post-processing were also performed as described in [62]. methylQA produced a bed file of unique alignments (mapping quality ≥ 10), virtually extended, for each MeDIP-seq alignment file. For MRE-seq, it removes reads that do not map to MRE cut sites. Chromosomes 1-22, X, Y, and M were included.

MRE-seq and MeDIP-seq read counts per 500 bp bin were generated using M&M. MeDIP-seq read counts were normalized to RPKM using the total number of input reads. For MRE-seq, bins were restricted to those containing an MRE cut site, and read counts were normalized to RPKM using the total number of reads mapped.

PCA was performed on all samples using the MeDIP-seq and MRE-seq RPKM per 500 bp bin as features with the prcomp() function. Matrices were scaled and centered, and only bins with variation across samples were included. The variance explained by each PC was calculated from the standard deviation.

### methylCRF

methylCRF was performed on the 28 primary tumor and normal lung samples to estimate the methylation level at each CpG using MRE-seq and MeDIP-seq data. The methylCRF package was downloaded and installed from http://methylcrf.wustl.edu/. methylCRF was run using perl 5.14.4, R 3.3.0, and samtools 1.3.1.

MRE normalization was performed on aligned MRE-seq reads as recommended, using a quality score of 10 and a base offset of 3 for sam2bed.pl and the 4-enzyme MRE fragment file as input for MRE_norm.pl.

methylCRF was performed on normalized MRE-seq read counts and MeDIP-seq extended alignments in bed format using the H1ES model-specific files, the hg19 genome-specific files, the 4-enzyme virtual digest file, and a gap size of 750 bp. Methylation estimates were assigned for 28,085,255 CpGs across chromosomes 1-22, X, Y, and M (chromosome M always missing methylation value). Chromosomes 1-22, X, and Y were used for all methylCRF analyses.

### Methylation distribution in 1 kb windows

Non-overlapping 1 kb windows were generated using *bedtools makewindows* across chromosomes 1-22, X, Y, and M. Windows were intersected with CpG methylation level estimates generated by methylCRF using *bedtools intersect*, and the mean methylation over all CpGs within each window was calculated for all samples. Windows that did not overlap a CpG were excluded from analyses.

Principal components analysis was performed on windows with variation between samples using the prcomp() function. Matrices were scaled and centered. The variance explained by each PC was calculated from the standard deviation. Group centroids were identified by calculating the mean along each PC by malignant status. The distance to the group centroid for normal and tumor samples was calculated using Euclidean distance over all PCs.

A Pearson correlation distance matrix between all samples across the 1 kb windows was generated.

### Average methylation over genomic features

The average methylation level over each feature was calculated by first internally collapsing features to reduce overlapping bases using *bedtools merge*, then intersecting the merged features with the CpG methylation level estimates generated by methylCRF using *bedtools intersect*. Then, the mean methylation level over all CpGs overlapping the feature was calculated for all samples.

### MRE cut site saturation

The number of sampled restriction enzyme cut sites and the number of cut site-filtered MRE-seq reads per sample was obtained from the methylQA reports.

Cut sites specific to normal and tumor samples were identified by comparing the CpGs included in the MRE-seq CpG bedGraph files output by methylQA. The number of normal and tumor samples in which each CpG was sampled was counted. Cut sites overlapping each genomic feature were identified using *bedtools intersect*, and the distribution was compared to that of all possible restriction enzyme sites in the genome (*n* = 10,214,062).

### Comparison to previously published methylCRF data

CpG methylation level estimates generated by methylCRF for 16 samples profiled in [63] (GEO ID GSE86505) were downloaded from https://wangftp.wustl.edu/~jli/final_hub/methylCRF_score/, and cancer types were extracted from sample names.

methylCRF was performed on three samples profiled by the Roadmap Epigenomics Project using MeDIP-seq and MRE-seq (GEO accessions GSM613862, GSM613842, GSM669614, GSM669604, GSM707021, GSM707017). methylQA was performed on the MeDIP-seq read alignments as recommended (http://methylqa.sourceforge.net/tutorial.php) after adding the “chr” prefix to each chromosome in the header with *samtools reheader*. MRE normalization was performed using the 3-enzyme fragment file with a base offset of 3, and methylCRF was performed using the 3-enzyme virtual digest file. The average methylation in 1 kb windows across the genome was calculated as above, excluding windows that did not overlap a CpG.

To compare to the samples profiled in this paper, which were processed with 4 restriction enzymes, methylCRF was performed on two samples (P4796_N_UC and P4796_T_UC) considering only 3 enzymes (excluding HpyCH4IV). MRE normalization was repeated using the 3-enzyme fragment file, and methylCRF was performed using the 3-enzyme virtual digest file. The average methylation in 1kb windows across the genome was calculated as above, excluding windows that did not overlap a CpG. Principal components analysis was performed as above on all primary NSCLC samples, the three Roadmap samples, and the two re-processed primary NSCLC samples simultaneously.

### M&M

M&M (R package methylMnM) was used to identify DMRs between samples using MRE-seq and MeDIP-seq data. First, the functions *countcpgbin* and *countMREcpgbin* were used to count the number of CpGs and restriction enzyme cut sites in 500 bp bins across the genome (chromosomes 1-22, X, Y, and M; *n* = 6,166,049 bins). Cut sites for the 4 enzymes used in MRE-seq were considered, and a blacklist file was provided (hg19 DAC Blacklisted Regions, e.g., wgEncodeDacMapabilityConsensusExcludable from the UCSC Genome Browser).

Then, 500 bp DMRs were identified using the read alignment bed files output by methylQA for MRE-seq and MeDIP-seq from two samples. The functions *countMeDIPbin* and *countMREbin* were used to produce normalized read counts per bin for each sample. Default values were used for *MnM.test*, *MnM.qvalue*, and *MnM.selectDMR*, and no PCR threshold was set.

DMRs were identified between patient-matched normal lung and tumors from the same patient (*n* = 11), between all pairs of normal samples (*n* = 55), and between all normal-tumor sample pairs from different patients (*n* = 176).

### DMR *q*-value threshold

The DMR *q*-value threshold was selected from among four potential thresholds (1E-2 to 1E-5). For each threshold, the false positive ratio was calculated for patient-matched normal vs. tumor comparisons as the mean number of DMRs identified between the given normal and all other normal samples divided by the number of patient-matched DMRs. The proportion of DMRs shared between patient-matched comparisons was also calculated for each threshold.

Additionally, all DMRs between patient-matched normal-tumor samples were intersected with the CpG methylation levels generated by methylCRF for each sample in the pair, and the mean methylation level across all CpGs within each DMR was calculated for each sample. The change in mean methylation level between patient-matched normal and tumor samples was calculated for each DMR identified in that pair. The proportion of DMRs with > 10% methylation change in either direction was calculated for each *q*-value threshold.

DMRs with a *q*-value < 0.001 from all comparisons were combined and assigned a unique ID by position. First, adjacent DMRs were merged and assigned a number based on location in the genome. Then, individual 500 bp DMRs were assigned a second number representing their position within the merged block. IDs are consistent across samples comparisons. There are 1,817,809 instances of 245,723 unique DMRs across all comparisons.

Because all patients with patient-matched normal lung and tumor samples are female, DMRs on chromosome Y (*n* = 1,112 instances of 80 unique DMRs, 0.03% of all DMRs) were excluded from downstream analyses.

### Feature overlap

To find the genomic context of DMRs identified in this study, all unique DMRs were intersected with GENCODE genic features, intergenic regions, CpG islands, and repeats using *bedtools intersect*. As background, genome-wide 500 bp bins (M&M input) were also intersected with the features. Only bins on chromosomes 1-22 and X that overlap a CpG (e.g., potential DMRs) and did not overlap a blacklisted CpG were considered (*n* = 5,269,276). The proportion of DMRs and 500 bp bins overlapping each feature category was calculated, and the log odds ratio enrichment of DMR overlap versus bin overlap was calculated for each feature.

Intersections of unique DMRs with individual CpG islands were identified by CpG island ID (*n* = 524, chromosomes 1-22 and X). The proportion of unique CpG islands overlapping a DMR (any or shared between multiple patient-matched comparisons) was calculated.

The number of GENCODE genes whose promoter(s) overlaps a patient-matched DMR (any or shared by multiple patients) was calculated and compared to the number of genes whose promoter(s) overlap a 500 bp bin containing a CpG (e.g., potential DMR) by gene biotype. Greater than 99.9% of transcripts and genes contain a potential DMR in their promoter(s) (chromosomes 1-22, X), including > 99% of all gene and transcript biotypes except mitochondrial rRNA/tRNA. Unique transcripts and genes were identified by ID, not gene name.

In addition, exclusive feature overlap categories were assigned to each DMR: promoter (DMRs overlapping a promoter), genic (DMRs overlapping a gene but not a promoter), and intergenic (intergenic DMRs that overlap neither a promoter nor a gene).

For each GENCODE transcript and gene, the number of unique patient-matched DMRs overlapping the promoter(s), the number of patient comparisons in which the promoter(s) overlapped a DMR, and the total number of DMR overlaps across comparisons was calculated by direction of methylation change.

For each DMR, the distance to the closest gene on either strand was found using *bedtools closest*, reporting all ties.

### Gene sets

Several gene lists were used to identify genes previously recognized as involved in cancer. A list of genes previously identified as altered in lung adenocarcinoma and squamous cell carcinoma was compiled from 9 publications [4–6, 8, 19, 20, 59, 60, 64]. The Cancer Gene Census database (v79) was downloaded from COSMIC. The Epifactors database (v1.7.3) was downloaded from the FANTOM5 consortium. CTAs (gene names plus alternative names) were obtained from the CTDatabase website (http://www.cta.lncc.br/; accessed 2/13/2019). Additionally, genes listed as “cancer/testis antigen” or “cancer testis antigen” in the NCBI gene alias table were also included. The top 100 most highly expressed genes in lung were obtained from the GTEx data portal, excluding mitochondrial genes (https://gtexportal.org/home/eqtls/tissue?tissueName=Lung).

Genes included in sets of interest were identified by gene name and linked to gene IDs using the GENCODE lookup table. For gene names not included in the GENCODE table, where possible, the name was identified among synonyms in the NCBI gene alias table and linked to the canonical name. For GTEx genes, the gene ID (without version suffix) provided by GTEx was used to restrict gene names to the correct Ensembl ID in the lookup table. In the one case where the GTEx-provided ID did not match any IDs for the gene name in the GENCODE lookup table (“TXNIP”), the ID from the lookup table was used instead. Four genes previously identified in NSCLC, 7 Cancer Gene Census genes, 5 Epifactors genes, 51 CTAs, and one GTEx gene were not present in either the GENCODE lookup table or the NCBI gene alias table.

GENCODE IDs were used to link genes between gene sets and to link genes and transcripts with gene set membership. The CGC “Role in Cancer” field was used to identify TSGs and oncogenes. The Epifactors “Function” field was used to identify gene function.

### Alternative promoters

To identify potential instances of promoter switching in primary NSCLC, all genes with multiple promoters overlapping DMRs and at least one hypo- and hypermethylated DMR in the same patient-matched comparison were identified. Genes that exhibited that combination in multiple patient-matched comparisons were selected for further investigation.

### Small RNA

miRNA and small RNA whose gene body overlapped patient-matched DMRs (transcript biotype: “miRNA”, “misc_RNA”, “snRNA”, “snoRNA”, “rRNA”, “Mt_tRNA”, “Mt_rRNA”) were identified. Predicted miRNA targets were obtained from miRDB information on the miRBase website.

### DAVID

DAVID was performed on lists of genes with hypo- or hypermethylated patient-matched DMRs in their promoter(s), scaled by the number of comparisons in which the gene overlaps a DMR. Genes were looked up using “official gene name”. The categories OMIM_DISEASE, COG_ONTOLOGY, UP_KEYWORDS, CHROMOSOME, BBID, BIOCARTA, KEGG_PATHWAY, and UP_TISSUE were tested. To accommodate the list of genes with hypermethylated DMRs in the promoter in any patient, the list was first split into six smaller lists, and gene IDs that were not mappable by DAVID were excluded. The remaining gene names were combined into a single list and analyzed as before.

Terms with a Benjamini-corrected *p*-value < 0.05 were considered significant.

### Roadmap Epigenomics Project epigenomes

Clinicopathologic data for the Roadmap Epigenomics Project consolidated epigenomes was obtained from the REMC data portal (https://egg2.wustl.edu/roadmap/web_portal/meta.html, Consolidated_EpigenomeIDs_summary_Table). Cancer cell lines were those with “leukemia” or “carcinoma” in the epigenome name (*n* = 5), which were excluded from analyses unless otherwise stated. Group colors are those assigned in the Roadmap publication [65].

Roadmap Epigenomics Project chromHMM annotations were downloaded from the REMC data portal, including for 127 samples annotated with the 15-state model and 98 samples annotated with the 18-state model (http://egg2.wustl.edu/roadmap/data/byFileType/chromhmmSegmentations/ChmmModels/coreMarks/jointMod <u>el/final/all.mnemonics.bedFiles.tgz</u> and https://egg2.wustl.edu/roadmap/data/byFileType/chromhmmSegmentations/ChmmModels/core_K27ac/jointModel/final/all.mnemonics.bedFiles.tgz). DMRs were intersected with the annotations, and the length of each DMR annotated with each chromHMM state in each sample was calculated.

As background, the proportion of samples E096 and E114 (Lung and A549 EtOH 0.02pct lung cancinoma) in each 18-state chromHMM state over 500 bp bins containing a CpG (M&M input, chromosomes 1-22 and X) was calculated. For E096, 500 bp bins were also split into promoter-overlapping, genic-exclusive (do not overlap promoters), and intergenic-exclusive categories (do not overlap promoters or genes). The proportion of all samples in each 15-state chromHMM state over 500 bp bins containing a CpG (M&M input, chromosomes 1-22 and X) was also calculated.

Colors used for chromHMM states are those assigned in [65] for the 15-state and 18-state models. Active regulatory states were states 1-3 and 6-7 for the 15-state model and states 1-4 and 7-11 for the 18-state model. Additionally, other 18-state chromHMM states were assigned to composite states: transcribed (states 5-6) and Polycomb (states 14-17).

methylCRF CpG methylation level estimates generated for 16 epigenomes by the Roadmap Epigenomics Project were downloaded from the REMC data portal and reformatted into bed format (Sample IDs: E003, E027, E028, E037, E047, E053, E054, E055, E056, E057, E058, E059, E061, E081, E082). DMRs were intersected with the methylation levels using *bedtools intersect*. Then, the mean methylation level over all CpGs overlapping the DMR was calculated for all epigenomes. Methylation levels were divided into three categories: hypomethylated (< 30%), intermediately methylated (30-70%), and hypermethylated (> 70%). 8 DMRs did not overlap any CpG in the Roadmap files.

### HOMER

Known motif enrichment analysis was performed using HOMER *findMotifsGenome.pl* on each DMR set (hg19, parameters *-size given -nomotif*). DMR sets included hypo- and hypermethylated DMRs in exclusive promoter, genic, and intergenic categories, scaled by the number of patient-matched comparisons in which they were found. The knownResults.txt output files were used in downstream analyses. Only motifs with a Benjamini-corrected *p*-value < 0.05 were considered.

The transcription factor name was extracted from the “Motif Name” field in the knownResults.txt file. Transcription factors present in TCGA gene expression data were identified using case-insensitive exact name matches.

### GREAT

GREAT analysis (version 4.0.4) was performed on each DMR set using the basal plus extension model (5000 bp upstream, 1000 bp downstream, 1000000 bp max extension), including curated regulatory regions. All significantly enriched GO Biological Processes with the following thresholds were downloaded: FDR *p*-value < 0.05 by hypergeometric and binomial test, at least one observed gene hit, minimum 2 region-based fold enrichment (default). DMR sets included hypo- and hypermethylated DMRs in the intergenic-exclusive category, scaled by the number of patient-matched comparisons in which they were found.

### DMR density in windows

The average genome-wide DMR density for each sample comparison and direction of methylation change was calculated as the proportion of 500 bp bins overlapping a CpG that were called as DMRs. The average DMR density across each chromosome was calculated in the same manner.

Non-overlapping windows of size 10 Mb, 1 Mb, 100 kb, and 10 kb were created using *bedtools makewindows* across the genome, excluding chromosome Y and chromosome M. The number of hypo- and hypermethylated DMRs per window was calculated for each patient-matched comparison. The DMR density per window was calculated as the proportion of 500 bp bins overlapping a CpG called as DMRs, for each direction of methylation change.

The proportion of DMRs exclusive to a patient-matched comparison that fall within high-density windows was calculated for each comparison and direction of methylation change.

The CpG and repeat density for each window was calculated by normalizing by the length of the window. GENCODE transcripts overlapping each window were also identified, and the density of genes, protein-coding genes, transcripts, and protein-coding transcripts was calculated using the biotype information from GENCODE.

All windows were intersected with the CpG methylation level estimates generated by methylCRF using *bedtools intersect*. Then, the mean methylation level over all CpGs overlapping the window was calculated for all samples. For comparisons to DMR density, the average mCRF level was restricted to the normal sample for that comparison.

In addition, the proportion of each window annotated with each 18-state chromHMM state in Roadmap sample E096 was calculated.

PCA was performed on all 1 Mb windows using gene, CpG, and repeat density and the proportion of the window in each chromHMM state as features with the prcomp() function. Matrices were scaled and centered prior to transformation. Variance explained was calculated for each principal component. The Pearson correlation of each variable with the first three PCs was calculated.

The Pearson correlation of all 1 Mb window features with DMR density in individual comparisons, the number of comparisons in which the window had > 1% DMR density, and the mean DMR density across comparisons was calculated.

### Clinicopathologic data-specific DMRs

Patient-matched DMRs specific to patient clinicopathologic data categories were identified using frequency information. Never-smoker-specific DMRs were those found between the normal lung vs. tumor samples of both never-smokers and not present in smokers, while smoker-specific DMRs were found in 3-4 smokers and no never-smokers, ignoring presence in patients with unconfirmed smoking status. Low-stage DMRs were those found in 4-5 patients with tumor stage 1/1A but not those with tumor stage 3A/3B, and high-stage DMRs were those found in at least 3 patients with tumor stage 3A/3B but not those with tumor stage 1/1A, ignoring presence in patients with unconfirmed tumor stage. Tumor subtype-specific DMRs were those found in all patients with that tumor subtype and in no other.

To determine whether the number of smoking status-specific DMRs was likely to be identified by chance, the smoking status of the six patients with confirmed status was shuffled to create all possible permutations with 2 never-smokers and 4 smokers (*n* = 15). Then, the number of smoking status-specific DMRs was recalculated for each permutation. For subtype-exclusive DMRs, the number of exclusive DMRs was recalculated for all possible pairs of patients (*n* = 45) and compared to the true number for acinar/acinic cell adenocarcinoma and adenosquamous carcinoma.

GREAT analysis was performed on patient-matched DMRs exclusive to never-smokers, smokers, adenosquamous carcinoma, and acinic cell/acinar cell adenocarcinoma by direction of methylation change using the same parameters as above. GREAT analysis was also performed on DMRs between never-smoker and smoker normal samples that were not found between the never-smoker samples. The hypomethylated DMRs exclusive to each category did not return any significant GO Biological Process.

The proportion of smoking status-specific DMRs that fall within high-density 1 Mb windows of the same direction of methylation change in each patient was calculated.

To determine whether smoking was associated with DNA methylation alterations in normal lung samples, normal vs. normal sample DMRs were assigned to never-never, smoker-smoker, or never-smoker comparison categories (*n* = 1,214). DMRs found between never-smoker and smoker normal samples, but not between the never-smoker samples, were identified. Additionally, for each normal sample, the number of DMRs found between the sample and never-smokers and the sample and smokers was calculated and tested for significant differences using Wilcox tests. Example DMRs (*n* = 28) were identified as those found in three or more never-smoker vs. smoker comparisons, including both never-smoker samples, and not between the two never-smoker samples.

### TCGA

TCGA LUAD CpG methylation and gene, isoform, and miRNA expression data were downloaded from the TCGA legacy data portal using the gdc client and manifests generated through the GDC legacy data portal (https://portal.gdc.cancer.gov/legacy-archive/).

Clinicopathologic data for all downloaded LUAD files (*n* = 2,214) was obtained from the legacy API endpoint using the file ID, including file name, data category, data type, sample ID, sample type, and case ID. A summary of the number of samples and files by data type per case is provided (Supplementary Table 3). All samples with gene expression data also have isoform expression data and vice versa.

All 450K array methylation level files were downloaded (Data Type: “Methylation beta value”, Platform: “Illumina Human Methylation 450”; *n* = 507). The average CpG methylation level over DMRs identified in this study and GENCODE transcript promoters was calculated in all TCGA LUAD and matched normal samples. First, promoters or unique DMRs with *q*-value < 0.001 were intersected with the 450K array CpG methylation levels from all LUAD samples. Then, the mean methylation level over all CpGs within each DMR/promoter in each TCGA sample was calculated. DMRs or promoters without methylation data in any TCGA sample were excluded from analysis.

The mean methylation level over each patient-matched DMR was averaged across TCGA samples by sample type (matched normal lung or primary LUAD) and by DMR direction of methylation change, excluding samples where the DMR was missing methylation data. The mean methylation level over patient-matched DMRs identified in smokers and non-smokers (either at all or recurrently) was also calculated across TCGA samples, by sample type, DMR direction of methylation change, and TCGA smoker status designation, again excluding samples where the DMR was missing methylation data. TCGA samples were classified as either smokers (having a value in the metadata category “cigarettes per day” or “years smoked”) or non-smokers (empty values in both metadata categories).

To compare 450K array CpG coverage to methylCRF CpG coverage, TCGA CpG positions were extended 1 bp upstream, and CpGs without a chromosome were excluded (*n* = 65). Of the remaining 485,512 CpGs, 0.8% do not overlap a methylCRF CpG (*n* = 3,868). 98.3% of methylCRF CpGs (*n* = 27,603,511) do not overlap a TCGA CpG.

All normalized gene and isoform expression files were downloaded (Experimental Strategy: “RNA-seq”, Data Type: “Gene expression quantification” or “Isoform expression quantification”, filtered to files ending in “rsem.genes.normalized_results” or “rsem.isoforms.normalized_results”; *n* = 576 each).

Gene symbols for gene and isoform expression levels in TCGA were extracted from the gene ID in the gene expression files and the TCGA hg19 gaf file, restricted to “transcript” entries. Mappings between UCSC knownGene isoforms and Ensembl transcripts (hg19.knownToEnsembl) were obtained from the UCSC Table Browser, and isoform IDs used in TCGA were connected to Ensembl transcripts using abbreviated isoform IDs without version suffixes.

For select genes, average promoter CpG methylation, gene expression, and isoform expression levels were plotted for all TCGA samples, comparing LUAD primary tumors and matched normal lung. All GENCODE transcripts associated with the gene name were included in the promoter methylation analysis. All TCGA genes and isoforms associated with the gene name were included in the expression analyses. The difference in median average promoter methylation level or the fold change in median expression level was calculated between primary tumors and matched normal lung, and Wilcox tests were used to test for significant differences for each gene, transcript, or isoform. Where possible, isoform IDs were linked to GENCODE transcript IDs. Because some promoters do not overlap CpGs and some GENCODE gene names are not included in the TCGA expression data, all analyses could not be performed for all genes.

All miRNA files processed with miRBase v20 were downloaded (Experimental Strategy: “miRNA-Seq”, Data Type: “miRNA gene quantification”, filtered to “mirbase20”; *n* = 555 files). miRNA were linked to gene symbols using the TCGA hg19 gaf file and the NCBI gene alias table. The TCGA hg19 gaf file was restricted to “pre-miRNA” entries, and gene names and miRNA IDs were extracted from their respective columns. Additionally, genes with synonyms containing “hsa-“ were extracted from the NCBI gene alias table along with the associated gene name. Both sources were used to link miRNA IDs assigned by miRBase with gene names in the TCGA data.

The miRNA expression level of specific genes in TCGA LUAD and normal lung samples was plotted and tested for significant differences between the sample types using a Wilcox test. All miRNA IDs associated with the gene name were included.

### Repeats

Transposable elements were considered those with class: “DNA”, “DNA?”, “LINE”, “LINE?”, “LTR”, “LTR?”, “Other”, “RC”, “SINE”, “SINE?”, “Unknown”, and “Unknown?”. Only repeats on chromosomes 1-22 and X were considered.

The Jukes-Cantor evolutionary distance for each repeat was calculated from the substitution rate compared to the RepBase subfamily consensus sequence. The total length of bases overlapping each subfamily, the number of elements, median Jukes-Cantor distance, and the number of unique methylCRF CpGs overlapping each subfamily was calculated.

The average methylation over each subfamily in each sample was calculated by identifying all unique CpGs that overlap the subfamily, then calculating the mean methylation level assigned by methylCRF over all CpGs in each sample. Seventeen subfamilies overlap no CpG. For each subfamily, a Wilcox test was performed on the mean methylation level in normal vs. tumor samples. Only subfamilies with at least 10 CpGs were considered for downstream analyses.

The log odds ratio enrichment for each subfamily and DMR direction was calculated from the proportion of DMRs overlapping the subfamily versus the proportion of total repeat length represented by the subfamily. Enrichment was calculated for all DMRs and those found in > 1 patient.

For all repeats overlapping a patient-matched DMR, the number of unique DMRs overlapping the repeat, the total number of DMRs overlapping the repeat across comparisons, and the number of comparisons in which the repeat overlaps a DMR was calculated by direction of methylation change. The closest gene on each strand was identified using *bedtools closest*, including all tied genes, ignoring genes upstream of the repeat on the same strand and downstream of the repeat on the opposite strand. TEs overlapping hypomethylated DMRs present in > 1 comparison upstream of a gene on the sense strand (or downstream on the opposite strand, in the case of LINE elements) were identified.

A list of TEs that splice into oncogenes to form alternative isoforms in TCGA cancer samples was downloaded from [15] (Supplementary Table 2). The TE coordinates were lifted over to hg19 from hg38 using liftOver, and TEs that overlap hypomethylated DMRs in this study were identified.

Repeat class colors are those assigned on the WashU Epigenome Browser RepeatMasker track.

### Software

The following software packages were used: vegan 2.5.6, ggplot2 3.2.1, reshape2 1.4.3, plyr 1.8.4, grid 3.5.1, gridExtra 2.3, RColorBrewer 1.1.2, knitr 1.26, readxl 1.3.1, UpSetR 1.4.0, combinat 0.0.8.

## Ethical statement

An IRB-approved waiver of informed consent was obtained for all patient samples analyzed in the present study.

## Data availability

A public WashU Epigenome Browser datahub is available for the MeDIP-seq, MRE-seq, M&M, and methylCRF data generated in this study (https://epigenomegateway.wustl.edu/browser/?genome=hg19&hub=https://wangftp.wustl.edu/~epehrsson/NSCLC_primary/NSCLC_primary). Tracks are linked to sample and patient clinicopathologic data. In addition, all sequencing data generated as a part of this study have been deposited in Gene Expression Omnibus and are accessible under the ID: GSE210957.

## Code availability

Custom scripts generated for analyzing data in the present study have been made publicly available at https://github.com/jaflynn5/DNA-Methylation-Changes-in-NSCLC-by-Smoking-Status.

## CRediT author statement

**Jennifer A. Karlow**: Software, Validation, Formal analysis, Investigation, Data curation, Writing – Original draft preparation, Writing – Review and editing, Visualization. **Erica C. Pehrsson**: Software, Validation, Formal analysis, Investigation, Data curation, Writing – Original draft preparation, Visualization. **Xiaoyun Xing**: Methodology, Investigation, Writing – Original draft preparation. **Mark Watson**: Conceptualization, Investigation, Data curation, Resources, Writing – Original draft preparation, Supervision, Funding acquisition. **Siddhartha Devarakonda**: Investigation, Data curation, Resources, Writing – Original draft preparation. **Ramaswamy Govindan**: Conceptualization, Investigation, Resources, Writing – Original draft preparation, Supervision, Funding acquisition. **Ting Wang**: Conceptualization, Validation, Formal analysis, Investigation, Data curation, Resources, Writing – Original draft preparation, Writing – Review and editing, Supervision, Funding acquisition.

## Competing interests

The authors declare no conflicts of interest.

## Acknowledgements

J.A.K is supported in part by the Siteman Cancer Center Precision Medicine Pathway (T32CA113275). E.C.P. is supported by a Postdoctoral Fellowship, PF-17-201-01, from the American Cancer Society. E.C.P., J.A.K. and T.W. are supported by NIH grants R01HG007354, R01HG007175, R01ES024992, U01CA200060, U24ES026699, U01HG009391, and American Cancer Society RSG-14-049-01-DMC.

## Supplementary material

**Supplementary Figure 1.**
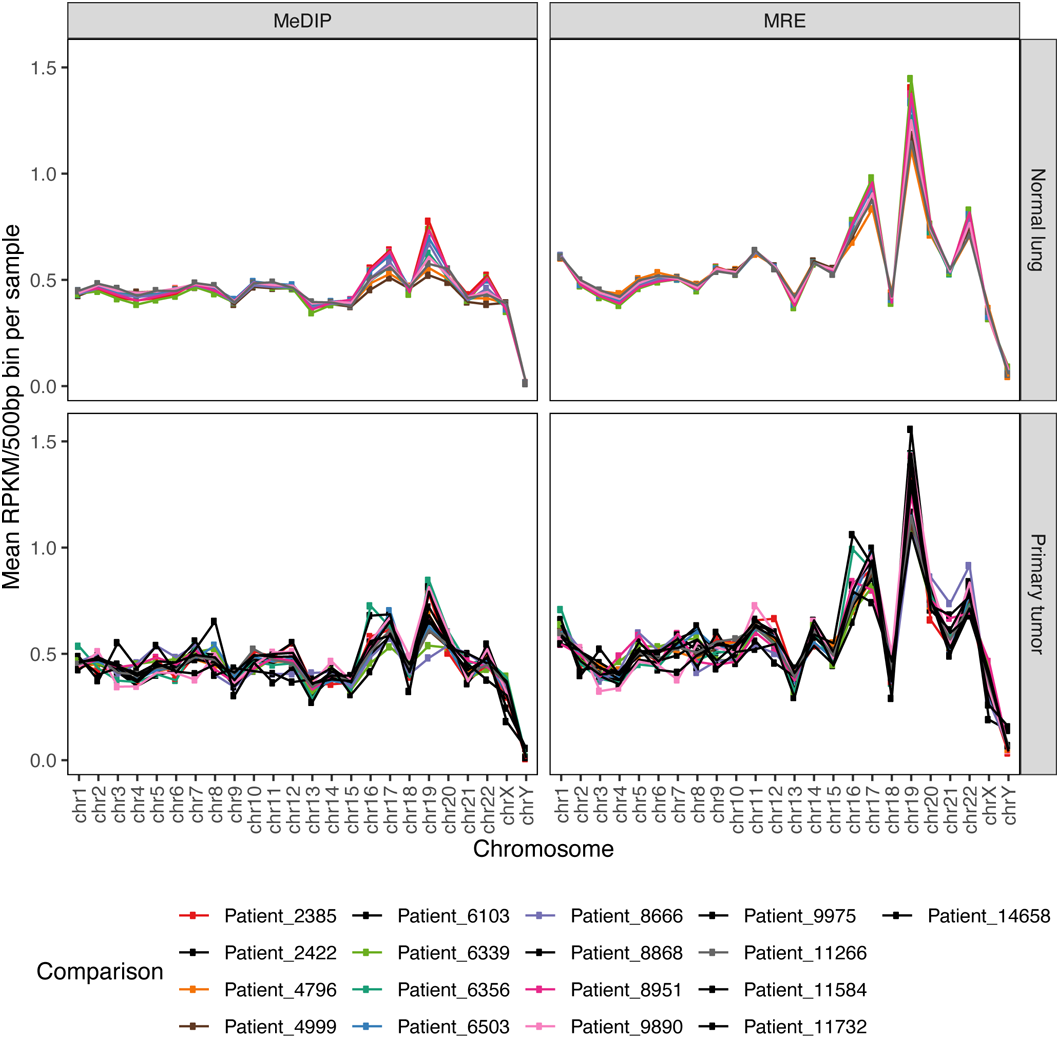
Read coverage over chromosomes. Mean MeDIP-seq and MRE-seq RPKM per 500 bp bin by sample for each chromosome, split by sample malignancy.

**Supplementary Figure 2.**
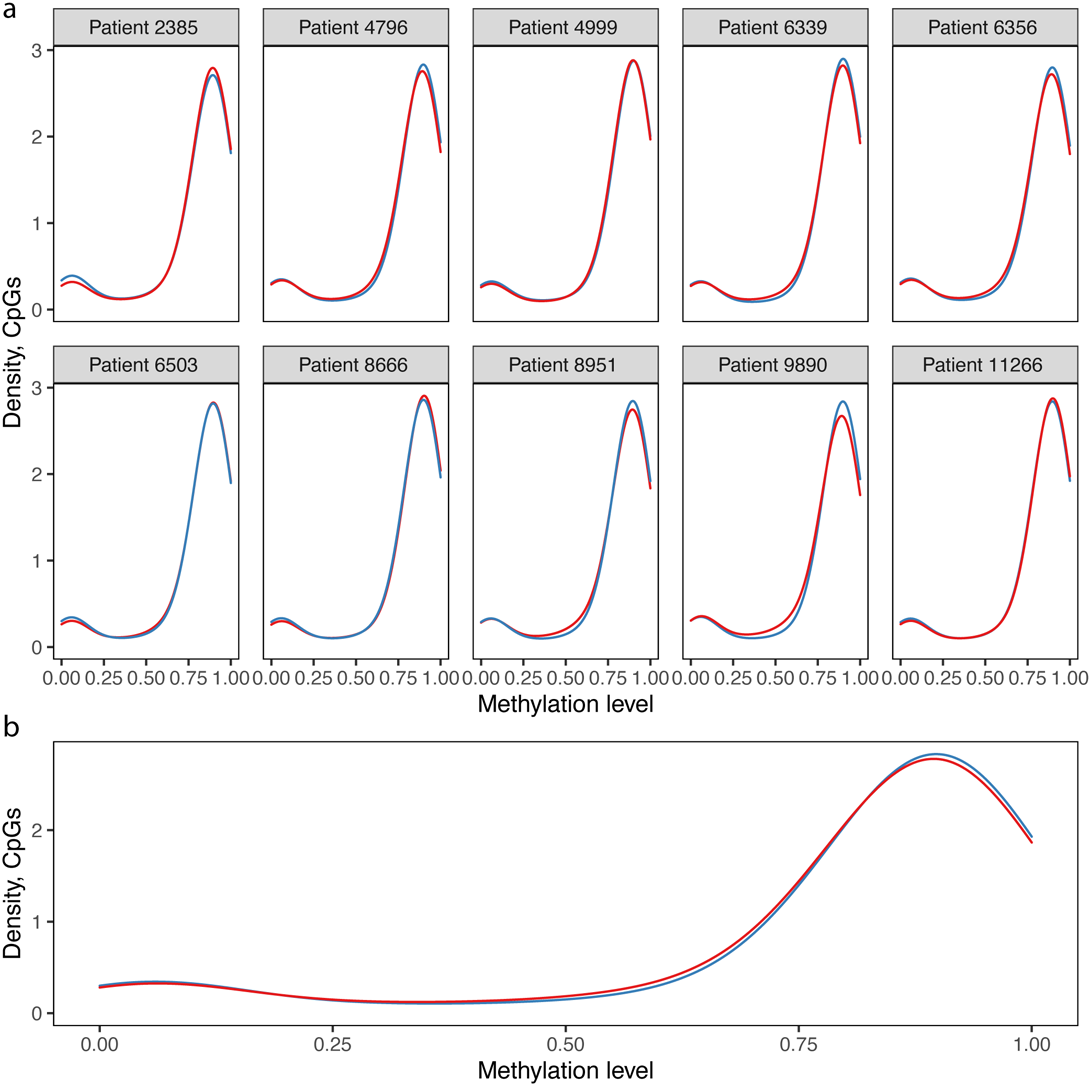
CpG methylation level distribution by sample and patient and averaged across disease state. **A.** Distribution of the number of CpGs at each methylation level in the normal lung and primary tumor samples for each patient. Samples are colored by malignancy, where blue is normal and red is tumor. **B.** Distribution of the number of CpGs at each methylation level as assigned by methylCRF, averaged over all samples by malignancy, where blue is normal and red is tumor.

**Supplementary Figure 3.**
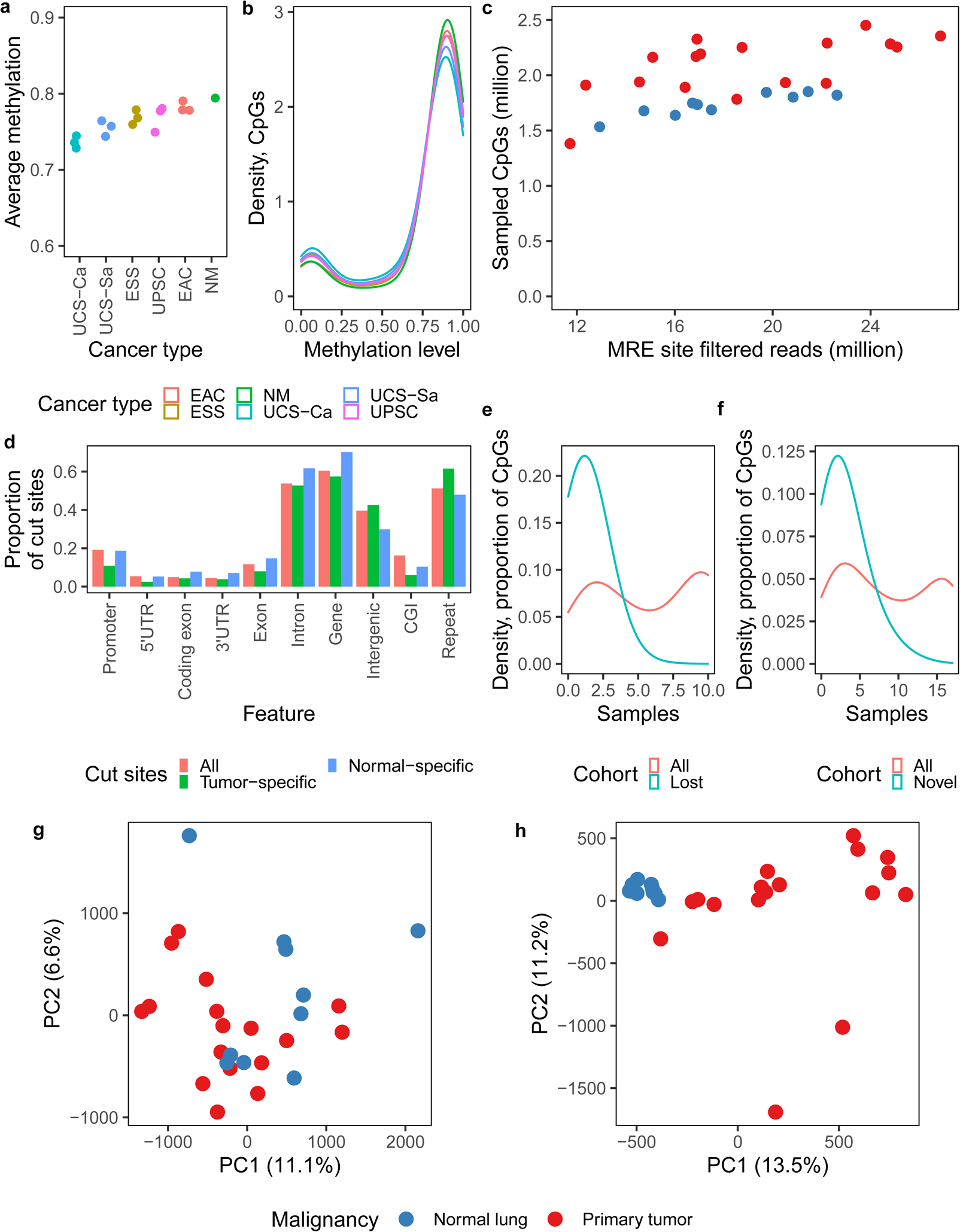
DNA methylation alterations using MRE-seq and MeDIP-seq data and comparison to previously published samples. **A-B.** Samples from [63], including a pooled normal endometrium sample (NM, 10 pooled samples), uterine carcinosarcoma carcinoma (UCS-Ca) and sarcoma (UCS-Sa) components, endometrioid carcinoma (EAC), endometrial stromal sarcoma (ESS), and endometrial serous carcinoma (UPSC). **A.** Average genome-wide CpG methylation level per sample, ordered by mean methylation level per cancer type. **B.** Distribution of the number of CpGs at each methylation level as assigned by methylCRF, averaged over all samples by cancer type. **C.** Number of sampled MRE CpG cut sites versus the number of MRE-seq reads mapped to cut sites per sample, colored by sample malignancy (see legend below Figure S3g-h, Wilcox test *p*-value when comparing normal vs. tumor sampled sites: *p* < 0.001). **D.** Location of MRE CpG cut sites, where “All” is all possible MRE cut sites. **E.** Distribution of the proportion of MRE CpG cut sites sampled in each number of normal samples, where “All” is all MRE cut sites sampled in normal samples and “Lost” is MRE cut sites sampled only in normal samples. **F.** Distribution of the proportion of MRE CpG cut sites sampled in each number of tumors, where “All” is all MRE cut sites sampled in tumors and “Novel” is MRE cut sites sampled only in tumors. **G.** PCA on MeDIP-seq RPKM per 500 bp bin, colored by sample malignancy. **H.** PCA on MRE-seq RPKM per 500 bp bin, colored by sample malignancy.

**Supplementary Figure 4.**
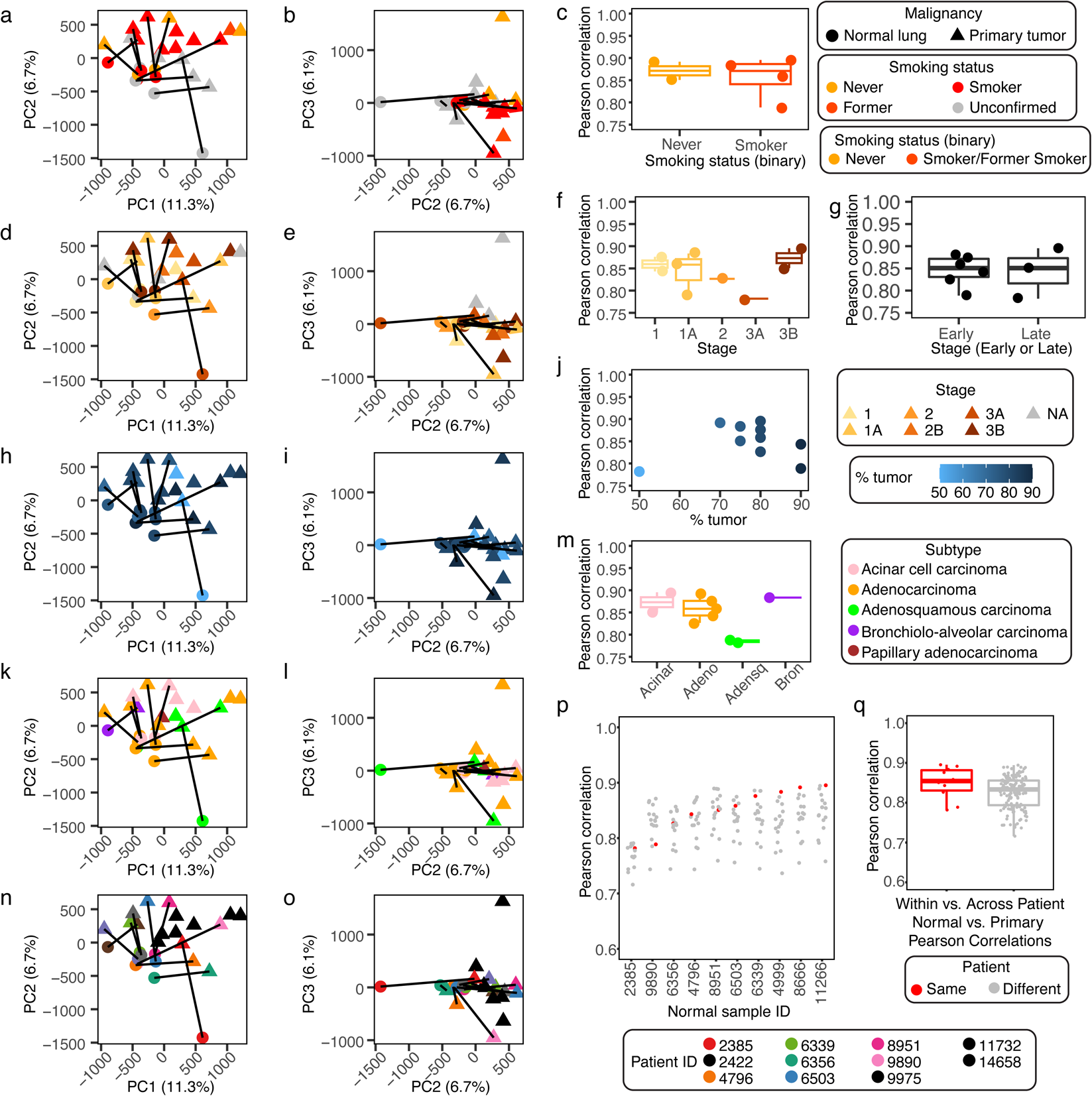
Global patient methylation changes by patient clinicopathologic data. **A,B,D,E,H,I,K,L,N,O.** PCA on normal lung and primary tumor samples, using the mean methylation over genome-wide 1 kb windows as features. Sample malignancy is indicated by shape (legend at top right). Axis titles display the amount of variance explained by each PC. Normal and primary tumor samples from the same patient are connected with a line. PCA plots are colored by **(A-B)** smoking status (legend to the right of panel c), **(D-E)** tumor stage (legend to the right of panel **J**), **(H-I)** percent tumor (legend to the right of panel **J**), **(K-L)** subtype (legend to the right of panel **M**), or **(N-O)** patient ID (legend below panel **P**). P2385_N_UC is the outlier on PC2, and P14658_T_NS is the outlier for PC3. **C,F,G,J,M.** Pearson correlation between paired normal and primary tumors, separated by **c**) smoking status (see binary legend, Wilcox test *p*-value = 1), **(F)** tumor stage, **(G)** binary tumor stage (early: stages 1, 1A, 2; late: stages 3A, 3B) (Wilcox test *p*-value = 1), **(J)** percent tumor purity (Pearson product moment correlation *p*-value = 0.6079), or **(M)** subtype (Wilcox test comparing adenocarcinomas to adenosquamous carcinomas *p*-value = 0.09524). **P.** Pearson correlation between each normal lung sample and all tumors over genome-wide 1 kb windows, colored by whether the tumor is from the same patient and ordered by increasing correlation with the tumor from the same patient. **Q.** Pearson correlation between each normal lung sample and all tumors over genome-wide 1 kb windows, grouped according to paired or unpaired comparisons (Wilcox test *p*-value = 0.06136). Acinar: acinar cell carcinoma, Adeno: adenocarcinoma, Adensq: adenosquamous carcinoma, Bron: bronchiolo-alveolar carcinoma, Unc.: Unconfirmed.

**Supplementary Figure 5.**
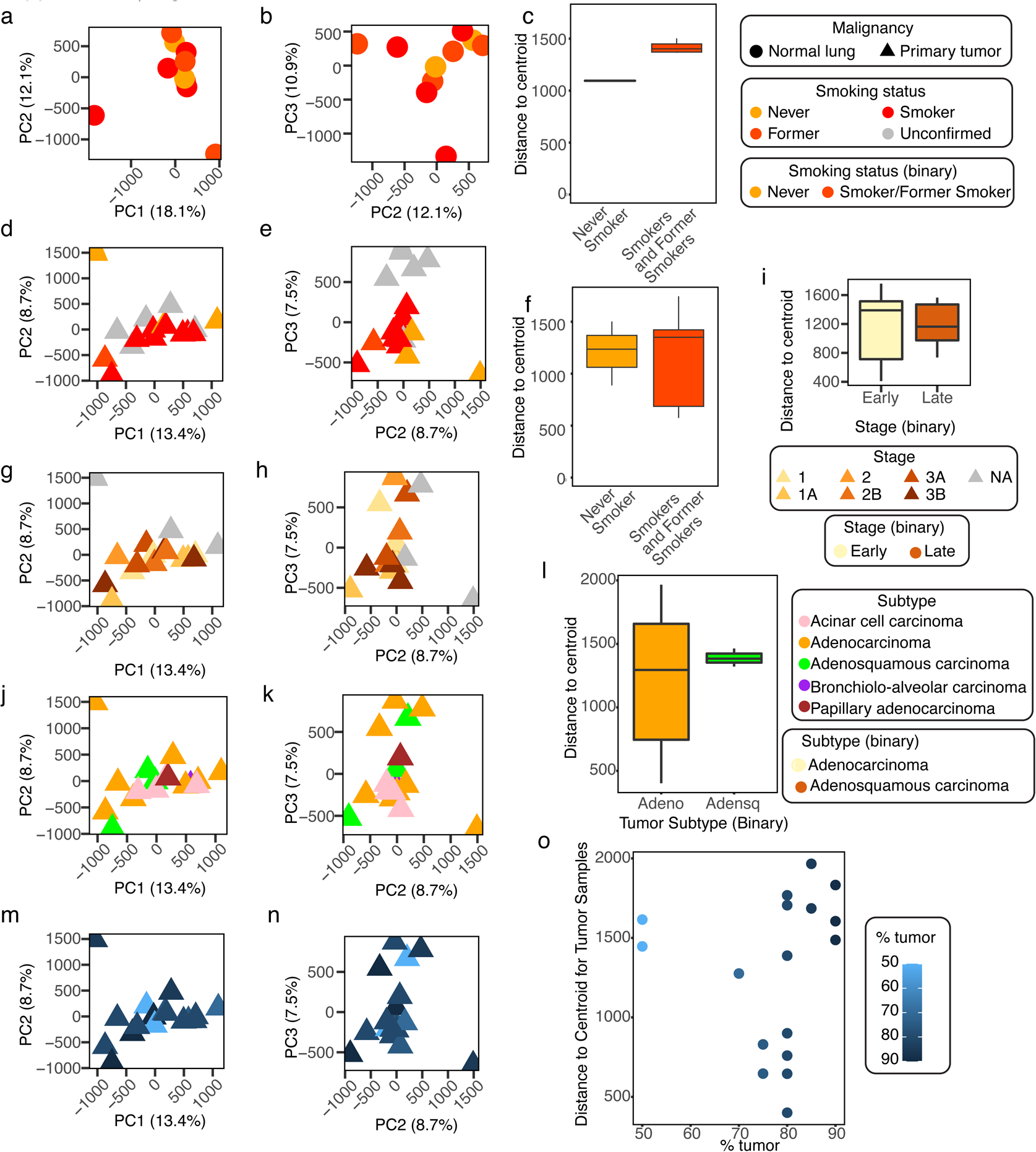
CpG methylation variation by patient clinicopathologic data. **A-B.** PCA on normal lung samples, using the mean methylation over genome-wide 1 kb windows as features, colored by smoking status (legend to the right of panel **C**). The outliers along PC1 are P2385_N_UC (left) and P4999_N_S (right), and the outlier along PC3 is P6356_N_UC. **C.** Distance to centroid for normal samples from smokers and from never smokers (Wilcox test *p*-value = 0.1002). **D,E,G,H,J,K,M,N.** PCA on tumor samples, using the mean methylation over genome-wide 1 kb windows as features, colored by **(D-E)** smoking status, **(G-H)** tumor stage (legend below panel **I**), **(J-K)** subtype (legend to the right of panel **L**), or **(M-N)** percent tumor purity (legend to the right of panel **O**). **F.** Distance to centroid of tumor samples from smokers and never smokers (Wilcox test *p*-value = 0.8636). **I.** Distance to centroid of early stage (1, 1A, 2, 2B) and late stage (3A, 3B) tumors (Wilcox test *p*-value = 0.8981) **L.** Distance to centroid of adenocarcinoma and adenoquamous carcinoma tumors (Wilcox test *p*-value = 0.7676). **O.** Distance to centroid of tumor samples compared to percent tumor purity (Pearson product moment correlation *p*-value = 0.7664). Adeno: adenocarcinoma, Adensq: adenosquamous carcinoma.

**Supplementary Figure 6.**
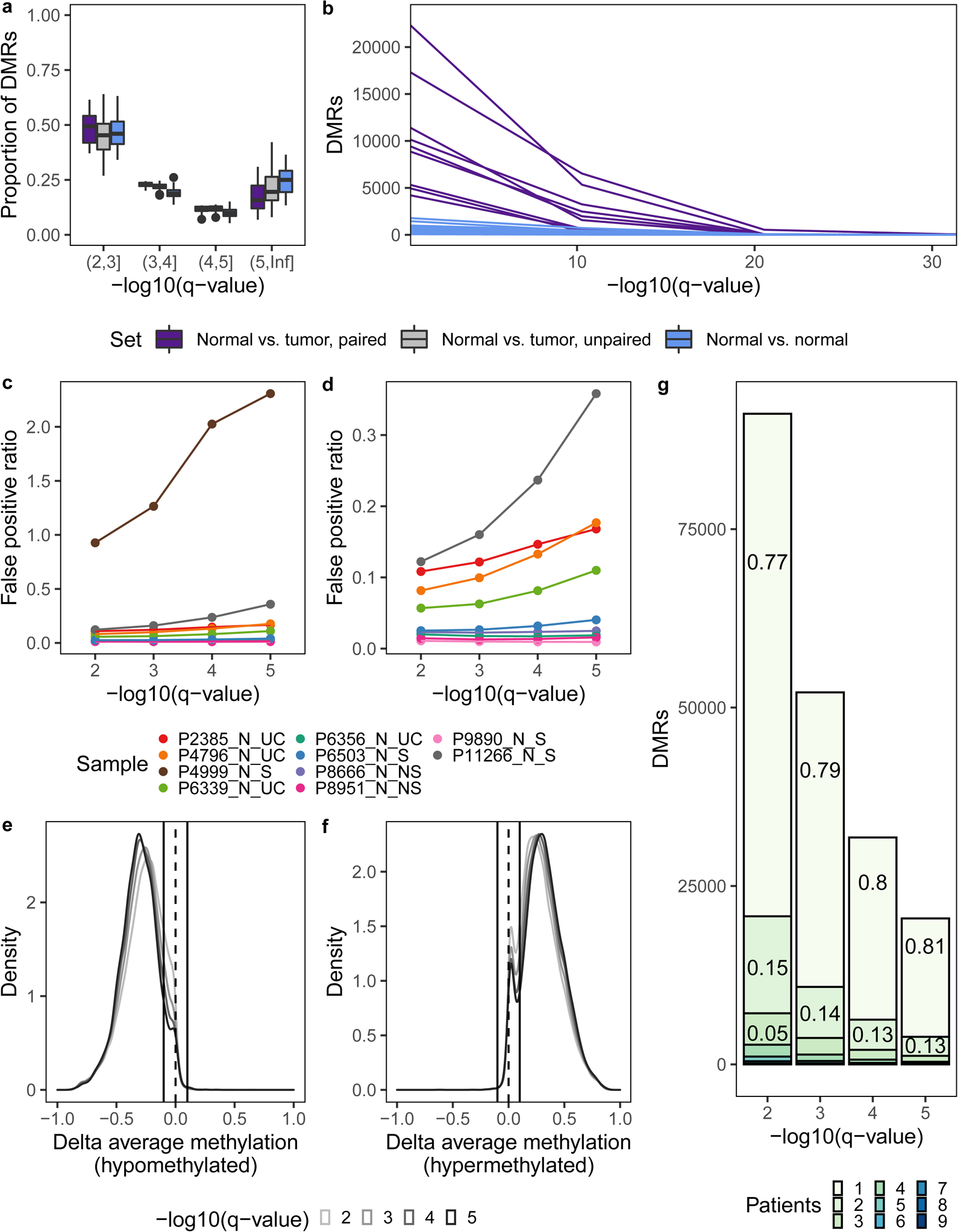
Selection of DMR *q*-value threshold. **A.** Proportion of DMRs within each *q*-value interval per comparison, by comparison set. **B.** Distribution of the number of DMRs by *q*-value for each comparison, colored by comparison set, patient-matched normal vs. tumor and normal vs. normal lung comparisons only. A pseudocount of 1e-300 was added to the *q*-value to avoid undefined values, and DMRs with *q*-value < 1e-30 are not shown (*n* = 1,433). **C-D.** False positive ratio (mean number of DMRs in comparison to other normal samples divided by the number of DMRs in comparison to patient-matched tumor) for each normal sample for DMRs below each *q*-value threshold, for **(C)** all samples or **(D)** excluding Patient 4999. **E-F.** Distribution of the change in mean CpG methylation level (as assigned by methylCRF) over each DMR between the patient-matched normal and tumor sample(s) in which the DMR was identified, for DMRs below four *q*-value thresholds. Dashed line indicates no change in methylation; solid lines indicate a methylation change of 10%. **G.** Proportion of DMRs below each *q*-value threshold that are shared by each number of patients. Proportions < 0.05 are not shown.

**Supplementary Figure 7.**
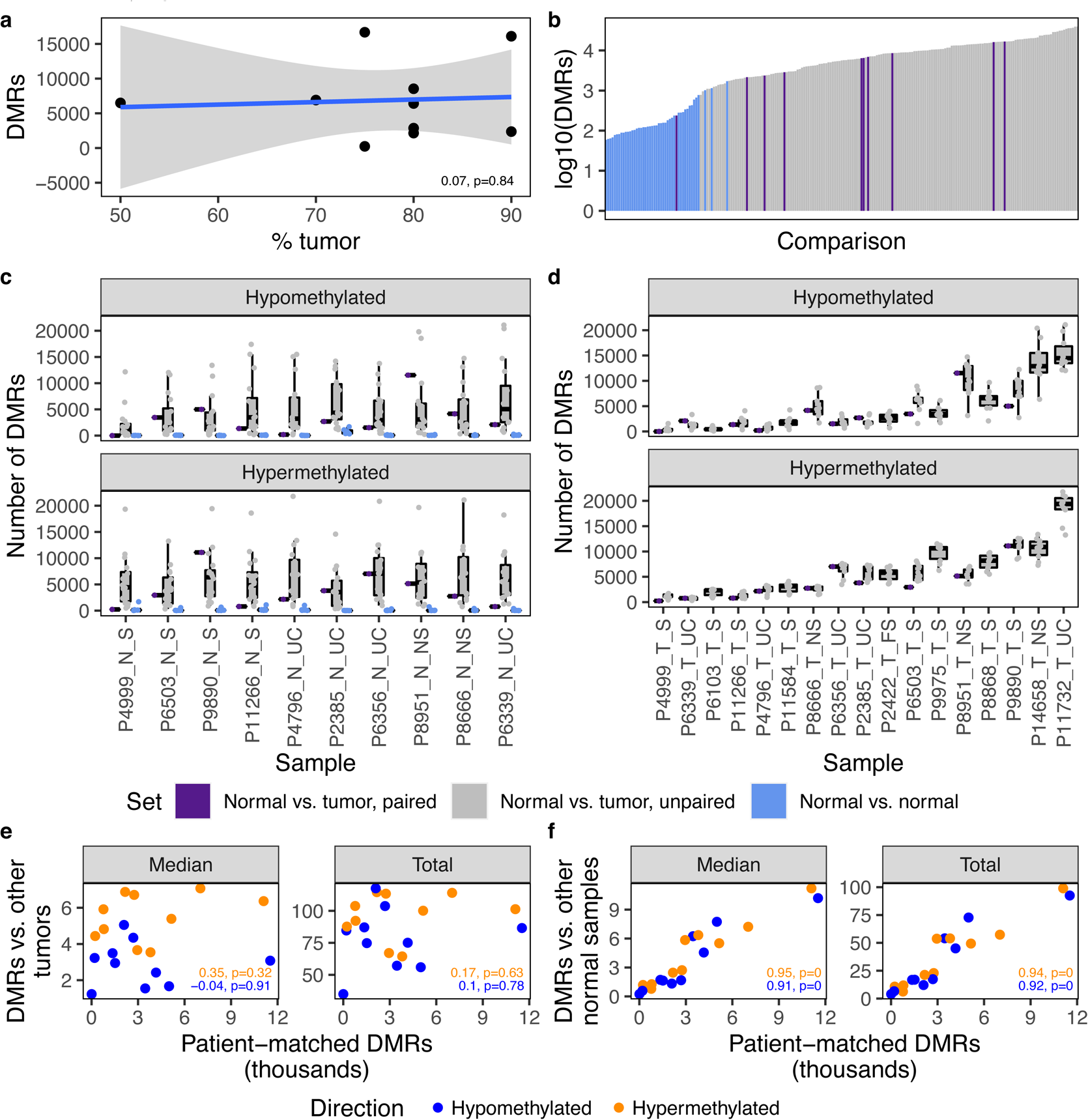
Number of DMRs per comparison by normal and tumor sample. **A.** Number of patient DMRs as a function of percent tumor. The linear relationship with shaded 95% confidence intervals is plotted. **B.** Number of DMRs per comparison (*x*-axis), colored by comparison set and ordered by increasing number of DMRs. **C-D.** Number of DMRs identified between **(C)** each normal sample and all other samples and **(D)** each tumor sample and all other samples, colored by comparison set, split by DMR direction. Samples are ordered by median number of DMRs across all comparisons. Normal vs. normal DMRs are counted for each normal sample. **E-F.** Number of DMRs between patient-matched normal and tumor samples (*x*-axis) versus the number of DMRs between **(E)** the normal sample and other tumors and **(F)** the tumor sample and other normal samples. Both the median and total number of DMRs in comparison to other samples are presented, and DMRs are split by direction. The Pearson correlation and *p*-value by DMR direction are listed on each graph.

**Supplementary Figure 8.**
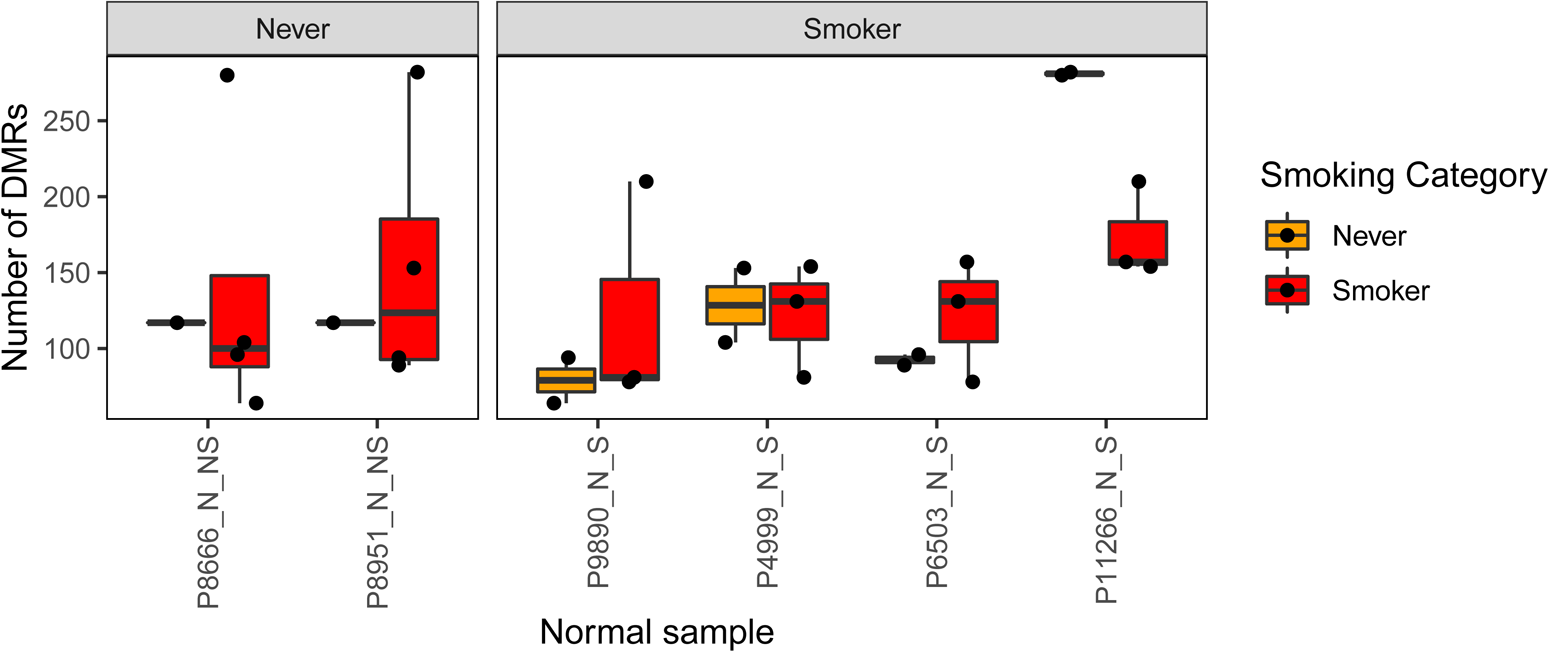
Field effects of smoking. Number of DMRs between normal lung samples. Normal samples are split into never-smokers and smokers by facets, and boxplot color indicates the smoking status of the second normal sample in the pair. Wilcox *p*-values > 0.1 for each normal sample.

**Supplementary Figure 9.**
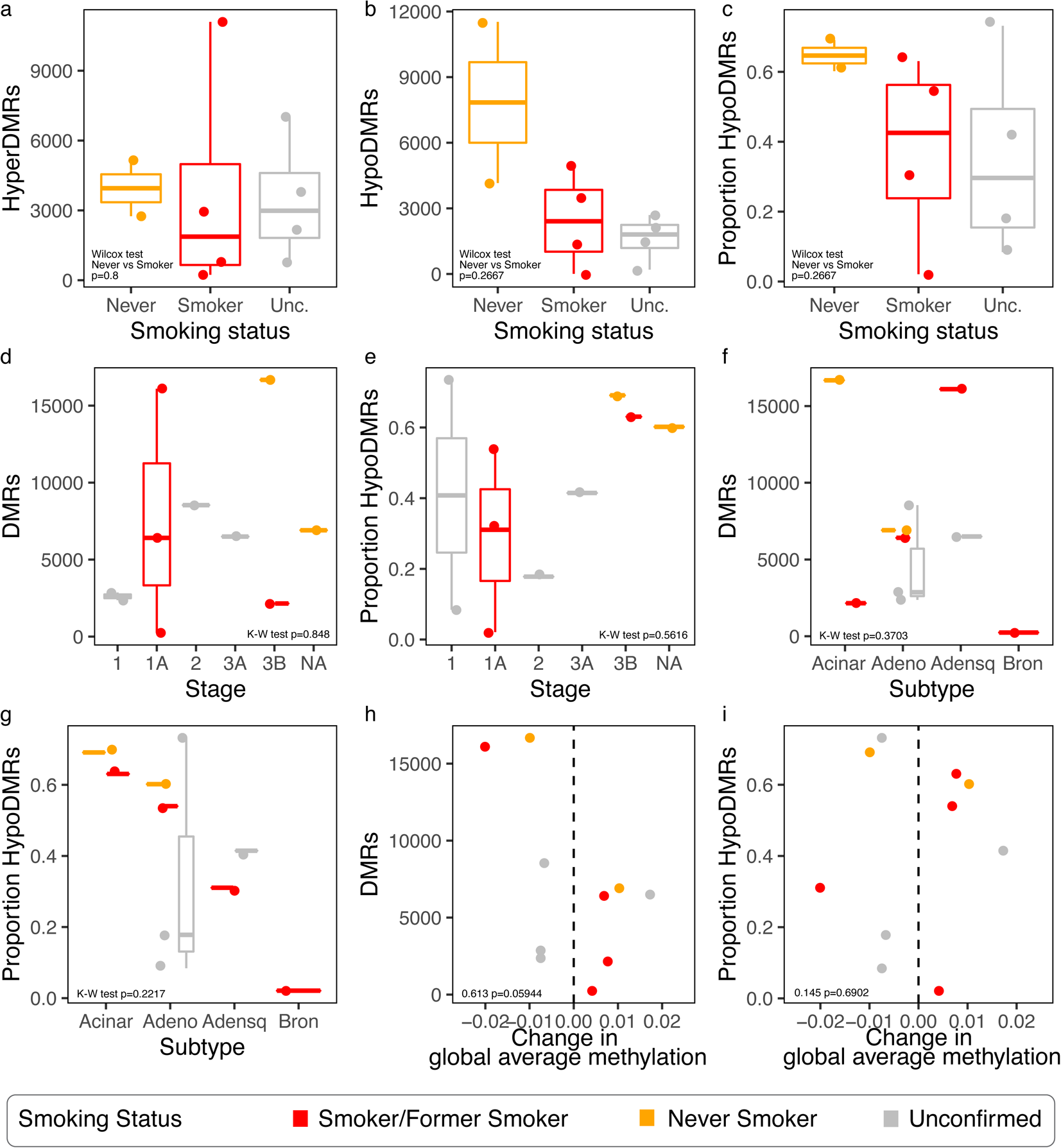
Number of DMRs and proportion hypomethylated by patient clinicopathologic data. **A-B.** Comparison of the number of **(A)** hyperDMRs and **(B)** and hypoDMRs for each patient according to smoking status. Wilcox tests comparing never-smokers (*n* = 2) to smokers (*n* = 4) were not significant for hyperDMRs (*p* = 0.8) or hypoDMRs (*p* = 0.2667). **C.** Comparison of the proportion of hypoDMRs by smoking status. Wilcox test comparing never-smokers (*n* = 2) to smokers (*n* = 4) was not significant (*p* = 0.2667). **D.** Comparison of the number of DMRs across patients according to tumor stage (*n* = 2,3,1,1,2,1), colored by smoking status. Kruskal-Wallis *p* = 0.848. **E.** Comparison of the proportion of hypoDMRs across patients according to tumor stage (*n* = 2,3,1,1,2,1), colored by smoking status. Kruskal-Wallis *p* = 0.5616. **F.** Comparison of the number of DMRs across patients according to tumor subtype (*n* = 2,5,2,1) colored by smoking status. Kruskal-Wallis *p* = 0.3703. **G.** Comparison of the proportion of hypoDMRs across patients according to tumor subtype (*n* = 2,5,2,1), colored by smoking status. Kruskal-Wallis *p* = 0.2217. **H.** Comparison of the number of DMRs across patients according to the change in genome-wide average CpG methylation level between normal lung and tumor sample, colored by smoking status. Dashed line indicates no change in methylation. Pearson’s product-moment correlation between number of DMRs and absolute change in methylation: 0.613, *p* = 0.05944. **I.** Comparison of the proportion of hypoDMRs across patients according to the change in genome-wide average CpG methylation level between normal lung and tumor sample, colored by smoking status. Pearson’s product-moment correlation between proportion of hypoDMRs and change in methylation: 0.105, *p* = 0.7718. Acinar: acinar cell carcinoma, Adeno: adenocarcinoma, Adenosq: adenosquamous carcinoma, Bron: bronchiolo-alveolar carcinoma.

**Supplementary Figure 10.**
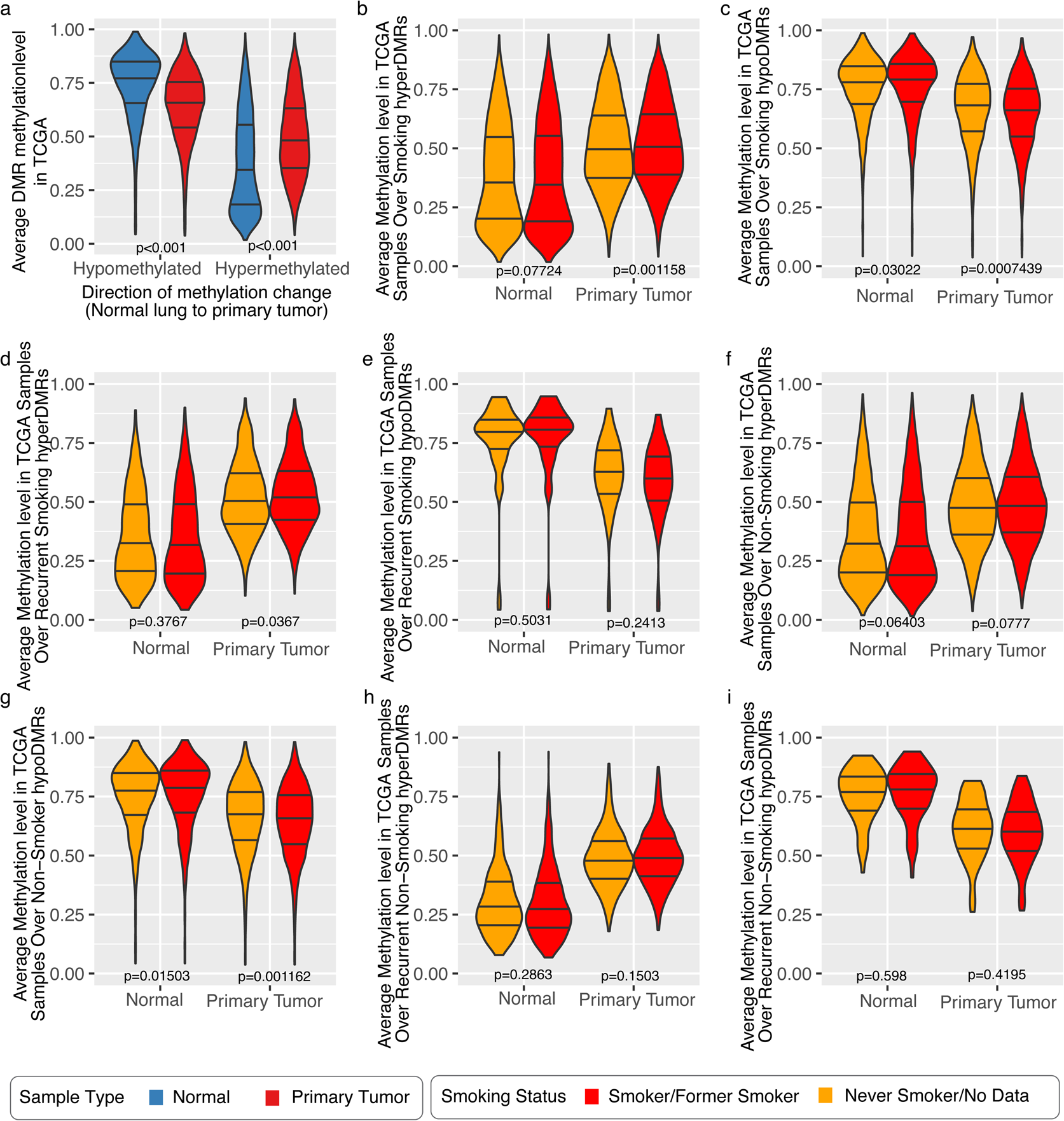
DMR methylation status across TCGA LUAD samples. **A.** Mean methylation level of each hypo- or hypermethylated DMR across all TCGA LUAD samples, split by sample type. Violin plot lines indicate quartiles. Wilcox *p* < 0.001 between LUAD normal lung (blue) and primary tumor (red) for both hypo- and hypermethylated DMRs. **B-I.** Average methylation in TCGA samples, separated by sample type and smoking status, over **(B)** hyperDMRs identified in smoking patients from our cohort (*n* = 7410), **(C)** hypoDMRs identified in smoking patients from our cohort (*n* = 1137), **(D)** hyperDMRs identified in at least two smoking patients from our cohort (*n* = 967), **(E)** hypoDMRs identified in at least two smoking patients from our cohort (*n* = 55), **(F)** hyperDMRs identified in never-smoker patients from our cohort (*n* = 4063), **(G)** hypoDMRs identified in never-smoker patients from our cohort (*n* = 2014), **(H)** hyperDMRs identified in both never-smoker patients from our cohort (*n* = 501), or **(I)** hypoDMRs identified in both never-smoker patients from our cohort (*n* = 74). *p*-values from Wilcox tests comparing TCGA normal samples from smokers and never-smokers and comparing TCGA primary tumor samples from smokers and never-smokers are indicated.

**Supplementary Figure 11.**
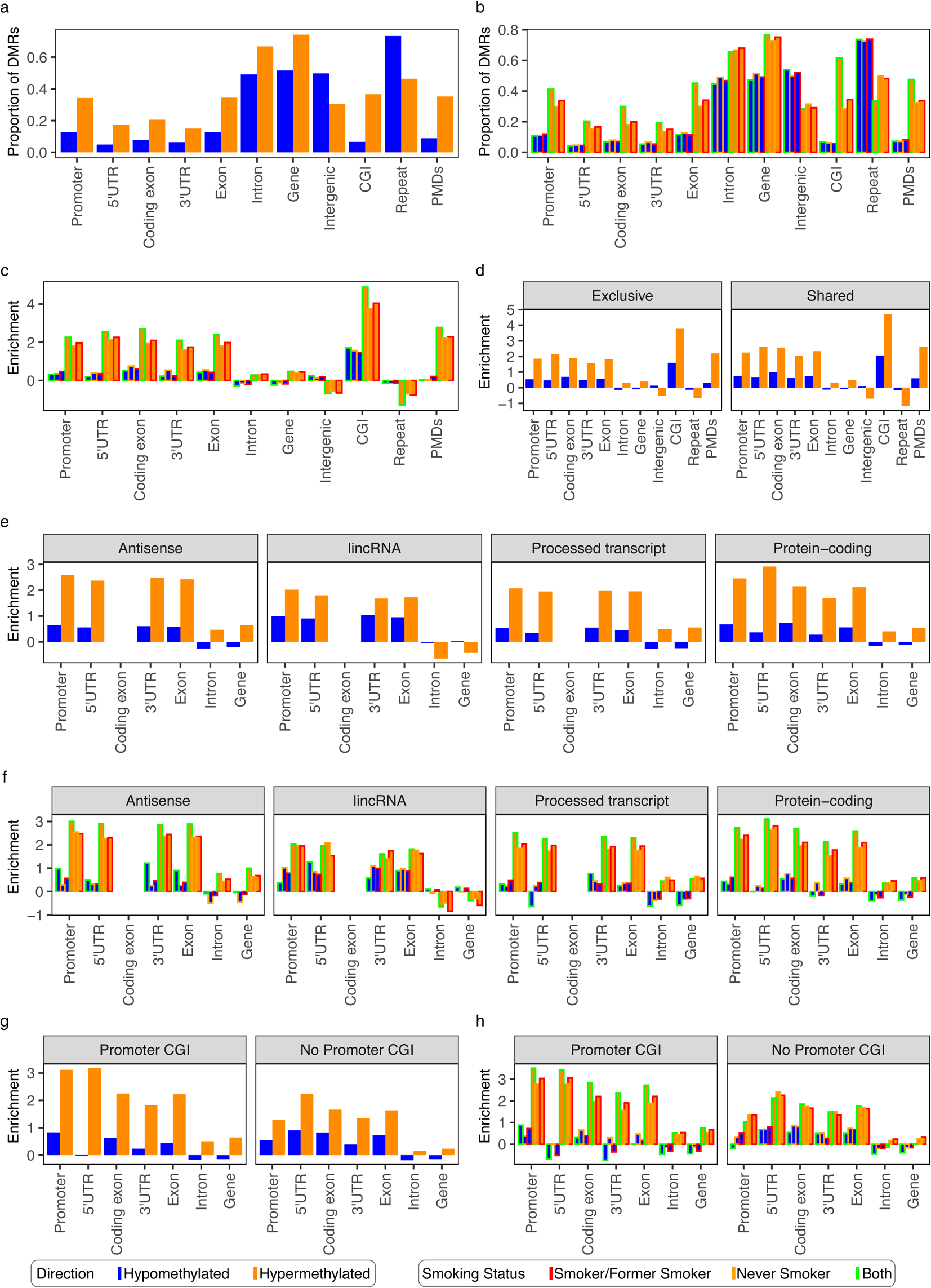
Genomic location of DMRs by frequency and transcript information. **A-B.** Proportion of patient-matched DMRs over genic features, intergenic regions, CpG islands (CGI), and repeats, by **(A)** DMR direction, and **(B)** smoking status. **C.** Enrichment of patient-matched DMRs over genic features, intergenic regions, CpG islands (CGI), and repeats, by DMR direction and smoking status. **D-H.** Log odds ratio enrichment of patient-matched DMRs over genic features, intergenic regions, CpG islands (CGI), and repeats compared to the background distribution of 500 bp bins containing CpGs, by DMR direction **(D,E,G)** and smoking status **(F,H)**. **D.** DMRs exclusive to a single patient or shared between multiple patients. **E-F.** Genic features restricted to four major categories of transcript biotype: protein-coding (*n* = 85,756, 45% of transcripts), processed transcript (*n* = 29,941, 16%), lincRNA (*n* = 11,773, 6%), and antisense (*n* = 9,713, 5%). **G-H.** Protein-coding genic features split by whether the transcript promoter overlaps a CpG island (yes: *n* = 52,222 transcripts, 64%; no: *n* = 29,357, 36%).

**Supplementary Figure 12.**
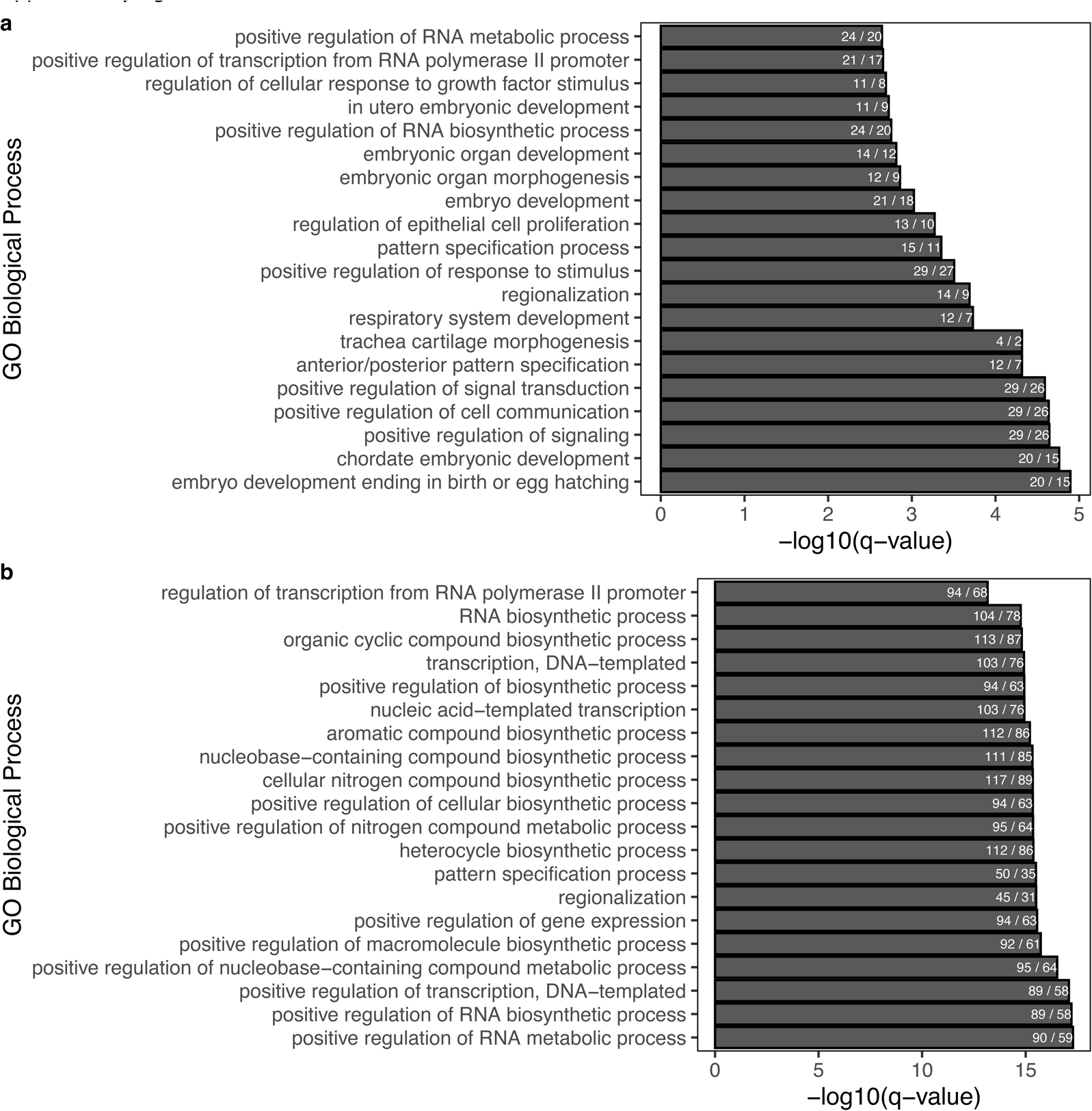
Smoking status-specific DMR enrichment. **A-B.** Top 20 significant GO Biological Processes as identified by GREAT (see Materials and methods) for **(A)** hyperDMRs identified in at least 3 smokers and not in either never-smoker (of 40 terms), **(B)** hyperDMRs identified in both never-smokers and not in any smoker (of 200 terms). Terms are ordered by FDR-corrected binomial *q*-value and are labelled by the number of DMRs over the number of genes involved.

**Supplementary Figure 13.**
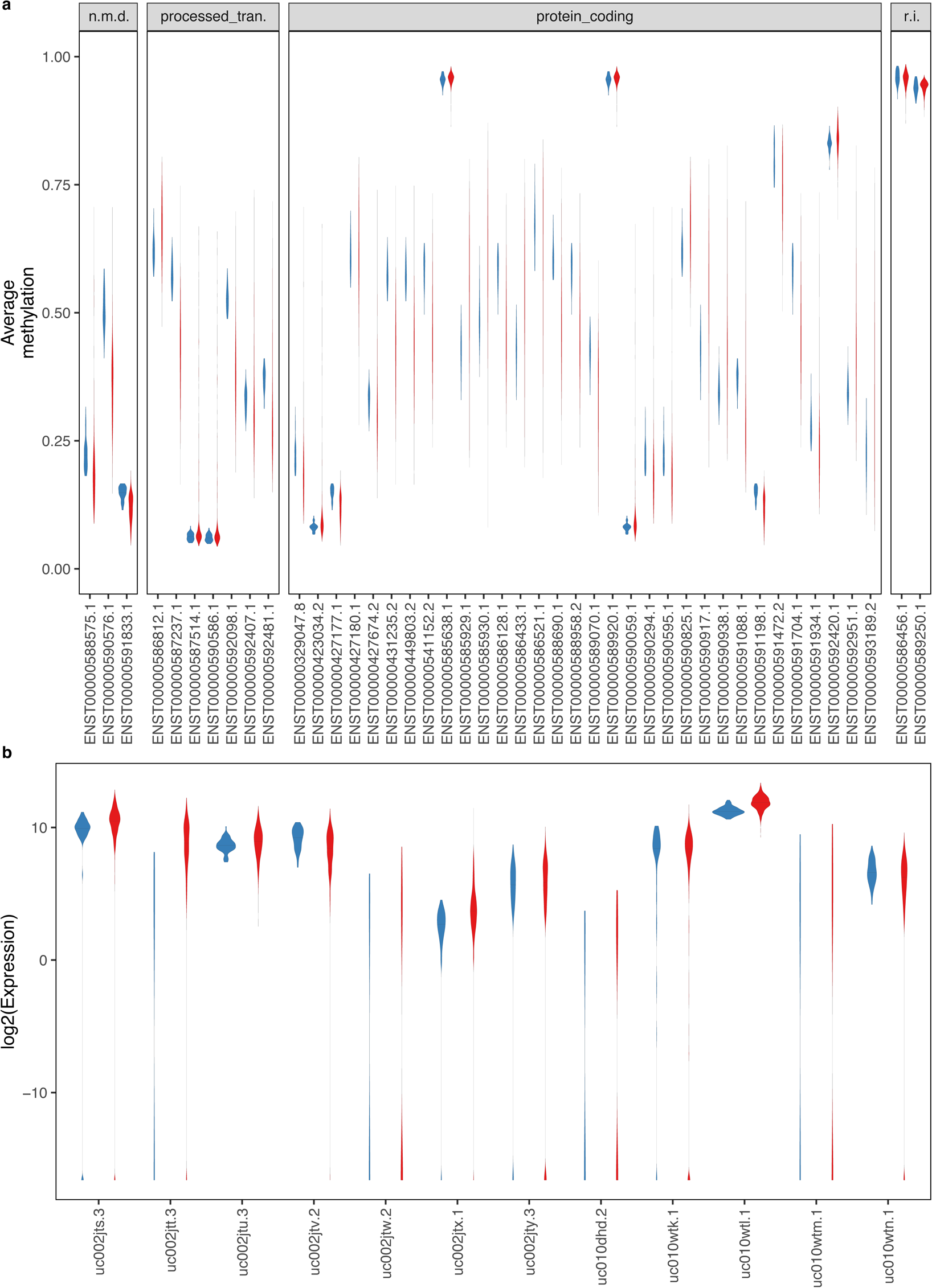
*SEPT9* isoform promoter methylation and expression in TCGA. **A.** Average *SEPT9* isoform promoter methylation levels in TCGA LUAD samples. n.m.d, nonsense_mediated_decay; processed_tran., processed_transcript; r.i., retained_intron. A pseudocount of 0.00001 was added to each value. Lines represent median values. **B.** Average expression of *SEPT9* isoforms in TCGA LUAD samples.

**Supplementary Figure 14.**
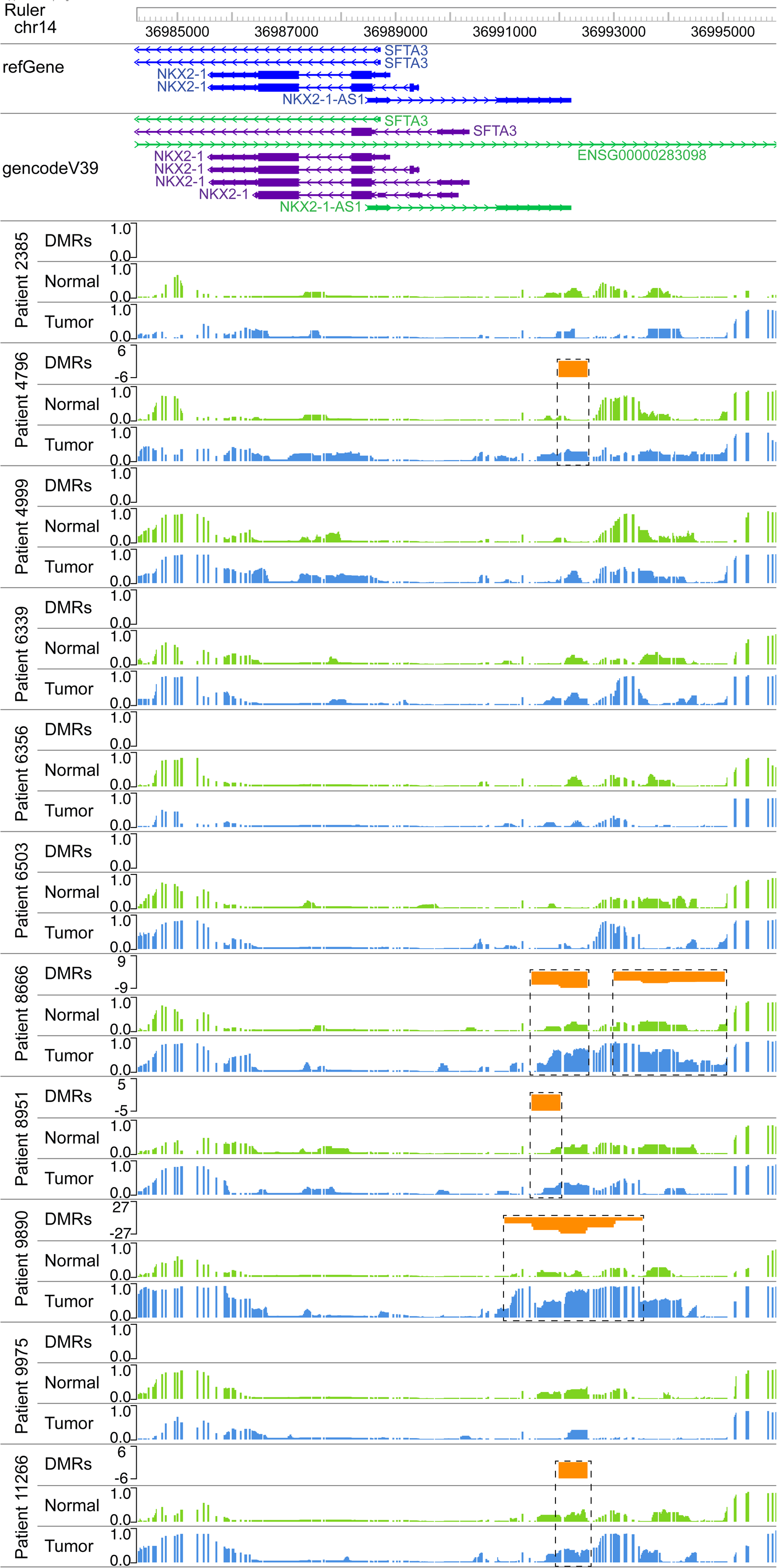

**Supplementary Figure 15.**
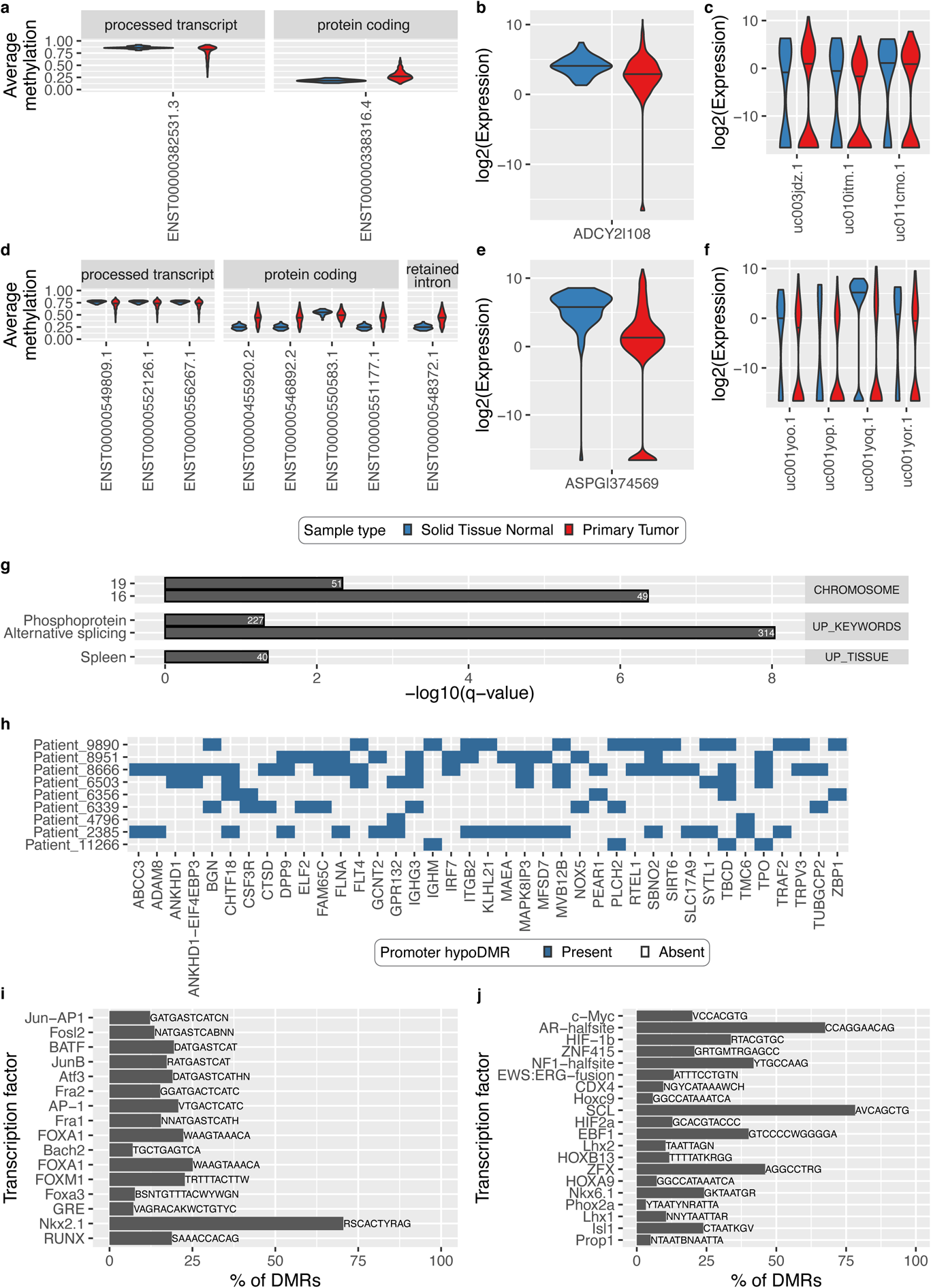
Enriched binding motifs and gene pathways for DMRs overlapping gene promoters. **A.** Average methylation of each *ADCY2* transcript promoter in TCGA LUAD samples and matched normal lung. **B.** *ADCY2* expression in TCGA samples. **C.** *ADCY2* isoform expression level in TCGA samples. **B-C.** A pseudocount of 0.00001 was added to each value. **D.** Average methylation of each *ASPG* transcript promoter in TCGA LUAD samples and matched normal lung. processed_tran., processed_transcript; r.t., retained_intron. **E.** *ASPG* expression in TCGA samples. **F.** *ASPG* isoform expression level in TCGA samples. **E-F.** A pseudocount of 0.00001 was added to each value. **A-F.** Lines represent median values. **G.** Significantly enriched gene sets among genes with a hypoDMR in the promoter in at least two patient comparisons, as determined by DAVID. Only terms with a Benjamini-corrected *p*-value < 0.05 are included. Terms are labelled with the number of selected genes and ordered by corrected *p*-value within each category. **H.** Indication of DAVID Spleen UP_TISSUE Pathway genes with a hypoDMR in the promoter in at least 2 patients, stratified by patient. **I-J.** Top 20 most enriched motifs within DMRs overlapping gene promoters in at least two patient comparisons, as determined by HOMER known motif analysis, for **(I)** hypoDMRs (*n* = 458) and **(J)** hyperDMRs (*n* = 3,050, out of 41 terms). Only motifs with a Benjamini-corrected *p*-value < 0.05 are included. Motifs are listed to the right of each bar, and the *y*-axis lists the corresponding transcription factor. Motifs are ordered by uncorrected *p*- value.

**Supplementary Figure 16.**
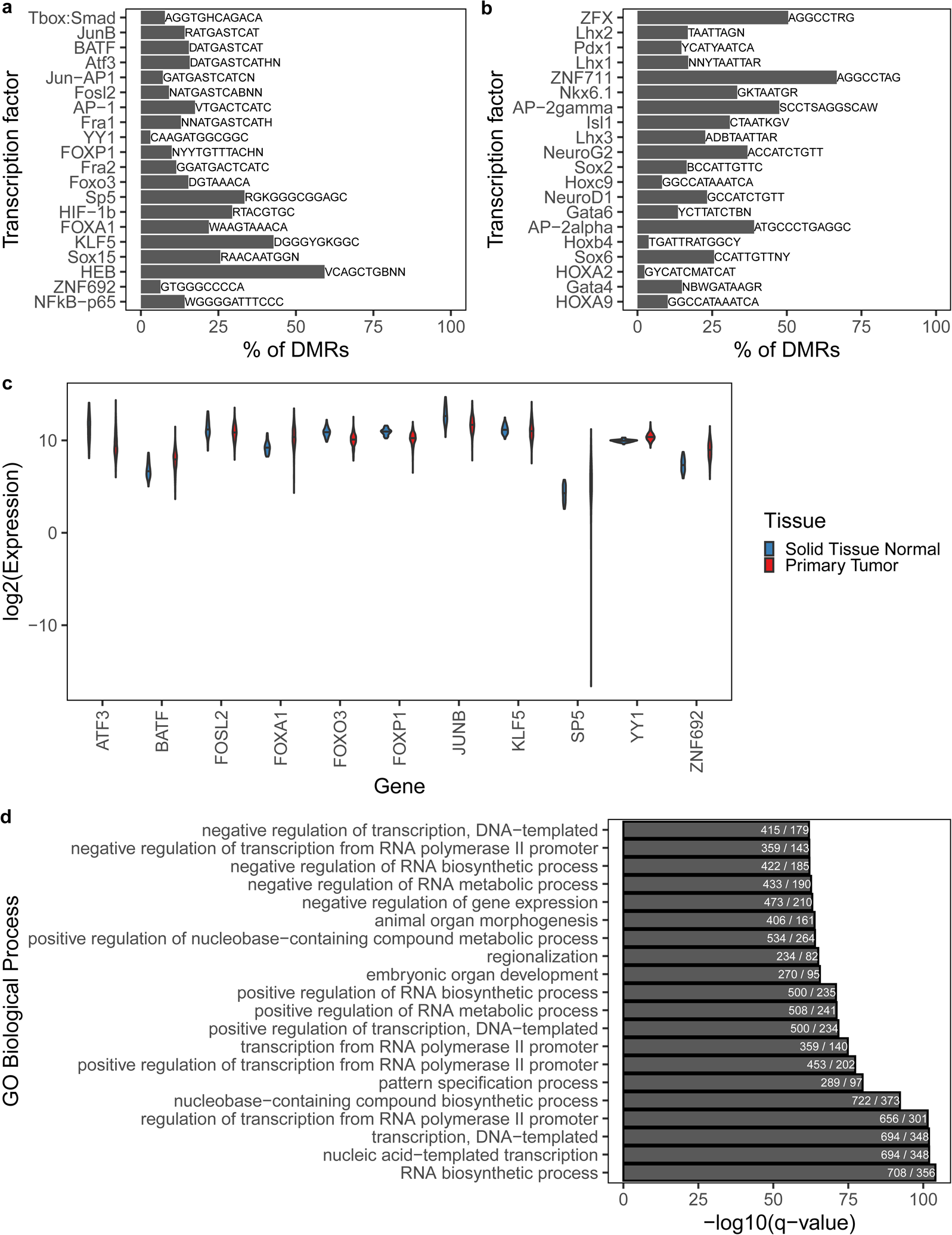
Enriched binding motifs and gene pathways for intergenic DMRs. **A-B.** Top 20 most enriched motifs within DMRs found in at least two patient comparisons that overlap intergenic regions but not genes or promoters (intergenic-exclusive), as determined by HOMER known motif analysis, for **(A)** hypoDMRs (*n* = 1,409 DMRs, 22 terms) and **(B)** hyperDMRs (*n* = 1,365 DMRs, 70 terms). Only motifs with a Benjamini-corrected *p*-value < 0.05 are included. Motifs are listed over each bar, and the *y*-axis lists the corresponding transcription factor. Motifs are ordered by uncorrected *p*-value. **C.** Expression level in TCGA LUAD samples of transcription factors with motifs enriched in intergenic hypoDMRs, split by sample type. Violin plot lines indicate median. A pseudocount of 0.00001 was added to avoid undefined values. Wilcox *p*-value < 0.001 for all but *KLF5*. **D.** Top 20 significant GO Biological Processes for intergenic-exclusive DMRs that are hypermethylated in at least two patients (out of 394 terms), as identified by GREAT (see Materials and methods). Terms are ordered by FDR-corrected binomial *q*-value and are labelled by the number of DMRs over the number of genes involved.

**Supplementary Figure 17.**
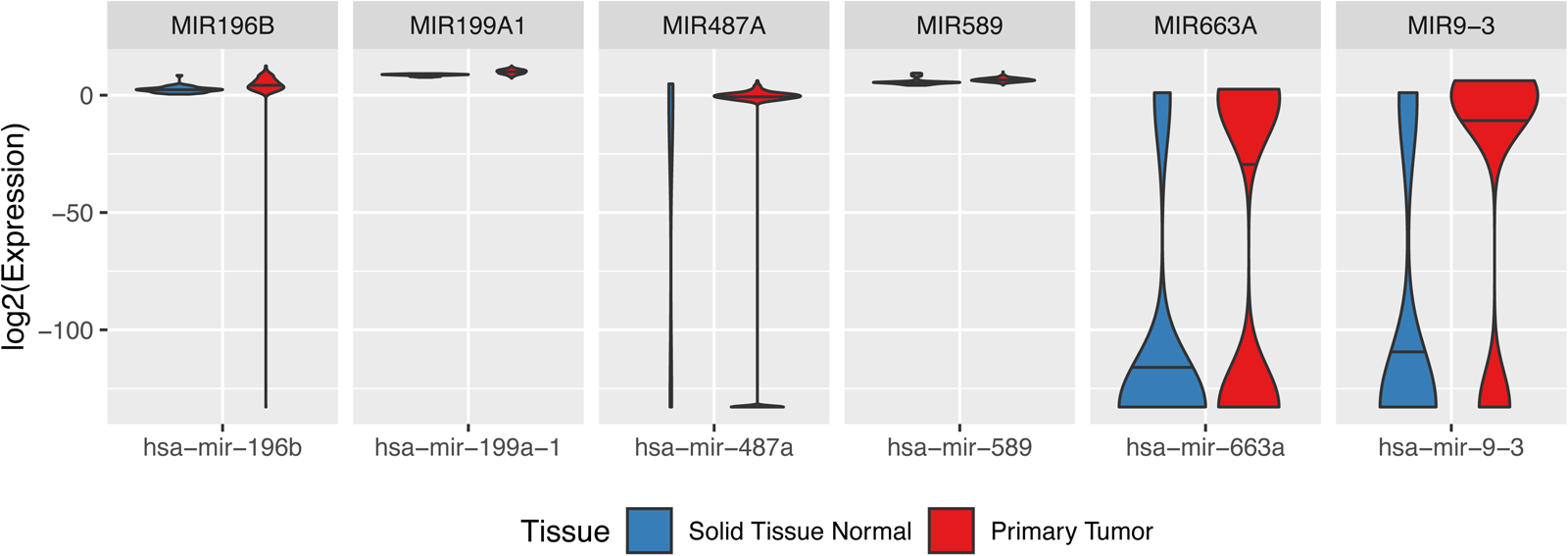
TCGA expression level of miRNA overlapping recurrent DMRs. miRNA expression level in TCGA LUAD and normal lung samples for miRNA that overlapped a DMR in multiple patients and had significantly different expression between TCGA sample types (Wilcox *p*-value < 0.01). A pseudocount of 1e-40 was added to avoid undefined values. Violin lines indicate median. The corresponding GENCODE gene name is listed above each graph. See Materials and methods for details on linking miRNA and GENCODE IDs.

**Supplementary Figure 18.**
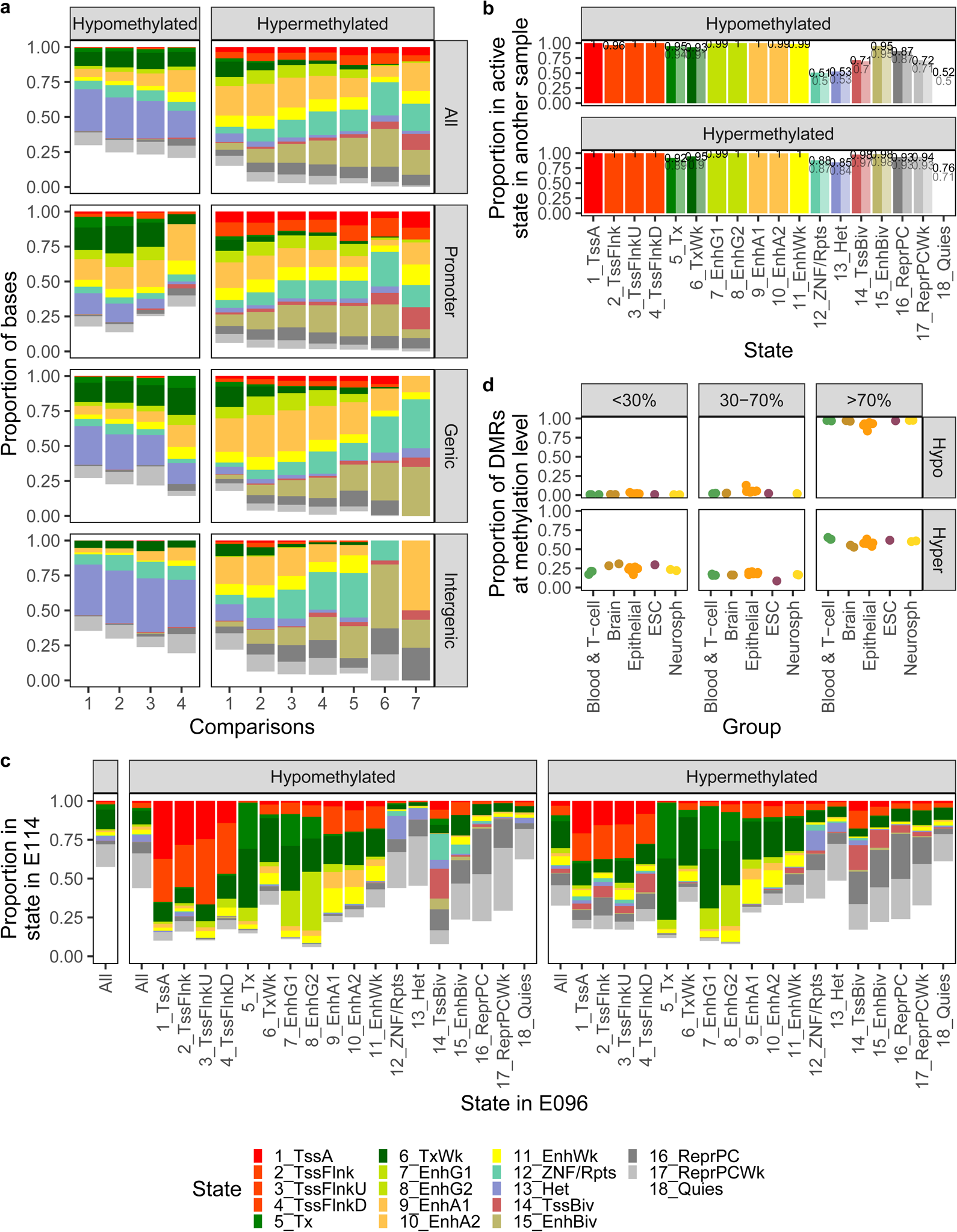
Epigenetic state of DMRs in Roadmap tissues. **A.** Proportion of Roadmap sample E096 (adult lung) bases overlapping hypo- or hyperDMRs in each 18-state chromHMM state, split by exclusive feature overlap and the number of patient comparisons in which the DMR is found (maximum 4 comparisons for hypoDMRs, 7 comparisons for hyperDMRs to avoid categories with < 10 DMRs). **B.** Proportion of DMRs in an active 15-state chromHMM state in a Roadmap sample besides E096, split by DMR direction, state in E096 (*x*-axis), and whether the DMR was in an active state in E096 (True: bold; False: faded). Proportions are listed above each bar. **C.** Proportion of Roadmap sample E114 (A549 lung carcinoma cell line) bases in each 18-state chromHMM state, overall or overlapping hypo- or hyperDMRs, and for DMRs split by chromHMM state in E096. The overall state in E114 was restricted to regions overlapping 500 bp bins that contained CpGs. **D.** Proportion of DMRs at each average methylation level in each Roadmap sample, split by sample group and DMR direction in the patient-matched NSCLC samples.

**Supplementary Figure 19.**
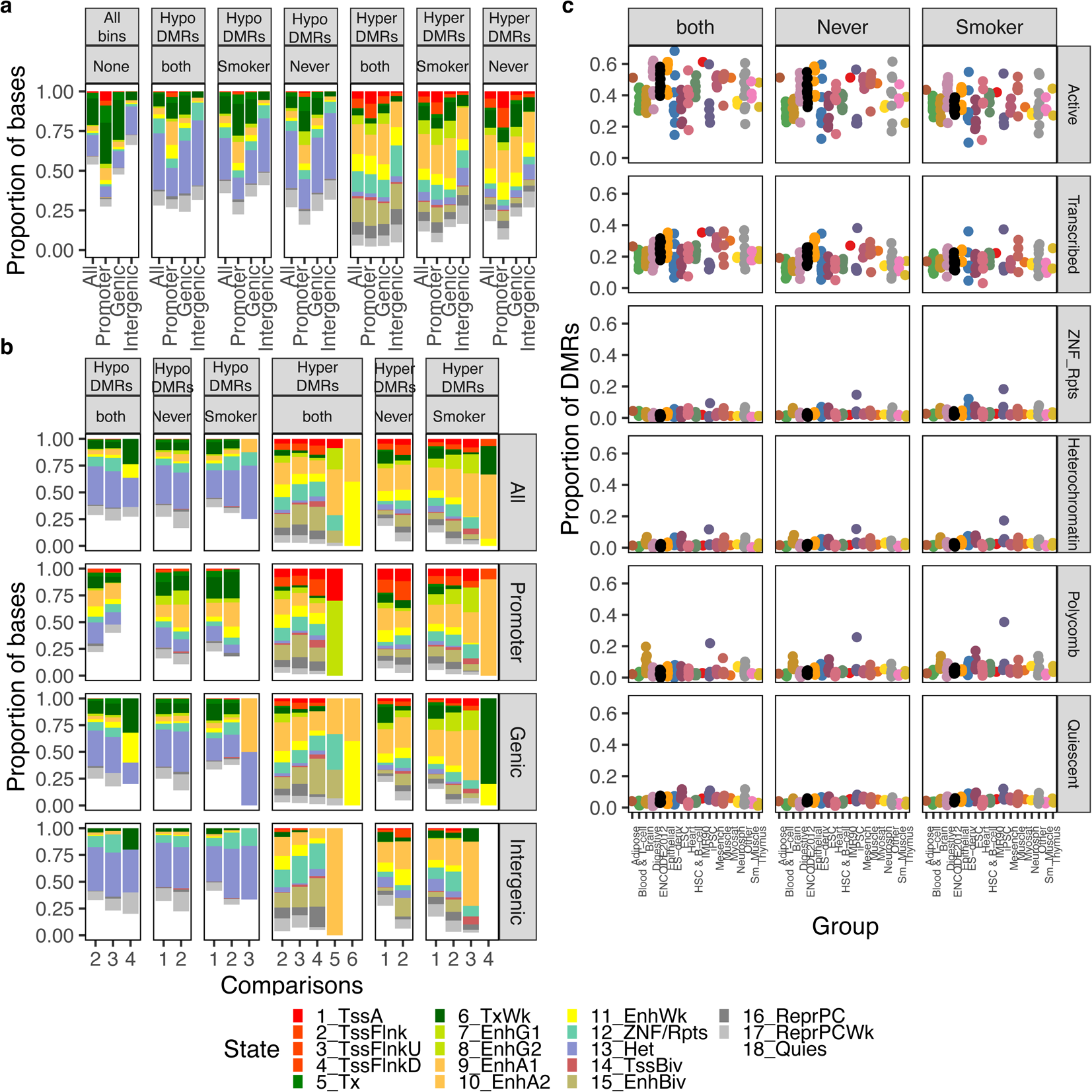
Epigenetic state of DMRs in Roadmap tissues, according to smoking status. **A.** Proportion of Roadmap sample E096 (adult lung) bases in each 18-state chromHMM state, overall and overlapping hypo- or hyperDMRs that were either exclusive to smokers, exclusive to never-smokers, or found in both, split by exclusive feature overlap. Genic DMRs overlapped genes but not promoters, and intergenic DMRs did not overlap genes or promoters. The overall state in E096 was restricted to regions overlapping 500 bp bins that contained CpGs. **B.** Proportion of Roadmap sample E096 (adult lung) bases overlapping hypo- or hyperDMRs either exclusive to smokers, never-smokers, or found in both, in each 18-state chromHMM state, split by exclusive feature overlap and the number of patient comparisons in which the DMR was found. **C.** The proportion of hypoDMRs exclusive to smokers, never-smokers, or found in both, in an active 15-state chromHMM state in each Roadmap sample, by sample group (columns) and its 18-state chromHMM state in E096 (rows) (see Materials and methods for composite state definitions). Hetero., heterochromatin.

**Supplementary Figure 20.**
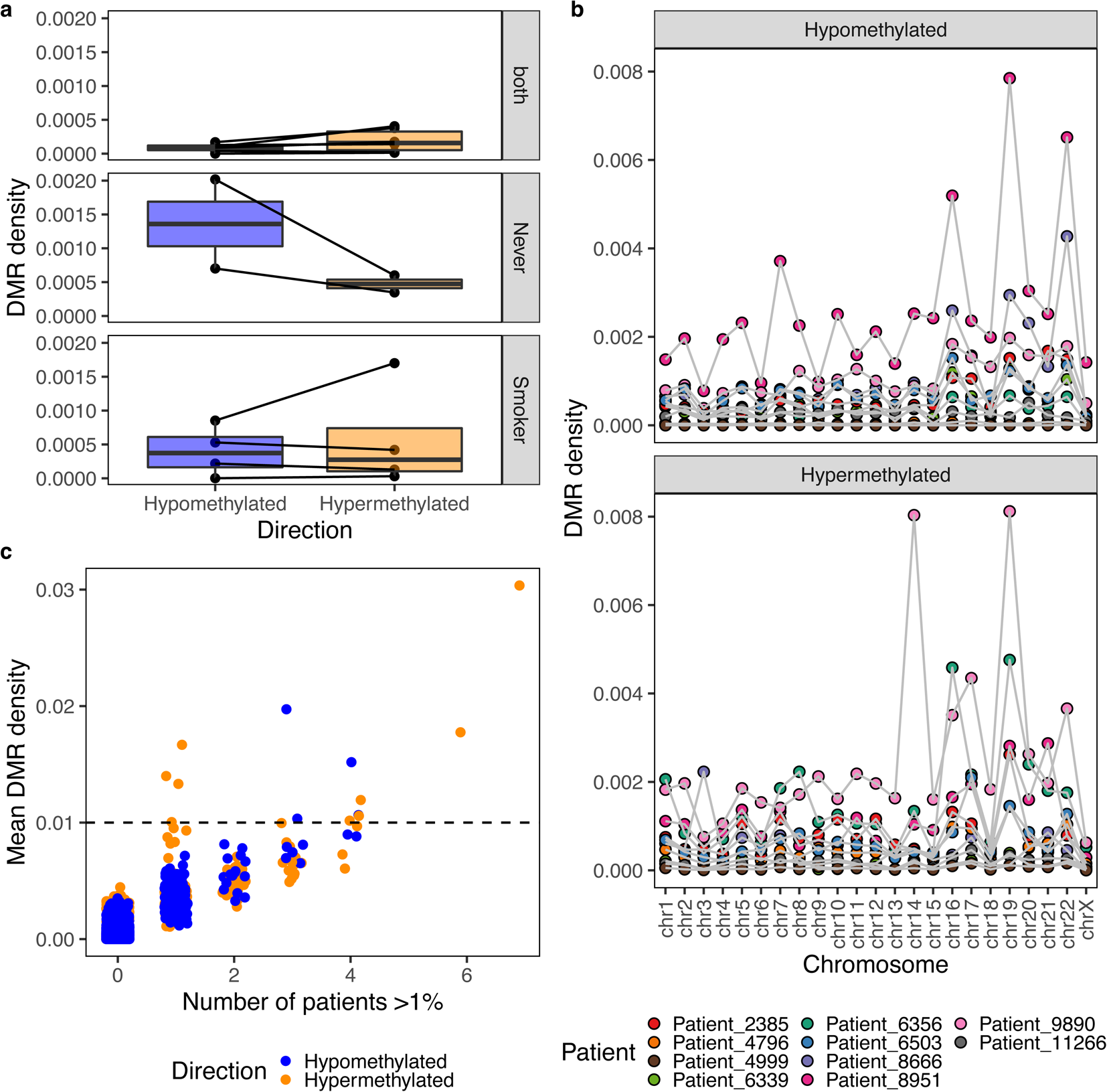
DMR density across patients and according to smoking status. **A.** Genome-wide DMR density per patient (number of DMRs versus number of 500 bp bins overlapping CpGs), by DMR direction, for DMRs exclusive to smokers (bottom), never-smokers (middle), or identified in both (top). **B.** DMR density per patient and chromosome, by DMR direction. **C.** Mean DMR density of each 1 Mb window across all patients versus the number of patients in which the DMR density was > 1%, by DMR direction. Dashed line indicates a mean DMR density of 1%. Windows that did not overlap a CpG were omitted.

**Supplementary Figure 21.**
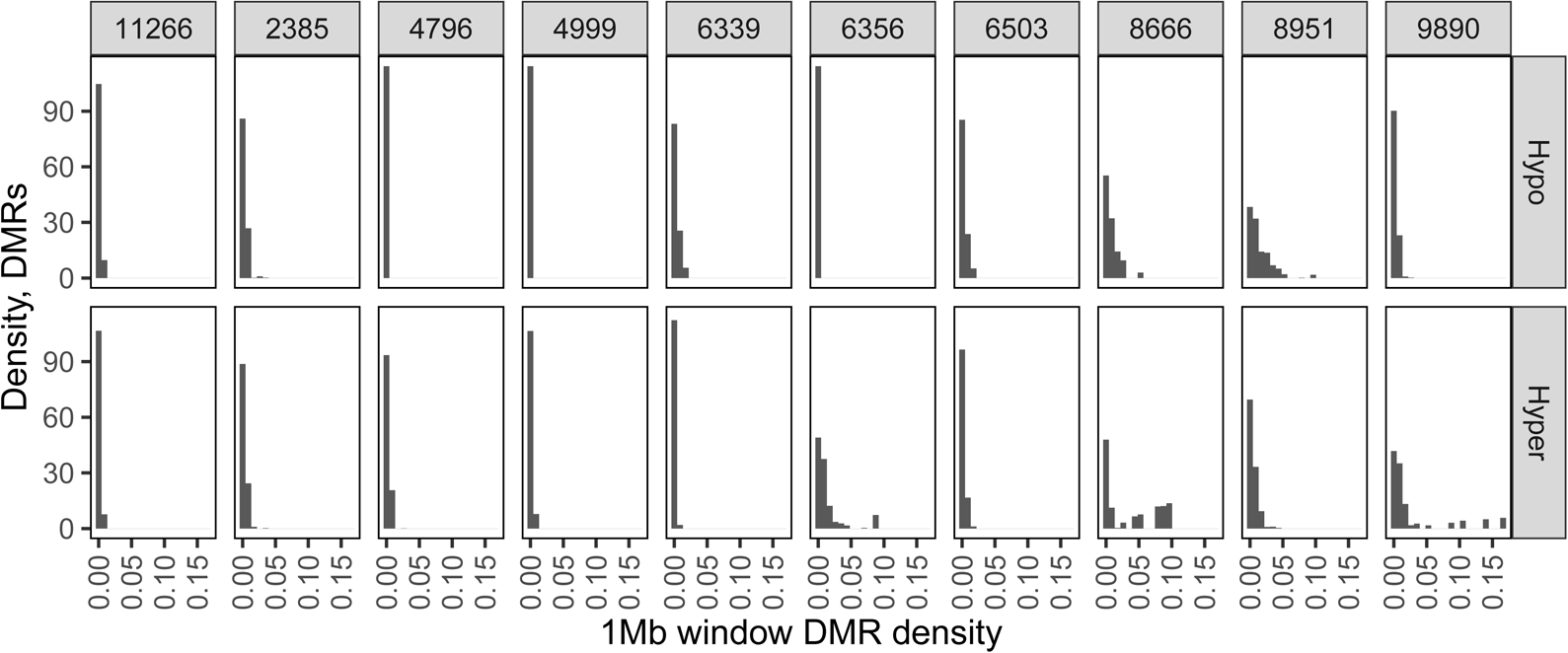
Proportion of DMRs in high-density windows. Density distribution of the proportion of DMRs exclusive to that patient that were within 1 Mb windows of each DMR density, by patient and DMR direction.

**Supplementary Figure 22.**
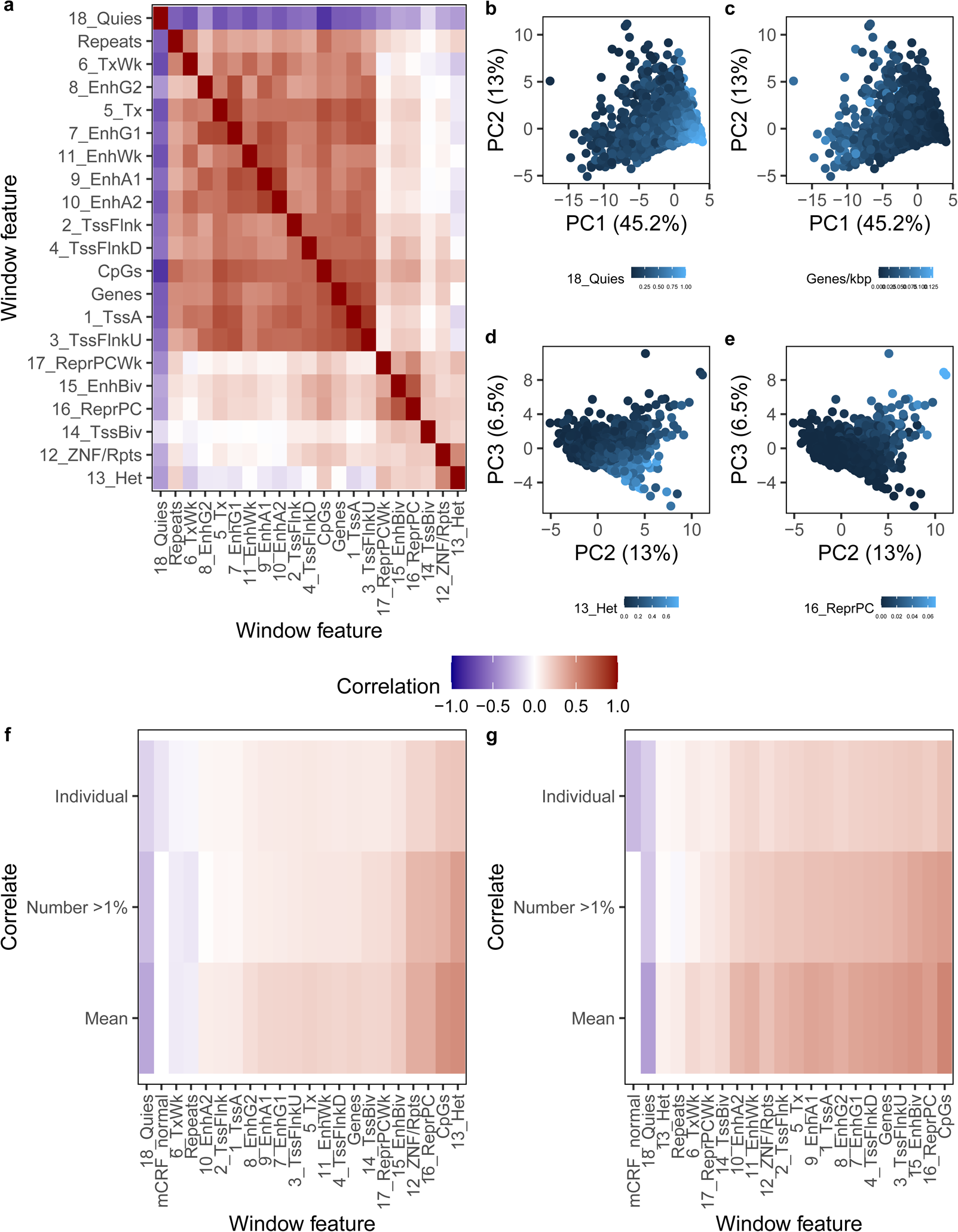
Window features correlating with DMR density. **A.** Pearson correlation between 1 Mb window features, including proportion in each 18-state chromHMM state in E096 and gene, CpG, and repeat density. Features are ordered via unsupervised hierarchical clustering with complete clustering. **B-E.** PCA on 1 Mb windows, using the above features as features. Windows are colored by **(B)** proportion in 18_Quies in E096, **(C)** gene density, **(D)** proportion in 13_Het in E096, and **(E)** proportion in 16_ReprPC in E096. Axis titles display the amount of variance explained by each PC. **F-G.** Pearson correlation between DMR density and 1 Mb window features for **(F)** hypo- and **(G)** hyperDMRs, including proportion in each 18-state chromHMM state in E096, gene, CpG, and repeat density, and average CpG methylation level in the corresponding normal sample (“mCRF_normal”). Correlations are presented for DMR density in individual comparisons (“Individual”), the number of patients in which the density was > 1% (“Number > 1%”), and the mean DMR density across patients (“Mean”). Features are ordered by median correlation. Missing values are in grey.

**Supplementary Figure 23.**
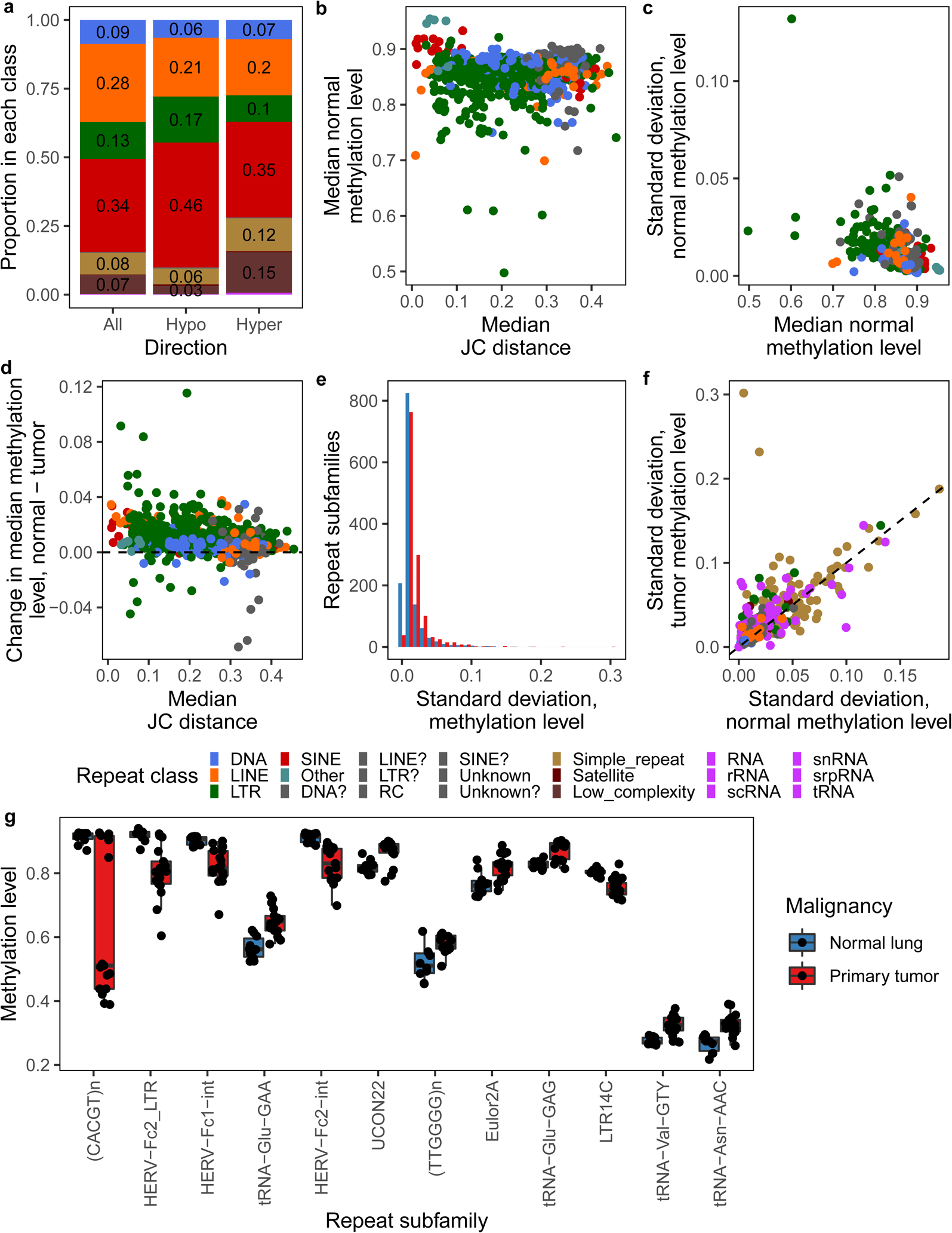
Repeat subfamily methylation level alternations in NSCLC. **A.** Proportion of repeats in each class, overall and overlapping any hypo- or hyperDMR. Proportions < 0.02 are not shown. **B.** Median subfamily Jukes-Cantor distance versus the median average CpG methylation level in normal lung samples, TE subfamilies only. **C.** Median subfamily average CpG methylation level in normal lung samples versus standard deviation across normal lung samples, TE subfamilies only. **D.** Median subfamily Jukes-Cantor distance versus the change in median average CpG methylation level between normal lung and primary tumor samples, TE subfamilies only. Dashed line indicates no change in median methylation. **E.** Distribution of standard deviations in subfamily average CpG methylation level across normal lung and tumor samples. **F.** Standard deviation in subfamily average CpG methylation level in normal lung versus tumor samples. Dashed line indicates identity. **A-D, F.** Colors legend below **F**. **G.** The average methylation level in all normal lung and tumor samples, for repeat subfamilies with a change in median average methylation between normal lung and tumors samples ≥ 0.05 and a Wilcox *p*-value < 0.01. Ordered by absolute change in median average methylation.

**Supplementary Figure 24.**
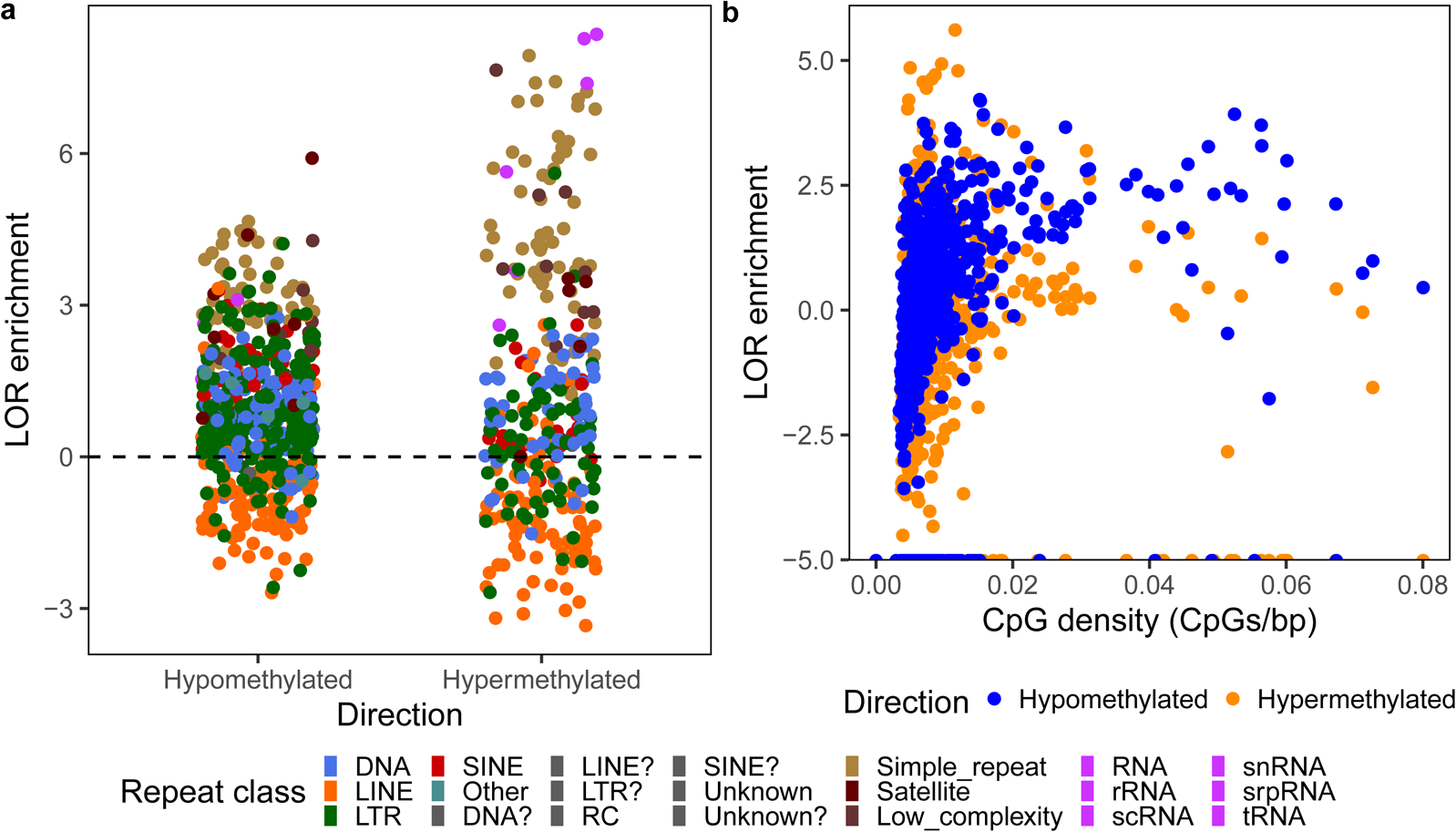
Enrichment of repeat subfamilies for overlap with DMRs. **A.** Log odds ratio enrichment of DMRs over repeat subfamilies, colored by repeat class, by DMR direction. Only subfamilies overlapping > 5 DMRs across all patients are shown. Dashed line represents no enrichment or depletion. **B.** Subfamily CpG density versus LOR enrichment for overlap with DMRs, by DMR direction, TE subfamilies only.

**Supplementary Table 1 Patient clinicopathologic data**

**Supplementary Table 2 Percent of genes in each category whose promoters overlapped DMRs**

**Supplementary Table 3 Summary of TCGA LUAD samples and files obtained by data type per case**

## Notes

**Financial support:** J.A.K is supported in part by the Siteman Cancer Center Precision Medicine Pathway (T32CA113275). E.C.P. is supported by a Postdoctoral Fellowship, PF-17-201-01, from the American Cancer Society. E.C.P., J.A.K. and T.W. are supported by NIH grants R01HG007354, R01HG007175, R01ES024992, U01CA200060, U24ES026699, U01HG009391, and American Cancer Society RSG-14-049-01-DMC.

**Conflict of interest:** The authors declare no potential conflicts of interest.

https://github.com/jaflynn5/DNA-Methylation-Changes-in-NSCLC-by-Smoking-Status

### Competing Interest Statement

The authors have declared no competing interest.

## References

1. Institute NC Cancer Stat Facts: Lung and Bronchus Cancer.

2. Nawaz K, Webster RM (2016) The non-small-cell lung cancer drug market. Nat Rev Drug Discov 15:229–230

3. Schumacher TN, Schreiber RD (2015) Neoantigens in cancer immunotherapy. Science (80-) 348:69–74

4. Herbst RS, Morgensztern D, Boshoff C (2018) The biology and management of non-small cell lung cancer. Nat Publ Gr 553:446–454

5. Govindan R, Ding L, Griffith M, et al (2012) Genomic landscape of non-small cell lung cancer in smokers and never-smokers. Cell 150:1121–1134

6. Subramanian J, Govindan R (2013) Molecular profile of lung cancer in never smokers. Eur J Cancer, Suppl 11:248–253

7. Alexandrov LB, Ju YS, Haase K, et al (2016) Mutational signatures associated with tobacco smoking in human cancer. Science (80-) 354:051417

8. Campbell JD, Alexandrov A, Kim J, et al (2016) Distinct patterns of somatic genome alterations in lung adenocarcinomas and squamous cell carcinomas. Nat Genet 48:607–616

9. Baylin SB, Jones PA (2016) Epigenetic Determinants of Cancer. Cold Spring Harb Perspect Biol 8:1–35

10. Hnisz D, Weintraub AS, Day DS, Valton A, Rasmus O (2016) Activation of proto-oncogenes by disruption of chromosome neighborhoods. https://doi.org/10.1126/science.aad9024

11. Lamprecht B, Bonifer C, Mathas S (2010) Repeat-element driven activation of proto-oncogenes in human malignancies. Cell Cycle 9:4276–4281

12. Wolff EM, Byun HM, Han HF, Sharma S, Nichols PW, Siegmund KD, Yang AS, Jones PA, Liang G (2010) Hypomethylation of a LINE-1 promoter activates an alternate transcript of the MET oncogene in bladders with cancer. PLoS Genet. https://doi.org/10.1371/journal.pgen.1000917

13. Babaian A, Mager DL (2016) Endogenous retroviral promoter exaptation in human cancer. Mob DNA 7:24

14. Brocks D, Schmidt CR, Daskalakis M, et al (2017) DNMT and HDAC inhibitors induce cryptic transcription start sites encoded in long terminal repeats. https://doi.org/10.1038/ng.3889

15. Jang HS, Shah NM, Du AY, et al (2019) Transposable elements drive widespread expression of oncogenes in human cancers. Nat Genet 51:611–617

16. Chiappinelli KB, Strissel PL, Desrichard A, et al (2015) Inhibiting DNA Methylation Causes an Interferon Response in Cancer via dsRNA Including Endogenous Retroviruses. Cell 162:974–986

17. Roulois D, Loo Yau H, Singhania R, et al (2015) DNA-Demethylating Agents Target Colorectal Cancer Cells by Inducing Viral Mimicry by Endogenous Transcripts. Cell 162:961–973

18. Wrangle J, Wang W, Koch A, et al (2013) Alterations of immune response of Non-Small Cell Lung Cancer with Azacytidine. Oncotarget 4:2067–79

19. TCGA Network, Hammerman PS, Lawrence MS, et al (2012) Comprehensive genomic characterization of squamous cell lung cancers. Nature 489:519–525

20. Collisson E a., Campbell JD, Brooks AN, et al (2014) Comprehensive molecular profiling of lung adenocarcinoma. Nature 511:543–550

21. Stevens M, Cheng JB, Li D, et al (2013) Estimating absolute methylation levels at single-CpG resolution from methylation enrichment and restriction enzyme sequencing methods. Genome Res 23:1541–1553

22. Zhang B, Zhou Y, Lin N, et al (2013) Functional DNA methylation differences between tissues, cell types, and across individuals discovered using the M&M algorithm. Genome Res 23:1522–1540

23. Karlow JA, Devarakonda S, Sankararaman S, et al Multiomic comparison of primary and brain metastatic non-small cell lung cancer suggests tumor cell reprogramming toward a glial cell phenotype.

24. Karlow JA, Devarakonda S, Xing X, Jang HS, Govindan R, Watson MA, Wang T Epigenetic reprogramming of brain development pathways during non-small cell lung cancer metastasis to brain.

25. Karlow JA, Miao B, Xing X, Wang T, Zhang B (2021) Common DNA methylation dynamics in endometriod adenocarcinoma and glioblastoma suggest universal epigenomic alterations in tumorigenesis. Commun Biol. https://doi.org/10.1038/s42003-021-02094-1

26. Weissferdt A, Moran CA (2014) Reclassification of early stage pulmonary adenocarcinoma and its consequences. J Thorac Dis 6:S581–S588

27. Yang J, Wang S, Yang Z, Hodgkinson CA, Iarikova P, Ma JZ, Payne TJ, Goldman D, Li MD (2015) The contribution of rare and common variants in 30 genes to risk nicotine dependence. Mol Psychiatry 20:1467–1478

28. Raghuwanshi SK, Nasser MW, Chen X, Strieter RM, Richardson RM (2008) Depletion of β-Arrestin-2 Promotes Tumor Growth and Angiogenesis in a Murine Model of Lung Cancer. J Immunol 180:5699– 5706

29. Furukawa C, Daigo Y, Ishikawa N, Kato T, Ito T, Tsuchiya E, Sone S, Nakamura Y (2005) Plakophilin 3 oncogene as prognostic marker and therapeutic target for lung cancer. Cancer Res 65:7102–7110

30. Chhabra D, Sharma S, Kho AT, et al (2014) Fetal lung and placental methylation is associated with in utero nicotine exposure. Epigenetics 9:1473–1484

31. Richardson TG, Richmond RC, North TL, Hemani G, Davey Smith G, Sharp GC, Relton CL (2019) An integrative approach to detect epigenetic mechanisms that putatively mediate the influence of lifestyle exposures on disease susceptibility. Int J Epidemiol 48:887–898

32. HAN Y, LI G, SU C, et al (2013) Exploratory study on the correlation between 14 lung cancer-related gene expression and specific clinical characteristics of NSCLC patients. Mol Clin Oncol 1:887–893

33. Kaczkowski B, Tanaka Y, Kawaji H, Sandelin A, Andersson R, Itoh M, Lassmann T, Hayashizaki Y, Carninci P, Forrest ARR (2016) Transcriptome analysis of recurrently deregulated genes across multiple cancers identifies new pan-cancer biomarkers. Cancer Res 76:216–226

34. Tang Z, Li J, Shen Q, et al (2017) Contribution of upregulated dipeptidyl peptidase 9 (DPP9) in promoting tumoregenicity, metastasis and the prediction of poor prognosis in non-small cell lung cancer (NSCLC). Int J Cancer 140:1620–1632

35. Fraile JM, Manchado E, Lujambio A, Quesada V, Campos-Iglesias D, Webb TR, Lowe SW, López-Otín C, Freije JMP (2017) USP39 deubiquitinase is essential for KRAS oncogene-driven cancer. J Biol Chem 292:4164–4175

36. Wang R, Deng X, Yoshioka Y, Vougiouklakis T, Park JH, Suzuki T, Dohmae N, Ueda K, Hamamoto R, Nakamura Y (2017) Effects of SMYD2-mediated EML4-ALK methylation on the signaling pathway and growth in non-small-cell lung cancer cells. Cancer Sci 108:1203–1209

37. Richter GM, Kruppa J, Munz M, et al (2019) A combined epigenome- and transcriptome-wide association study of the oral masticatory mucosa assigns CYP1B1 a central role for epithelial health in smokers. Clin Epigenetics 11:1–18

38. Richtmann S, Wilkens D, Warth A, Lasitschka F, Winter H, Christopoulos P, Herth FJF, Muley T, Meister M, Schneider MA (2019) FAM83A and FAM83B as prognostic biomarkers and potential new therapeutic targets in NSCLC. Cancers (Basel) 11:1–16

39. Dhandapani L, Yue P, Ramalingam SS, Khuri FR, Sun SY (2011) Retinoic acid enhances TRAIL-induced apoptosis in cancer cells by upregulating TRAIL receptor 1 expression. Cancer Res 71:5245– 5254

40. Milewski D, Balli D, Ustiyan V, Le T, Dienemann H, Warth A, Breuhahn K, Whitsett JA, Kalinichenko V V., Kalin T V. (2017) FOXM1 activates AGR2 and causes progression of lung adenomas into invasive mucinous adenocarcinomas. PLoS Genet 13:1–21

41. Yang X, Gao L, Zhang S (2017) Comparative pan-cancer DNA methylation analysis reveals cancer common and specific patterns. Brief Bioinform 18:761–773

42. Rauch T, Wang Z, Zhang X, Zhong X, Wu X, Lau SK, Kernstine KH, Riggs AD, Pfeifer GP (2007) Homeobox gene methylation in lung cancer studied by genome-wide analysis with a microarray-based methylated CpG island recovery assay. Proc Natl Acad Sci U S A 104:5527–5532

43. Shiraishi M, Sekiguchi A, Oates AJ, Terry MJ, Miyamoto Y (2002) HOX gene clusters are hotspots of de novo methylation in CpG islands of human lung adenocarcinomas. Oncogene 21:3659–3662

44. Zhou X, Updegraff BL, Guo Y, et al (2017) PROTOCADHERIN 7 acts through SET and PP2A to potentiate MAPK signaling by EGFR and KRAS during lung tumorigenesis. Cancer Res 77:187–197

45. Yamaguchi T, Hosono Y, Yanagisawa K, Takahashi T (2013) NKX2-1/TTF-1: An Enigmatic Oncogene that Functions as a Double-Edged Sword for Cancer Cell Survival and Progression. Cancer Cell 23:718–723

46. Zhang Y, Xu X, Zhang M, Wang X, Bai X, Li H, Kan L, Zhou Y, Niu H, He P (2016) MicroRNA-663a is downregulated in non-small cell lung cancer and inhibits proliferation and invasion by targeting JunD. BMC Cancer 16:1–10

47. Zhang C, Chen B, Jiao A, Li F, Sun N, Zhang G, Zhang J (2018) MiR-663a inhibits tumor growth and invasion by regulating TGF-β1 in hepatocellular carcinoma 11 Medical and Health Sciences 1112 Oncology and Carcinogenesis. BMC Cancer 18:1–14

48. Cho JG, Park S, Lim CH, et al (2016) ZNF224, Krüppel like zinc finger protein, induces cell growth and apoptosis-resistance by down-regulation of p21 and p53 via miR-663a. Oncotarget 7:31177–31190

49. Chang RM, Xiao S, Lei X, Yang H, Fang F, Yang LY (2017) miRNA-487a promotes proliferation and metastasis in hepatocellular carcinoma. Clin Cancer Res 23:2593–2604

50. O’Reilly D, Dienstbier M, Cowley SA, Vazquez P, Drozdz M, Taylor S, James WS, Murphy S (2013) Differentially expressed, variant U1 snRNAs regulate gene expression in human cells. Genome Res 23:281–291

51. Stadler PF, Chen JJL, Hackermüller J, et al (2009) Evolution of vault RNAs. Mol Biol Evol 26:1975– 1991

52. Pehrsson EC (2019) The epigenomic landscape of transposable elements across normal human development and anatomy. https://doi.org/10.1787/acgd-2013-23-en

53. Powrózek T, Krawczyk P, Kucharczyk T, Milanowski J (2014) Septin 9 promoter region methylation in free circulating DNA - Potential role in noninvasive diagnosis of lung cancer: Preliminary report. Med Oncol. https://doi.org/10.1007/s12032-014-0917-4

54. Shen SY, Singhania R, Fehringer G, et al (2018) Sensitive tumour detection and classification using plasma cell-free DNA methylomes. Nature. https://doi.org/10.1038/s41586-018-0703-0

55. Hulbert A, Jusue-Torres I, Stark A, et al (2017) Early detection of lung cancer using DNA promoter hypermethylation in plasma and sputum. Clin Cancer Res 23:1998–2005

56. Sur I, Taipale J (2016) The role of enhancers in cancer. Nat Rev Cancer 16:483–493

57. Lee KWK, Pausova Z (2013) Cigarette smoking and DNA methylation. Front Genet 4:1–11

58. Freeman JR, Chu S, Hsu T, Huang Y (2016) Epigenome-wide association study of smoking and DNA methylation in non-small cell lung neoplasms. Oncotarget

59. Tessema M, Yingling CM, Liu Y, Tellez CS, Van Neste L V., Baylin SS, Belinsky SA (2014) Genome-wide unmasking of epigenetically silenced genes in lung adenocarcinoma from smokers and never smokers. Carcinogenesis 35:1248–1257

60. Huang T, Chen X, Hong Q, et al (2015) Meta-analyses of gene methylation and smoking behavior in non-small cell lung cancer patients. Sci Rep 5:8897

61. Chuong EB, Elde NC, Feschotte C (2016) Regulatory activities of transposable elements: from conflicts to benefits. Nat Rev Genet. https://doi.org/10.1038/nrg.2016.139

62. Li D, Zhang B, Xing X, Wang T (2015) Combining MeDIP-seq and MRE-seq to investigate genome-wide CpG methylation. Methods 72:29–40

63. Li J, Xing X, Li D, Zhang B, Mutch DG, Hagemann IS, Wang T (2017) Whole-Genome DNA Methylation Profiling Identifies Epigenetic Signatures of Uterine Carcinosarcoma. Neoplasia 19:100–111

64. Rauch T a, Zhong X, Wu X, Wang M, Kernstine KH, Wang Z, Riggs AD, Pfeifer GP (2008) High-resolution mapping of DNA hypermethylation and hypomethylation in lung cancer. Proc Natl Acad Sci U S A 105:252–257

65. Consortium RE, Kundaje A, Meuleman W, et al (2015) Integrative analysis of 111 reference human epigenomes. Nature 518:317–330

