## Supplementary Text for "Genome-Wide Epigenomic Profiling of Primary Non-Small Cell Lung Cancer Reveals Specific and Recurrent DNA Methylation Alterations in Smoker Versus Never-Smoker Patients"

### Genome-wide methylation

There is no significant difference in the mean genome-wide CpG methylation level or the proportion of CpGs lowly- or highly methylated (< 30% or > 70%) based on sample malignancy, even when an unusually lowly methylated normal sample, P2385_N_UC, is excluded (*p* > 0.05, Wilcox test; Figure 1a, Figure S2a). However, there are still significantly more intermediately methylated CpGs in primary NSCLC than in normal lung without P2385_N_UC (Wilcox *p* < 0.001). There is no significant difference for any of the metrics between smokers and never-smokers within normal lung or tumor samples (excluding patients with unconfirmed smoking status) or based on sample malignancy within smoking status, except for intermediately methylated CpGs in smokers (Wilcox *p* < 0.05). Finally, there is no significant difference in any of the metrics among tumor samples based on tumor stage (Kruskal-Wallis test), patient sex (Wilcox test), or % tumor (Pearson correlation), nor is there any significant difference in the by-patient normal-tumor shift in any of the metrics based on clinicopathologic data, except % tumor.

There is no significant difference in the Pearson correlation between normal lung and tumor samples from the same patient based on tumor stage or subtype (Kruskal-Wallis test), smoking status (Wilcox test), or % tumor (Pearson correlation) when all patients are included.

There is not a significant difference between never-smokers and smokers in the average methylation level of CpGs over promoters, within either normal lung or primary NSCLC samples (Wilcox *p* > 0.5).

There is also no significant difference between never-smokers and smokers in either the number of filtered MRE reads or the number of sampled cut sites within normal or tumor samples (Wilcox *p* > 0.05).

### Comparison of methylCRF to previously published results and MRE-seq/MeDIP-seq data

Sixteen samples profiled with methylCRF, including normal endometrium and five types of endometrial cancer, were downloaded (1) (Figure S3a-b). Although the average methylation level for the normal endometrium is higher than for the tumor samples, the difference can be slight. Furthermore, methylation is lowest in the laser-microdissected tumor components, which likely have higher purity.

Additionally, analysis of MRE-seq data independently confirms that the primary NSCLC samples are more hypomethylated than normal lung. The number of sampled restriction enzyme cut sites is greater in tumor samples regardless of read depth, indicating that more cut sites are unmethylated in tumors (Wilcox *p* < 0.001 for sampled cut sites, *p*=0.60 for number of reads; Figure S3c).

In comparison to all possible MRE cut sites, CpGs exclusive to tumor samples (*n*=1,162,595) are enriched in intergenic regions and over repeats, while CpGs exclusive to normal lung (*n*=81,623) are enriched over genes (Figure S3d). Due to known limitations of methylCRF, the bias toward hypomethylated sites in these regions may lead to an underestimation of total genome-wide hypomethylation in primary NSCLC. As expected, both normal-exclusive and tumor-exclusive cut sites are less frequently represented than those that are shared (Figure S3e-f).

Finally, we performed PCA on MeDIP-seq and MRE-seq RPKM in 500 bp bins across the genome. MRE-seq much more clearly separates normal lung from tumor samples, with lower variability between normal samples (Figure S3g-h).

### Selection of DMR q-value threshold

The DMR *q*-value threshold was selected from among four potential thresholds: 1e-2 (M&M default threshold), 1e-3, 1e-4, and 1e-5. Although the proportion of DMRs in each *q*-value interval is similar for patient-matched normal vs. tumor DMRs, normal vs. normal DMRs, and unpaired normal vs. tumor DMRs (Figure S6a), the number of DMRs is far smaller for normal vs. normal comparisons, particularly at higher *q*-values. (Figure S6b). The exception is Patient 4999, which has a similar number of DMRs as the normal vs. normal comparisons.

For all DMRs below each of the potential *q*-value thresholds, a false positive ratio was calculated for each patient by dividing the mean number of DMRs between the normal lung sample and other normal samples by the number of DMRs in comparison to the patient-matched tumor (Figure S6c). Patient 4999 was excluded from this analysis due to its unusually low number of DMRs (Figure S6d). For some samples, the false positive ratio increases as the threshold becomes more stringent, with a greater increase below 1e-3 than between 1e-2 and 1-e3. With a *q*-value threshold of 1e-3, the greatest by-patient false positive ratio is 16% (excluding Patient 4999).

Next, for all DMRs below each potential *q*-value threshold, we analyzed the change in mean CpG methylation level over the DMR between its normal lung and tumor sample as predicted using methylCRF. With a lower *q*-value threshold, DMRs have a greater change in methylation in the expected direction (Figure S6e-f). With a *q*-value threshold of 1-e3, 88% of hypomethylated DMRs and 86% of hypermethylated DMRs have a methylation change > 10% in the expected direction. Of the hypermethylated DMRs with a methylation change < 10%, 1,036 are on chromosome 14 of Patient 9890, 739 are on chromosome 3 of Patient 8666, and 405 are on chromosome 8 of Patient 6356, which have unusually high DMR densities (Figure 5b).

Finally, we calculated the proportion of DMRs below each *q*-value threshold that are shared between patients (Figure S6g). As the threshold becomes more stringent, the proportion of DMRs that are shared decreases, but the drop is largest from 1e-2 to 1e-3.

### Number of DMRs per comparison

Neither the total number of DMRs nor the proportion hypomethylated correlates with any patient clinicopathologic data, including tumor purity (Pearson correlation), tumor stage or subtype (Kruskal-Wallis test), and smoking status (Wilcox test) (*p* > 0.05). Pearson correlations with change in average CpG methylation level between patient-matched normal and tumor samples (directional or absolute) are also not significant (*p* > 0.05).

### Normal vs. normal DMRs

There are 12,160 DMR instances between normal samples (0.7% of all instances versus 21% of all comparisons) across 4,626 unique DMRs. However, only 307 unique DMRs are exclusive to normal vs. normal comparisons (0.1%). 7,114 of the normal vs. normal DMR instances are between P2385_N_UC and another normal sample (59% versus 20% of normal vs. normal comparisons), 97% of which are hypomethylated in the other sample compared to P2385_N_UC.

The normal vs. normal comparisons with P2385_N_UC have the highest number of total DMRs, and without them, the maximum number of normal v. normal DMRs is 340.

Based on genome-wide methylation profiles, normal lung samples were more homogenous than primary NSCLC. Consistent with this observation, normal samples had a relatively uniform distribution of DMRs in comparison to all tumors and other normal lung samples, including the normal sample for the bronchioloalveolar carcinoma (Figure S7c). In contrast, the number of DMRs between each primary tumor and any normal sample varied considerably by tumor but was internally consistent (Figure S7d). This confirmed that some tumors were more epigenetically similar to normal lung, and that inter-tumor variability drove the variation observed in the number of patient-matched DMRs. In support of this observation, the number of DMRs between each tumor and its patient-matched normal sample strongly correlated with the number of DMRs between the tumor and other normal samples (Pearson correlation ≥ 0.91, *p* < 0.001, median and total number of hypo- and hypermethylated DMRs) and did not correlate with the number of DMRs between the normal and other tumor samples (*p* > 0.5 for all comparisons; Figure S7e-f).

### Feature overlap

Many DMRs overlap multiple genic features and are counted for both categories, either because 500 bp DMRs can span multiple features (as in the case of intergenic/exonic or intronic/exonic DMRs) or because feature definitions overlap (e.g., intergenic/promoter). 29% of intronic DMRs and 14% of intergenic DMRs also overlap promoters, which is higher for hypermethylated than hypomethylated DMRs. In contrast, the overlap between intergenic and intronic regions is low.

### DMR density feature correlation

The density of genes and transcript (all and protein-coding) over 1 Mb genome-wide windows is highly correlated, with a Pearson correlation > 85% for all pairwise comparisons. The correlation of each metric with DMR density over 1Mb windows has a narrow range (< 4% for both hypo- and hypermethylated DMRs, *p* < 0.001), so the metric with the highest correlation (gene density) was used for downstream analyses.

The remaining window features are highly correlated (Figure S22a). The 18_Quies state is anticorrelated with all other chromHMM states and gene, CpG, and repeat density. Active regulatory and transcribed chromHMM states are highly correlated with each other and with gene, CpG, and repeat density, with the Polycomb states and the ZNF/Rpts and heterochromatin states forming two additional clusters.

We also performed PCA on the window features to determine which contributes the most to variation among windows. PC1 (45% of variance) is highly correlated with the quiescent state (Pearson correlation 0.84, *p*=0) and anticorrelated with gene and CpG density (Pearson correlation -0.82 and -0.89, *p*=0) and other active states (Figure S22b-c). PC2 (13% of variance) separates quiescent and active states from repressed states, while PC3 (7% of variance) separates the Polycomb-repressed states from the heterochromatin and ZNF/Rpts states and repeat density (Figure S22d-e).

### Repeats

In both directions, 96% of the repeats that overlap DMRs overlap only one unique DMR. The maximum is 9 unique DMRs per repeat (a GSATX satellite element, chr12:38545411 -38551396).

### References

1.	Li J, Xing X, Li D, Zhang B, Mutch DG, Hagemann IS, et al. Whole-Genome DNA Methylation Profiling Identifies Epigenetic Signatures of Uterine Carcinosarcoma. Neoplasia. 2017;19(2):100–11.
